## Supplementary information for "Automated optimisation of solubility and conformational stability of antibodies and proteins"

#### COMPUTATIONAL METHODS

##### Benchmarking the accuracy of stability-change predictions

The accuracy of our pipeline in predicting protein stability changes upon mutation was tested on a curated subset of experimental data from the ProTherm database, which, in the context of this work, we refer to as the Frenz et al. dataset[1]. Protein thermodynamic stability changes upon mutation are expressed in terms of ΔΔG, which is the difference in the free energy difference (ΔG) between native and unfolded states of the mutational variant under scrutiny and that of the WT. The curated dataset we employed contains 755 ΔΔG values from 81 different proteins, and it was originally compiled by removing known biases in the ProTherm for the specific purpose of benchmarking software to predict the change in protein stability upon point mutation[1]. Given that from a thermodynamic point of view the definition of the wild type is ultimately arbitrary,[2] it can be assumed that the absolute value of free-energy change |ΔΔG| is the same in going from one molecule to the other, and what changes are only the signs. Therefore, for each experimentally determined ΔΔG value associated with a given mutation, we also consider the inverse variation (i.e. the mutation that reverts the variant to the original protein) by using the value of the experimental measurement with the opposite sign. This "symmetrization" procedure is commonly used in the field[2], as it balances the distribution of stabilizing and destabilizing mutations in the dataset (i.e. it yields the same number of positives and negatives in the benchmarking dataset).

In total 81 individual structures were considered, 3 of which were originally in complex with another subunit, for a total of 1510 single-point mutants. Before running the calculation of the FoldX ΔΔGs on the single-point mutants, the structures listed in the dataset were stripped down to just the chain containing the mutation under scrutiny. These structures were then repaired with the RepairPDB function of the FoldX method. The repairing process is aimed at removing potential crystal-packing artifacts and consists in correcting for incorrect rotamer assignment of all Asn, Gln and His in the protein structure, eliminating small VanderWaals' clashes through sidechain optimization, and finding new energy minima for residues with bad original energy by exploring different rotamer combinations.

All point mutations in the dataset were then applied individually to their corresponding repaired structure and their FoldX ΔΔG values were recorded. The ΔΔG values used in the benchmark were the mean values of three different FoldX runs. The same procedure was applied to the inverse mutations, by using as a starting structure the FoldX model obtained for the forward mutation (which was also then subjected to the RepairPDB command for consistency). Therefore, for the predicted ΔΔG values, it is not guaranteed that the ΔΔG of the inverse mutation is exactly minus that of the forward mutation.

##### Test of the False Discovery Rate (FDR) upon filtering with phylogenetic information

The FDR was calculated as: $FDR = \frac{FP}{FP+TP}$.

Phylogenetic information was obtained for each sequence extracted from the structures of the Frenz et al. dataset with HHblits. More specifically, the HHBLITS search was carried out with the command *hhblits -i input_file -d uniclust_database -o result_file -oa3m result_alignment -ohhm result_pssm -n 3 -diff 0 -id 95 -cov 60* on its default Uniclust30 database as downloaded from https://uniclust.mmseqs.com in November 2018 (-n 3 means three iterations, and -diff 0 means do not filter the MSAs). Different combinations were explored for the id and cov parameters, which are respectively the maximum pairwise identity and the coverage (see

**Fig. S1**). The *result_pssm* file was used as the Position Specific Scoring Matrix (PSSM) employed to apply phylogenetic filtering, and frequencies may be obtained from the integer scores in that file with $f = 2^{-score} \times 10^{-3}$ as reported in the HHblits manual.

The HHblits search was performed considering two coverage parameters (-cov 60,75) and three identity rates parameters (-id 90,95,99), which guide the filtering of homologous sequences. Moreover, we considered two versions of the PSSM: one with raw frequency counts (sometimes called Position Probability Matrix, PPM) and a log-likelihood PSSM, containing the log2 enrichment of each amino acid at each position estimated using as background frequencies the overall frequencies of each residue in the whole alignment.

We then used these PSSMs to filter the dataset to keep only those mutations with a frequency/log-likelihood higher than zero, and then to further filter it to those mutations also having a PSSM frequency/loglikelihood score higher than its WT counterpart (i.e., Δlog-likelihood or Δfrequency > 0). The FDR was then calculated on the whole datasets and on these filtered subsets, and the results are plotted in **Fig.S1**.

**Statistical significance of the FDR improvement**

To assess the statistical significance of our results we performed a random resampling test for each of the filtering conditions tested. Applying a PSSM filtering means analyzing the performance of FoldX on a subset of the database with *N* entries satisfying the filtering condition. *N* depends on the specific filtering employed, and to a lesser extent on the parameters used in the HHBLITS search, as these affect the resulting PSSM. Thus, for each filtering tested, we randomly sample *N* entries from the database (where *N* is equal to the number of entries passing the filter under scrutiny) and we assess the FDR of this random set. Repeating this procedure 100.000 times yields a normal distribution of FDR values, which enables the explicit calculation of a p-value as the probability of obtaining by random chance an FDR value as or more extreme than that obtained from the PSSM filtering under scrutiny. This approach is illustrated by an example in **Fig. S2**.

Comparing the effects that the different PSSM implementations have on the FDRs (**Fig. S1**) shows that, while simply filtering out mutations with a frequency equal to zero (i.e. never observed) doesn't significantly, or not at all, decrease the FDRs, filtering out also those mutations with log-likelihood < 0 results in a marked and significant decrease in the FDR. The FDR is then further improved in a significant way by considering only those mutations with Δ frequency/ Δ log-likelihood > 0. Finally, the lowest and most statistically significant FDRs are registered for filtering cases involving a log-likelihood PSSM obtained from an alignment with coverage 60 and identity 95, albeit these two parameters only have a very small effect on the resulting FDR.

We also verified that the improvement in the FDR observed when implementing phylogenetic filtering is not a simple consequence of removing mutations with predicted ΔΔG close to 0, which would be within the expected error of FoldX[3]. To this end, we compared the FDRs calculated at different predicted ΔΔG cutoffs values (from -4.5 to 0 kcal/mol) in three filtering conditions of **Fig. 1B** (i.e. no filters, log-likelihood > 0 and both log-likelihood and Δlog-likelihood > 0, all applied according to the PSSM obtained with coverage 60 and identity 95). In this context, the ΔΔG cutoff is the value of the FoldX predicted ΔΔG below which a mutation is considered stabilizing, with the default being 0. For each cutoff value, the FDR is calculated only on those mutations with a predicted ΔΔG smaller than the cutoff, and all other entries in the dataset are ignored. As expected, the FDR increases as the ΔΔG cutoff value approaches 0, with a seemingly linear trend from -1 onward (**Fig. S3)**. Importantly, the difference between the different filtering conditions becomes apparent before the cutoff value approaches the FoldX prediction error, which is commonly estimated at ΔΔG ~ 0.5 kcal/mol[3], indicating that the observed improvement is not a consequence of removing mutations with predicted ΔΔG close to 0. For example, a ΔΔG cutoff of - 0.74 kcal/mol would be needed to obtain the same FDR observed using the default cutoff of 0 by applying the most stringent of the phylogenetic filters (gray vertical line in **Fig. S3**).

**Conservation index calculations**

The conservation index is calculated from the raw frequency PSSM (also known as PWM), recomputed ignoring gaps, using Formula 2.2 from Ref. [4], which for position $i$ in the alignment is $C(i) = \sqrt{\sum_{a=1}^{20}[f_a(i) - f_a]^2}$ where $f_a$ is the overall alignment frequency of amino acid $a$ while $f_a(i)$ is its frequency at position $i$ under scrutiny.

**Generation of the pre-compiled alignments of immunoglobulin variable domains**

Precompiled multiple sequence alignments (MSAs) of antibody variable regions (Fv) were constructed with the program ANARCI[5] using the AHo numbering scheme[6]. We found that in some instances gaps may be opened in unexpected positions (sometimes in framework 1 or framework 2) leading to a misalignment of the subsequent part of the sequence, including the fully conserved cysteines that form the intra-domain disulphide bond (AHo positions 23 and 106). Therefore, a custom python script was run to adjust possible inaccuracies in the alignment of each sequence within the MSA. This script maximises the identity between the MSA consensus sequence and the sequence under scrutiny calculated at all positions with conservation index greater than 0.9, which include the two fully conserved cysteines. We found that misalignments were rare for VH sequences but more common for VL sequences. For example, in the post-phase-I clinical-stage antibodies MSA discussed below, the sequences with misaligned cysteines were 4 VH (0.8% of total, all fixed by the script), and 112 VL (21.5% of the total, of which 99.1% were fixed by the script). We note that this high error rate may stem from aligning kappa and lambda light chains together in a single MSA, and/or from the fact that, to the best of our knowledge, ANARCI is not widely used with the AHo numbering scheme. Sequences whose alignment could not be fixed, or that didn't have two cysteines at the conserved positions (because of e.g. sequencing errors) were discarded. Furthermore, we only kept complete Fv sequences, and VH sequences missing up to two amino acids at their C-terminus were automatically completed by adding the conserved serine at these positions.

We provide 5 different precompiled immunoglobulin-variable-domain MSAs to meet most potential user needs. All the MSA were compiled as described above, using sequences from various sources.

The first MSA is compiled from the sequences of clinical-stage antibodies that were successful at least in Phase I clinical trial (called post-phase-I). The sequences of 526 post-phase-I clinical-stage antibody therapeutics were extracted from the structural models provided within the Therapeutic Antibody Profiler[7] (opig.stats.ox.ac.uk/webapps/newsabdab/sabpred/tap), which are themselves extracted from the therapeutic antibody database (Thera-SAbDab)[8] by filtering for post-phase-I clinical candidates or approved drugs. Antibodies successful at least in Phase I clinical trial proved safe for use in humans. Given the hefty investments required to start clinical trials, it is also reasonable to assume that these antibodies all had favourable manufacturability and could be formulated to high concentrations with a good enough shelf-life. Therefore, selecting this MSA should provide maximum guarantee of only shortlisting mutations that do not affect the toxicity or developability of the antibody under scrutiny. We note that clearly, toxicity, immunogenicity, and some aspects of developability are complex properties that depend on the presence of sequence and structural patterns, rather than just on the presence or absence of individual amino acids at specific positions. Therefore, while selecting the post-phase-I MSA is the best way to minimise the occurrence of potentially problematic motifs, one should still run additional assessments on the returned designs.

The second MSA was compiled from human antibody Fv sequences obtained from a subset of the Observed Antibody Space (OAS)[9]. More specifically, the authors of the BioPhi platform[10] compiled from all human studies within the OAS a dataset comprising 100 sequences per subject and per germline family (downloaded from github.com/Merck/BioPhi-2021-publication) representing a highly diverse and representative set of the known human repertoire.

The third and fourth MSAs were compiled respectively from 1000 mouse and 1000 human VH and VL sequences corresponding to a subset of the abYsis database[11] (obtained from http://www.abysis.org/abysis/searches/search_form.cgi by selecting the relevant species in the organism tab, and then moving to the download tab to get a fasta file, which is limited to 1000 sequences).

Finally, the fifth MSA was compiled from single-domain VH antibody sequences, known to fold and function independently of a light-chain counterpart. Sequences for this MSA were downloaded from the single-domain antibody database[12] as well as from the Protein Data Bank using the Structural Antibody Database interface to filter for nanobodies[13]. Only unique sequences were retained after merging the two data sources. The final MSA comprises 1388 VHH sequences. This MSA should be used for nanobodies or other single-domain antibodies.

**Calculation of the Mutation Score**

The Mutation Score of a mutational variant is defined as a linear combination of the changes from the WT of the CamSol solubility score (ΔCamSol), of the predicted stability (FoldX ΔΔG) and in PSSM frequency (Δfrequency), as follows:

$$\text{Mutation Score} = \Delta\text{CamSol} - \text{FoldX}\,\Delta\Delta\text{G} * \begin{cases} 0.1\,, & if\ \Delta\Delta\text{G} < 0 \\ 0.2, & otherwise \end{cases} + 0.06 * \Delta\text{frequency}$$

where numerical coefficients have been empirically determined by running various examples to give a reasonable balance to the three properties, given their different absolute scales (mutations predicted to be destabilizing are weighted twice as much so that they are penalized more even if they are predicted to be highly solubilizing). This empirical score is simply used to enable a simple ranking of different mutations with respect to how beneficial they are in improving the stability and solubility of the WT protein. While the specific values of these coefficients are in no way particularly optimal, the strength of the method is in selecting only mutations that benefit (or leave unaltered) both solubility and stability by requiring both ΔCamSol > 0 and FoldX ΔΔG < 0 (plus the chosen filtering on the PSSM frequencies). It should be noted that the option to consider mutations with FoldX ΔΔG > 0 is provided in the score above to handle those very rare cases where no mutation is found that have a beneficial effect on both stability and solubility, but this is not part of the standard implementation uploaded to the webserver, where only mutations with FoldX ΔΔG < 0 are considered. The Mutation Score above is simply used to provide a convenient ranking of all beneficial mutations, which facilitates the implementation of the combination of mutations. What ultimately matters for the final output is not the Mutation Score per se, but rather the predicted contributions to solubility and stability, which are provided separately to the user.

**Contact map definition**

In the combination process of the algorithm, a contact map is used to flag mutant combinations as "potentially synergistic". Here, two candidate mutation sites are defined as in contact if any of their non-hydrogen sidechain atoms are found at a distance smaller than 3Å.

**Best mutant combination groups identification: first derivative and weighted analysis**

Towards the end of the pipeline, the algorithm identifies the best combination groups (a group contains all mutational variants with the same number of mutations), defined as those that mark a change in the increment of the mutation score from one recursive combination step to the next. To detect these points, we combine two types of analysis (**Fig. S4**). Initially, we calculate the first derivative of the function defined by the maximum Mutation Score for each group as a function of the number of mutations in that group (group of single-point mutants, double mutants, three, four, and so on). Here, the absolute maximum of the first derivative (i.e. the number of mutations at which the steepest increase is observed), and any other point that is at least 10% higher than the point that precedes it, are considered as a best mutant combination group. Then, we perform what we call a "weighted analysis", which is carried out in the cartesian space (x-axis: number of mutations; y-axis: Mutation Score), and, for each group, it consists in measuring the distance between the point (number of mutations in group, maximum Mutation Score of the group) and an auxiliary point with coordinates (0, maximum Mutation Score observed among all groups). This auxiliary point can be regarded as the maximum achievable improvement (i.e., the maximum observed mutation score) with the minimum number of mutations (i.e., zero). The measurement of the distance is repeated iteratively multiple times, and each time a weight of 0.5 is added to the y coordinate of the auxiliary point. At each iteration, the group that is closest to the moving auxiliary point is recorded, and at the end of the analysis, we obtain an array of counts telling us how many times each group has been registered as the nearest to the moving auxiliary point. Those groups corresponding to counts higher than the count obtained for the groups with N+1 mutations are labelled as the best mutant combinations. The two analyses give as output two lists of integers indicating the number of mutations within the identified best mutation combination group (e.g. [1,4,7]). These lists are then merged keeping only unique elements, which correspond to the best combination groups for which the algorithm returns multiple designs (one design is returned anyway for each combination group).

**Automated design of adalimumab, golimumab, CR3022, H11-H4 and H11-D4**

The inputs to the automated computational design pipeline were chosen to be the biological assemblies from the PDB database, which consist in the antibody-antigen complex. Specifically, PDB ID, heavy, light, and antigen chain IDs were respectively 3wd5, H, L, and A for adalimumab, which was run twice once by selecting the post-phase-I MSA and once the OAS-human MSA; 5yoy, H, E, and C for golimumab, which was run with the OAS-human MSA; 6w41, H, L, and C for CR3022, for which we also ran the alternative structure 7jn5, H, L, and F. Both CR3022 runs were performed using the post-phase-I MSA. For the nanobodies PDB ID, nanobody, and antigen chain IDs were respectively 6yz5, F, and E for H11-D4, and 6zbp, B, and A for H11-H4, both of which were run with the single-domain-antibody MSA. We limited the design to 5 and 3 maximum number of mutations for scFvs and nanobodies respectively. We selected a total of 26 variants for experimental validation, comprising the 5 WTs and 21 mutational variants. These designs and the main reasons for selecting them are listed in **Table S3**, to access full details on the automated calculations see Supplementary Files 3 to 9.

***Antigen-contact analysis and flagged mutations.*** Given that most of the suggested mutations are in CDR regions, and that paratope positions were not explicitly excluded from the design, we carried out an additional check on the returned mutations. Specifically, we used the UCSF Chimera program [14] to visually inspect the mutation sites in the context of the antigen-bound WT structure. The tools “find hydrogen bonds” and “find contacts” were employed with default options to identify those mutation sites likely to make important interactions with the antigen, and that therefore may impact affinity if mutated. Once run for each mutation site in each antibody, this analysis flagged L106 in the nanobody H11-H4, TH52 in adalimumab (Threonine, heavy chain, pdb resnumber 52), YL30 in golimumab, and no mutation site in H11-D4 and CR3022. Shortlisted mutations at these 3 flagged sites were all predicted to have a negative (stabilising) $\Delta\Delta G$ by FoldX. However, for these sites the calculated $\Delta\Delta G$ will have a contribution coming from interactions with the antigen and one coming from interactions with the antibody itself, and these two contributions may be conflicting. This fact is most apparent for site Y30.L in golimumab, where the mutation YL30G removes a hydrogen bond with the antigen. The calculated $\Delta\Delta G$ of YL30G is negative but so small that zero is within its computed uncertainty. Nevertheless, this mutation is selected because it is singly predicted to be the best at improving the CamSol solubility score and has the highest $\Delta$log-likelihood (see Supplementary file 5). Similarly, L106P in H11-H4 makes several contacts with the antigen, and a substitution to a proline is likely to affect the CDR3 loop conformation and thus severely impact binding. Conversely, the mutation TH52S in adalimumab is predicted to make an extra hydrogen bond with the antigen, and therefore may potentially improve affinity or at least not severely compromise it. Finally, we noted that the mutation WH53P in adalimumab, while not in direct contact with the antigen according to the UCSF Chimera default parameters, is introducing a proline in a binding loop and therefore may also severely impact affinity by changing the loop conformation (for instance the adjacent position 52 is in direct contact with the antigen). This mutation was not flagged as it isn’t antigen contacting according to the definition used in this analysis.

#### EXPERIMENTAL METHODS

##### Antibody production and purification

The antibodies were expressed with a C-terminal linker and 6X His tag using a pTT5 expression vector, the plasmids were purchased from Twist Bioscience. Expi293F cells from Gibco were transiently transfected according to the manufacturer's protocol at a final volume of 100 ml for the Nb.B201 variants and 60 mL for all other variants. They were harvested on day 5 by centrifugation and filtration. Purification was performed on an Äkta Xpress system with a two-step purification process, using an automated multistep procedure. First, using a HisTrap Excel 1 mL affinity chromatography (AC) column and subsequently a Superdex 75 size exclusion chromatography (SEC) column. Before purification, the columns were equilibrated in a buffer comprising of 20 mM Hepes and 150 mM NaCl, pH 7.4. The samples were loaded at a flow rate of 0.5 mL/min onto the AC column, on which it was washed with 5 CV equilibration buffer, then 15 CV wash buffer comprising of 20 mM Hepes, 1000 mM NaCl and 30 mM Imidazole, pH 7.5, and lastly with 15 CV equilibration buffer. The antibodies were then eluted into a collecting loop using 20 mM Hepes, 150 mM NaCl and 500 mM Imidazole, pH 7.4, and run directly onto the SEC column at a flow rate of 1 mL/min. The peaks were monitored using UV280 and fractionated into a 96-well deep well collector plate.

##### Liquid chromatography–mass spectrometry

The mass of all antibodies was verified by LC-MS on an Agilent Technologies 6224 (ESI-TOF LC-MS) using a Waters MassPREP desalting column. 10 ul of sample was injected with a flow rate of 0.4 mL/min and the analysis was carried out at RT. Buffer A comprised of 0.1% formic acid in water and buffer B of 0.1% formic acid in acetonitrile. The chromatographic program was running as follows: 0-3 min of 5% buffer B, 3-4 min of a gradient 5 to 60% of buffer B, 4-6 min of 60% buffer B, 6-7 min of 60-95% buffer B, 7-8 min of 95% buffer B followed by four alternating 1 min washes of 95% and 5% buffer B to regenerate the. The MS data settings were set to 4 GHz at high resolution in a 100 to 3200 m/z mass range using dual ESI at positive mode and the spraying conditions were set to 325°C, using a N2 flow rate of 13 L/min, and the voltage set to 300V. The masses were deconvoluted using the AgilentMassHunter Qualitative Analysis B.05.00 program. A reduced mass of about 18 Da, consistent with pyroglutamination (-17) and the presence of a disulphide bond (-2 Da) was detected in the variants containing a N-terminal glutamine.

##### Size-Exclusion Chromatography

Analytical size-exclusion high performance liquid chromatography (SE-HPLC) was applied to evaluate the purity of the antibodies. 5 ul sample was loaded on an Agilent 1200 Series HPLC system and run on a BioSep SEC-s3000 column. They were run at a flow rate of 1.5 ml/min at RT using a running buffer comprising of 122 mM Disodium phosphate, 78 mM Sodium phosphate, 300 mM Sodium chloride and 10% Isopropanol, pH 6.9, with a runtime of 14 min. The signal was monitored using fluorescence detection at 354 nm.

##### Circular dichroism (CD)

CD was performed on a Chirascan Applied Photophysics spectropolarimeter using a 0.1 cm pathlength quartz cuvette. The antibodies were measured at a concentration of 0.1 mg/ml in a buffer containing 20 mM sodium phosphate and 100 mM sodium chloride, pH 8. The spectra were recorded from 200 to 250 nm at 25 °C and five accumulations were taken for each sample.

##### Biolayer interferometry

The affinities of the variants were evaluated using BLI. The Octet HTX system was employed running 16-sensors in parallel using 96 well half area plates with a volume of 100 ul for all wells. Streptavidin (SA) sensors were pre-hydrated in Assay buffer containing 20 mM HEPES, 150 mM NaCl and 0,05% Tween20, pH 7.4.

The loading wells contained 5 ug/ml HSA, RBD or TNF-α, which were biotinylated using NHS chemistry. The association wells contained the antibodies at concentrations of 1500, 500 and 167 nM for Nb.b201 variants, 90, 30 and 10 nM for CR3022 and H11 variants and 9, 3 and 1 nM for Humira and Golimumab variants. All variants and antigens were diluted in Assay buffer. The assay was run at 30 ºC with a shaking speed of 1000rpm. First, the sensors were moved into the Assay buffer for a baseline step, followed by loading of the respective antigens, then a second baseline step in the Assay buffer, followed by either 180 s association and dissociation for the Nb.b201 variants or 300 s association and dissociation for the rest of the variants. The affinities of the antibodies were evaluated in the Data analysis HT 12.0 software from FORTÉBIO, where the data was fitted to a 1:1 binding model using unlinked Rmax values.

**Nano differential scanning fluorimetry**

NanoDSF was run using a Prometheus NT.48. Standard capillaries were employed, and the samples were run at 30 uM for the Nb.b201 variants and 0,1 mg/ml for the rest. All variants were run in duplicates. A discovery scan was first performed, and the excitation power was adjusted accordingly to reach a fluorescent count between 1000 and 15000. The antibodies were unfolded by applying a 1°C/min temperature ramp going from 20°C to 95°C. The NanoTempers PR.Control software was used to analyse the antibodies unfolding as the peak of the first derivative of the 350 nm emission signal.

**Cross-Interaction Chromatography (CIC)**

Evaluation of the nonspecific interaction between the variants and IgGs from human serum (Sigma, I4506) was performed by CIC. The column was prepared following the description in Supplementary Ref. [15]. 10 mg of IgG was immobilized to a HiTrap NHS-activated HP column (column volume 1 mL) on an Äkta pure system (Cytiva, Sweden). and run on an Agilent Infinity HPLC system. 5 uL sample at 1 mg/mL for the Nb.b201 variants, and 0,1 mg/ml for all other variants, was injected in duplicates with a running buffer comprising of 20 mM sodium phosphate and 140 mM sodium chloride, pH 7.4. The flow rate was 0.1 mL/min and the runtime was 30 min. UV absorbance was monitored at 280 nm.

**PEG and AMS precipitation**

PEG and AMS precipitations were carried out following closely the protocol described in detail in Supplementary Ref. [16]. The buffer was 10 mM Citrate 10 mM Phosphate buffer at pH 7.4. In the case of AMS, a stock of 3.8 M dissolved in the same buffer (and with re-adjusted pH) was used.

| Antibody fragment | Purified sequence | PDB ID & Chain IDs |
|---|---|---|
| Nb.b201 | QVQLQESGGGLVQAGGSLRLSCAASGYISDAYYMGWYRQAPGK<br>EREFVATITHGTNTYYADSVKGRFTISRDNAKNTVYLQMNSLK<br>PEDTAVYYCAVLETRSYSFRYWGQGTQVTVSS<br>GGGSHHHHHH | 5vnw chain C |
| Nb H11-D4 | QVQLVESGGGLMQAGGSLRLSCAVSGRTFSTAAMGWFRQAPGK<br>EREFVAAIRWSGGSAYYADSVKGRFTISRDKAKNTVYLQMNSL<br>KYEDTAVYYCARTENVRSLLSDYATWPYDYWGQGTQVTVSS<br>KHHHHHH | 6yz5 chain F |
| Nb H11-H4 | QVQLVESGGGLMQAGGSLRLSCAVSGRTFSTAAMGWFRQAPGK<br>EREFVAAIRWSGGSAYYADSVKGRFTISRDKAKNTVYLQMNSL<br>KYEDTAVYYCAQTHYVSYLLSDYATWPYDYWGQGTQVTVSS<br>KHHHHHH | 6zbp chain B |
| Adalimumab scFv | EVQLVESGGGLVQPGRSLRLSCAASGFTFDDYAMHWVRQAPGK<br>GLEWVSAITWNSGHIDYADSVEGRFTISRDNAKNSLYLDMNSL<br>RAEDTAVYYCAKVSYLSTASSLDYWGQGTLVTVSSSGGGGSGG<br>GGSGGGGSGGGGSDIQMTQSPSSLSASVGDRVTITCRASQGIR<br>NYLAWYQQKPGKAPKLLIYAASTLQSGVPSRFSGSGSGTDFTL<br>TISSLQPEDVATYYCQRYNRAPYTFGQGTKVEIKR<br>GGGSHHHHHH | 3wd5 chains H & L |
| Golimumab scFv | EIVLTQSPATLSLSPGERATLSCRASQSVYSYLAWYQQKPGQA<br>PRLLIYDASNRATGIPARFSGSGSGTDFTLTISSLEPEDFAVY<br>YCQQRSNWPPFTFGPGTKVDIKTSGGGGSGGGGSGGGGSGGGG<br>SQVQLVESGGGVVQPGRSLRLSCAASGFIFSSYAMHWVRQAPG<br>NGLEWVAFMSYDGSNKKYADSVKGRFTISRDNSKNTLYLQMNS<br>LRAEDTAVYYCARDRGIAAGGNYYYYGMDVWGQGTTVTVSS<br>GGGSHHHHHH | 5yoy chains H & E |
| CR3022 scFv | QMQLVQSGTEVKKPGESLKISCKGSGYGFITYWIGWVRQMPGK<br>GLEWMGIIYPGDSETRYSPSFQGQVTISADKSINTAYLQWSSL<br>KASDTAIYYCAGGSGISTPMDVWGQGTTVTVSSSGGGGSGGGG<br>SGGGGSGGGGSDIQLTQSPDSLAVSLGERATINCKSSQSVLYS<br>SINKNYLAWYQQKPGQPPKLLIYWASTRESGVPDRFSGSGSGT<br>DFTLTISSLQAEDVAVYYCQQYYSTPYTFGQGTKVEIKR<br>GGGSHHHHHH | 6w41 chains H & L<br>(also 7jn5 chains H & L, see Table S3) |

**Table S1.** The full sequences of the WT nanobodies (Nb) and single-chain antibodies (scFv) used in experiments. All sequences end with a poly-His purification tag. The N-terminal Q is typically pyroglutamine in the expression system employed. The residue position used for mutations in the text, figures, and other tables is the PDB residue numbering (last column).

| Variant | LC-MS mass (Da) | Theoretical Mass (Da) | ΔMass (Da) | SE-HPLC purity (%) |
|---|---|---|---|---|
| Nb.b201 WT | 14158.17 | 14176.45 | -18.28 | 98.33 |
| Nb.b201 A31D | 14202.93 | 14220.46 | -17.53 | 99.72 |
| Nb.b201 H53P | 14118.19 | 14136.42 | -18.23 | 99.85 |
| Nb.b201 H53R | 14177.45 | 14195.49 | -18.04 | 99.92 |
| Nb.b201 T55D | 14172.44 | 14190.43 | -17.99 | 99.78 |
| Nb.b201 T55G | 14115.04 | 14132.39 | -17.35 | 99.68 |
| Nb.b201 T100K | 14185.07 | 14203.52 | -18.45 | 99.82 |
| Nb.b201 T100R | 14213.46 | 14231.53 | -18.07 | 99.85 |
| Nb.b201 A31D T55D | 14216.5 | 14234.44 | -17.94 | 99.76 |
| Nb.b201 A31D T55G | 14158.6 | 14176.4 | -17.8 | 99.72 |
| Nb.b201 H53P T55G | 14074.91 | 14092.37 | -17.46 | 99.87 |
| Nb.b201 A31D H53P T55G | 14119.1 | 14136.38 | -17.28 | 99.77 |
| Nb.b201 A31D H53P T55G T100K | 14146.57 | 14163.45 | -16.88 | 99.63 |
| Nb.b201 Y33A Y58A Y103A F105A | 13805.83 | 13824.06 | -18.23 | 97.78 |
| Adalimumab WT | 27455.75 | 27458.92 | -3.17 | 100.00 |
| Adalimumab AH23K AH40P SH49G WH53P AL94P | 27445.83 | 27448.96 | -3.13 | 100.00 |
| Adalimumab AH23K TH52S WH53P SH55G AL94P | 27405.83 | 27408.90 | -3.07 | 100.00 |
| Adalimumab AL94P | 27481.80 | 27484.95 | -3.15 | 100.00 |
| Adalimumab SH55G | 27425.84 | 27428.89 | -3.05 | 100.00 |
| Adalimumab WH53P | 27366.76 | 27369.82 | -3.06 | 100.00 |
| Adalimumab AH23K AH40P SH49G | 27508.16 | 27512.02 | -3.86 | 98.33 |
| Adalimumab AH23K AH40P SH49G SH55G | 27477.98 | 27482.00 | -4.02 | 96.30 |
| Adalimumab AH23K AH40P SH49G TH52S SH55G | 27463.83 | 27467.97 | -4.14 | 94.01 |
| Adalimumab AH23K AH40P SH55G | 27508.11 | 27512.02 | -3.91 | 84.30 |
| Adalimumab AH23K SH49G SH55G | 27452.37 | 27455.96 | -3.59 | 92.12 |
| Adalimumab AH23K SH49G TH52S SH55G | 27438.09 | 27441.93 | -3.84 | 93.85 |
| Adalimumab AH40P SH49G SH55G | 27420.79 | 27424.90 | -4.11 | 87.61 |
| Adalimumab AH40P SH49G TH52S SH55G | 27407.33 | 27410.87 | -3.54 | 88.80 |
| CR3022 MH40P | 27912.45 | 27932.54 | -20.09 | 100.00 |
| CR3022 MH40P AL86P | 27938.45 | 27958.58 | -20.13 | 100.00 |
| CR3022 TH9P MH40P QH67R TL59K AL86P | 27989.80 | 28009.72 | -19.92 | 100.00 |
| CR3022 TH9P MH40P SH89E SH56G | 27920.44 | 27940.56 | -20.12 | 99.90 |
| CR3022 TH9P MH40P TL59K AL86P | 27961.58 | 27981.66 | -20.08 | 100.00 |
| CR3022 WT | 27946.52 | 27966.62 | -20.10 | 100.00 |
| Golimumab AH23K AH40P SH52D TH121L | 28286.13 | 28288.93 | -2.80 | 100.00 |
| Golimumab AH23K AH40P SH52D TH121L YL30G | 28179.82 | 28182.80 | -2.98 | 100.00 |
| Golimumab AH40P | 28188.92 | 28191.77 | -2.85 | 97.07 |
| Golimumab SH52D | 28190.96 | 28193.74 | -2.78 | 99.91 |
| Golimumab WT | 28162.84 | 28165.73 | -2.89 | 98.46 |
| H11-D4 | 14976.28 | 14989.56 | -13.28 | 100.00 |
| H11-D4 F88P T111H W112I | 14868.48 | 14886.48 | -18.00 | 100.00 |
| H11-D4 T111H | 15007.53 | 15025.59 | -18.06 | 100.00 |
| H11-D4 Y88P T111H | 14941.44 | 14959.54 | -18.10 | 100.00 |
| H11-D4 Y88P W112I | 14832.64 | 14850.45 | -17.81 | 99.93 |
| H11-H4 | 15012.18 | 15025.58 | -13.40 | 100.00 |
| H11-H4 T111H | 15043.53 | 15061.62 | -18.09 | 100.00 |
| H11-H4 Y88P | 14941.72 | 14959.53 | -17.81 | 100.00 |
| H11-H4 Y88P L106P T111H | 14961.43 | 14979.52 | -18.09 | 100.00 |
| H11-H4 Y88P T111H | 14977.56 | 14995.56 | -18.00 | 100.00 |

**Table S2. LC-MS mass and SE-HPLC purity of the antibody variants used in this study.** A difference in mass (Δmass) of -17 Da indicates that pyroglutamine is present at the N-terminus instead of glutamine. The theoretical mass is calculated without considering disulphide bonds (-2 Da difference each).

| Antibody | Mutation(s) | Notes on design and reasons for selection |
|---|---|---|
| Adalimumab | WT | |
| | WH53P | 1st single mutation, post-phase-I MSA |
| | AL94P | 2nd single mutation, post-phase-I MSA |
| | SH55G | 3rd single mutation, post-phase-I MSA |
| | AH23K;TH52S†;WH53P;SH55G;AL94P | Top 5-mut. design, post-phase-I MSA |
| | AH23K;AH40P;SH49G;WH53P;AL94P | Top 5-mut. design, human MSA |
| | AH23K;AH40P;SH49G<br>AH23K;SH49G;SH55G<br>AH40P;SH49G;SH55G<br>AH23K;AH40P;SH55G | 2nd round designs, all possible triple combinations of the returned mutations (from designs above) excluding flagged TH52S, false positive AL94, and affinity hampering WH53P. |
| | AH23K;AH40P;SH49G;SH55G<br>AH23K;SH49G;TH52S;SH55G<br>AH40P;SH49G;TH52S;SH55G<br>AH23K;AH40P;SH49G;TH52S;SH55G | 2nd round designs, with all returned mutations except false positive AL94, and affinity hampering WH53P |
| Golimumab | WT | |
| | AH40P | 1st single mutation |
| | SH52D | 2nd single mutation |
| | AH23K;AH40P;SH52D;TH121L | Top 4-mut. design, without flagged mutation YL30G |
| | AH23K;AH40P;SH52D;TH121L;YL30G† | Top 5-mut. design |
| CR3022 | WT | |
| | MH40P | 1st single mutation |
| | MH40P;AL86P | 1st double mutant |
| | TH9P;MH40P;SH89E;SH56G | 4-mut design composed from the run on the alternative structure PDB 7jn5. Here MH40P and SH56G have respectively 1st and 2nd best ΔΔG, while TH9P and SH89E 1st and 2nd best Δlog-likelihood. |
| | TH9P;MH40P;TL59K;AL86P | Top 5-mut. design without QH67R, as based on the Nb.B201 results we suspected that a substitution to R may worsen CIC performance (even if position 67 is not in the CDRs). |
| | TH9P;MH40P;QH67R;TL59K;AL86P | Top 5-mut. design |
| H11-H4 | WT | |
| | Y88P | 1st single mutation |
| | T111H | 3rd single mutation (2nd is flagged) |
| | Y88P;T111H | Top 2-mut. design, without flagged mutation L106P |
| | Y88P;L106P†;T111H | Top 3-mut. design |
| H11-D4 | WT | |
| | T111H | 3rd single mutation for comparison with H11-H4 |
| | Y88P;T111H | 2-mut. design for comparison with H11-H4 |
| | Y88P;W112I | Top 2-mut. design |
| | Y88P;T111H;W112I | Top 3-mut. design |

**Table S3. Nanobody and scFv variant produced in the second stage of experimental validation.** The dagger symbol (†) denotes mutations flagged by the antigen-contact analysis. The last column lists the main reason for selecting each design. To access all the details of each design calculation, see **Supplementary Files 3 to 9**.

**Supplementary file 1.** Included as a separate pdf file. Final report from the webserver from a run on bacillus licheniformis alpha-amylase (PDB ID 1bli). The report is produced as a html page by the webserver. Page breaks in this and all other supplementary files below resulted from the conversion to pdf, and may therefore have non-ideal formatting.

**Supplementary file 2.** Included as a separate pdf file. Final report from the webserver from a run on nanobody Nb.b201 (chain C of PDB ID 5vnw).

**Supplementary file 3.** Included as a separate pdf file. Final report from the webserver from a run on adalimumab Fab using the post-phase-I MSA with a limit of 5 simultaneous mutations.

**Supplementary file 4.** Included as a separate pdf file. Final report from the webserver from a run on adalimumab Fab using the OAS human MSA with a limit of 5 simultaneous mutations.

**Supplementary file 5.** Included as a separate pdf file. Final report from the webserver from a run on golimumab scFv using the OAS human MSA with a limit of 5 simultaneous mutations.

**Supplementary file 6.** Included as a separate pdf file. Final report from the webserver from a run on CR3022 Fab using the post-phase-I MSA with a limit of 5 simultaneous mutations. PDB ID 6w41 was used as input.

**Supplementary file 7.** Included as a separate pdf file. Final report from the webserver from a run on CR3022 Fab using the post-phase-I MSA with a limit of 5 simultaneous mutations. PDB ID 7jn5 was used as input. 6w41 differs from 7jn5 in the VH domain, which ends respectively with TVSS and TVVS followed by a different linker to the constant domain.

**Supplementary file 8.** Included as a separate pdf file. Final report from the webserver from a run on nanobody H11-H4.

**Supplementary file 9.** Included as a separate pdf file. Final report from the webserver from a run on nanobody H11-D4.

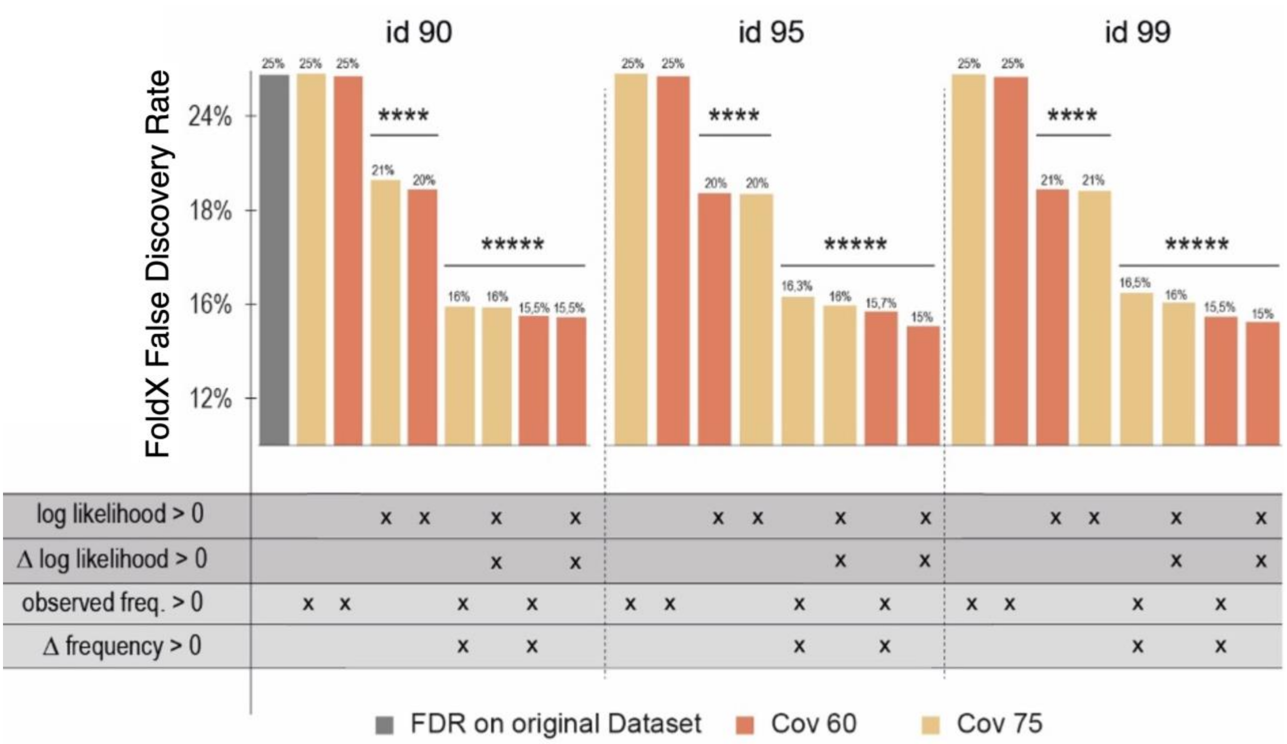


**Figure S1.** Summary of statistical test results for all three identity rates (id) used in the alignment. The originally calculated FDR for the Frenz et al dataset is represented by the grey bar and plotted against the FDRs obtained in each filtering case. The statistically significant results are marked by asterisks, which in turn define the magnitude of the corresponding p-value. Below the graph, a table shows which filters have been applied for obtaining every FDR. The rows describe the different filtering conditions and the “x” signs mark which have been used in each testing case. Highlighted in yellow are the FDRs obtained with an alignment with coverage (Cov) 75 and in red are the ones obtained with coverage 60. Above each bar is written the precise FDR calculated for the respective filtering case.

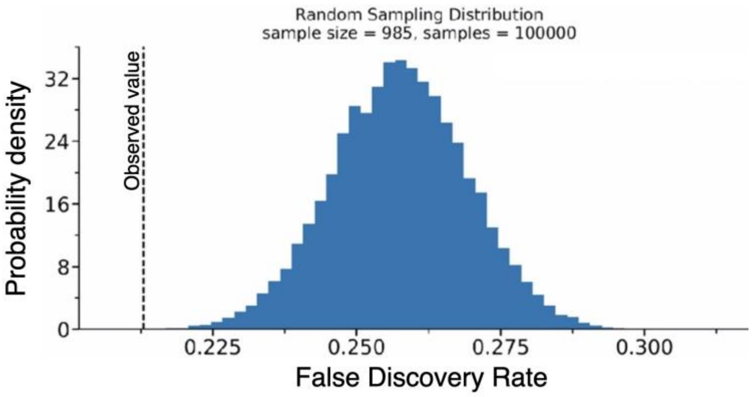


**Figure S2. Example of the random resampling procedure used to calculate the p-values for each phylogenetic filtering considered.** The graph shows the distribution of FoldX FDR values calculated on 100.000 random subsets of *N* entries from the Frenz et al dataset. *N* is the number of entries meeting the phylogenetic filtering threshold under scrutiny, in this example *N*=985. The bell-shaped curve evenly distributes around the mean value, which, as expected, corresponds to the FDR calculated from the whole dataset without applying any filters. The dashed vertical line corresponds to the observed FDR, as computed from the actual subset that meets the phylogenetic filtering threshold under scrutiny, and the p-value is calculated as the integral of this probability density function from 0 to this value (i.e., from 0 to the dashed line).

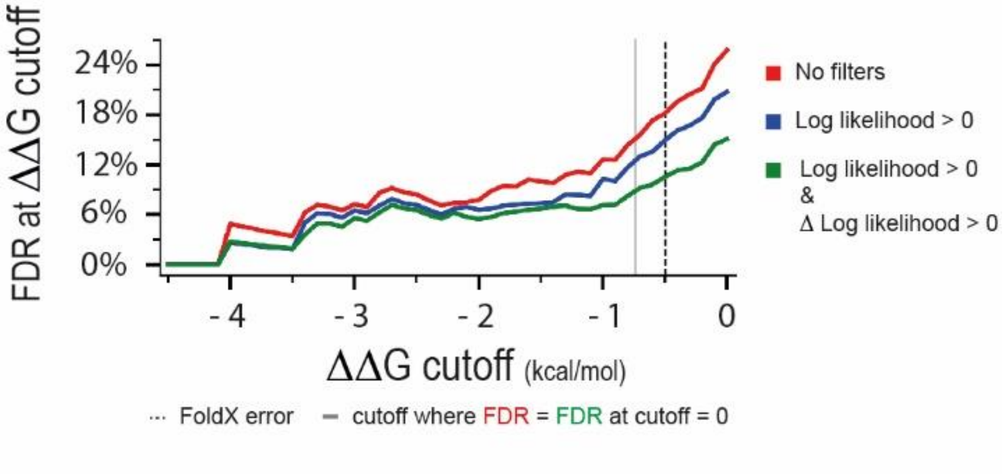


**Figure S3.** The graph shows FDR values calculated for the Frenz et al dataset at different filtering conditions (see legend) and at different ΔΔG cut-offs (x-axis). An FDR of 0% is obtained when the filtered dataset does not contain any False Positives. The dotted line shows the ΔΔG cut-off commonly regarded as a typical error of the FoldX predictions, corresponding to ± 0.5 kcal/mol. The grey line pinpoints the cut-off value (- 0.74 kcal/mol) that should be applied to the unfiltered dataset in order to obtain an FDR equal to the filtered dataset (green) at a cut-off value of 0.

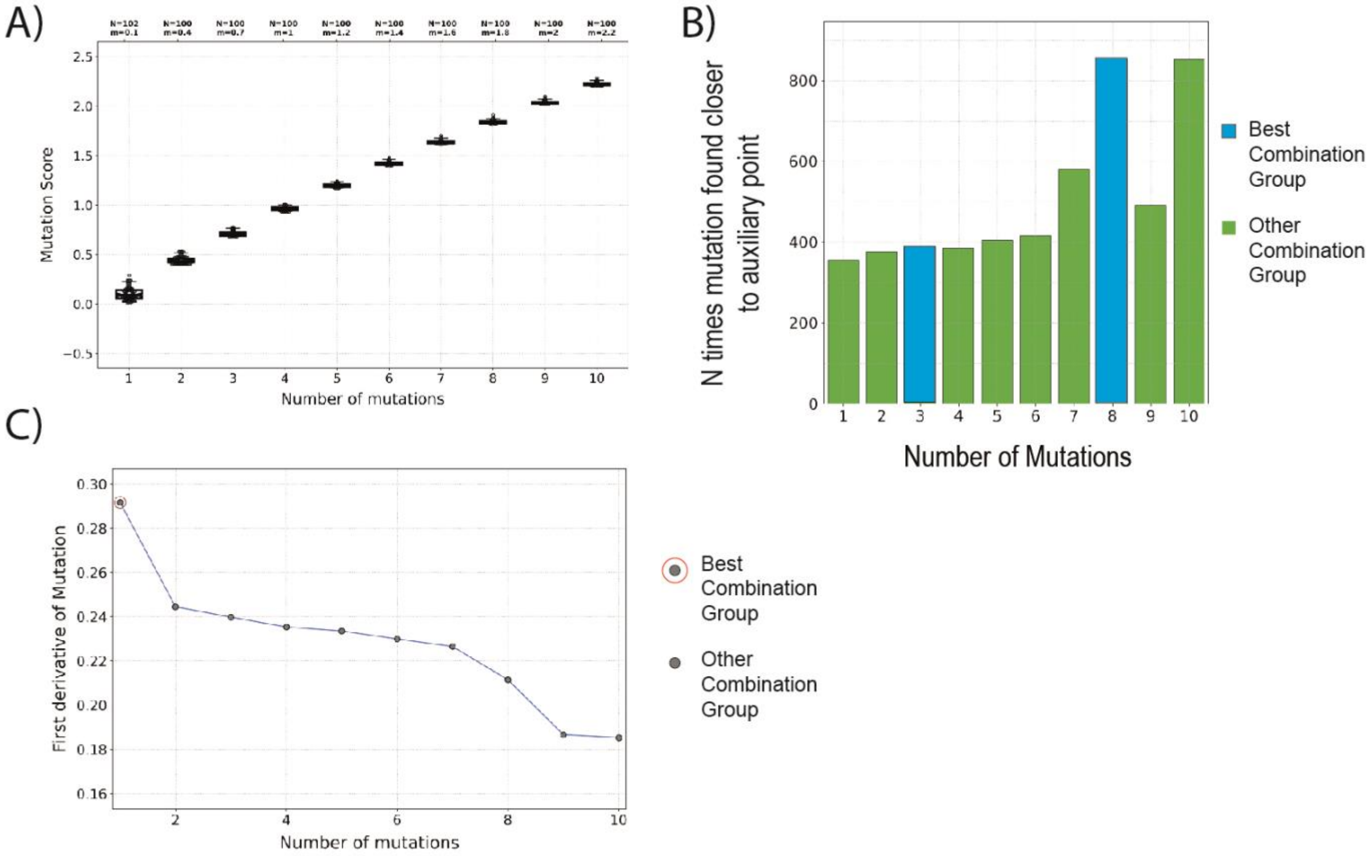


**Figure S4. Example of the best Combination Group analysis.** The graphs are obtained from running the pipeline on pdb 1bli (Amylase) with a PSSM obtained from an alignment with coverage 70 and identity 95, no excluded residues, and a limit of 10 simultaneous mutations in a combination. (**A**) The graph plots the mutation score against the number of mutations in combination. In this example, the mutation score grows monotonically although not in a linear way. This type of growth can be expected for larger proteins where there are plenty of potential mutations that can improve stability and solubility (or just one of the two). For smaller proteins, the growth often slows more rapidly sometimes reaching a plateau. (**B**) Weighted best combination group analysis. In the graph are highlighted in blue the combination groups identified as *Best Combination Groups.* The other groups are coloured in green. (**C**) The graph highlights the best mutation groups identified through what we called "first derivative analysis" in the Method section. Best combination groups are highlighted by a red circle in the graph. In this specific example, the first derivative decreases monotonically but this is not always the case and sometimes multiple best groups are identified in this way. The final set of best combination groups is the combination of the groups identified in plots (B) and (C).

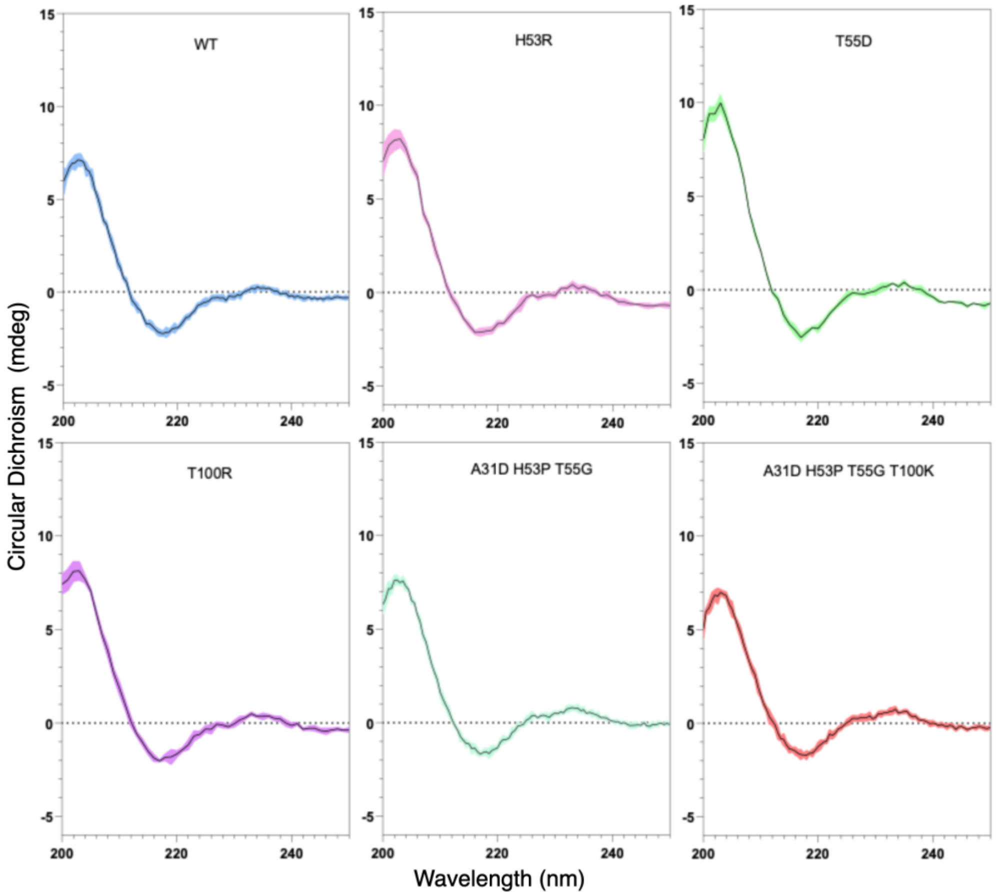


**Figure S5. Circular Dichroism spectra of WT and designed Nb.b201 variants.** CD spectra are plotted for WT, triple and quadruple mutants and all single mutants not contained in the quadruple (see title). The shape of the spectra with the minimum at 218nm is characteristic of a well-folded VHH domain.

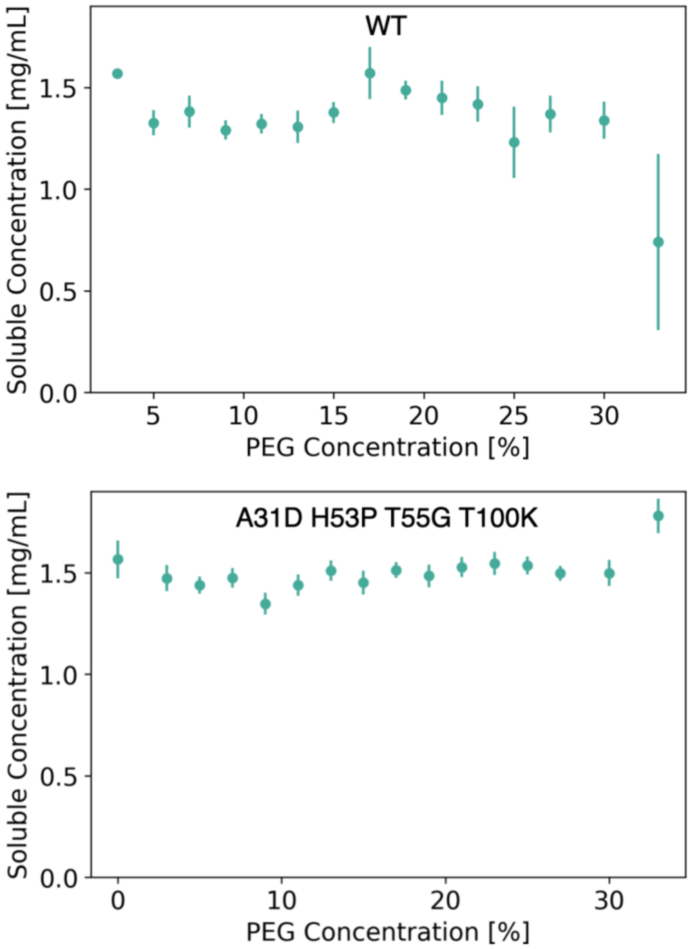


**Figure S6. PEG-precipitation of the WT and quadruple mutant.** The assay was carried out following the procedure described in detail in Ref. [16] using PEG-6K as a precipitant. The x-axis reports the PEG concentration in weight/volume percent. The y-axis reports the soluble concentration of the nanobody measured by absorbance at 280nm from the supernatant transferred to a fresh UV-transparent plate following centrifugation.

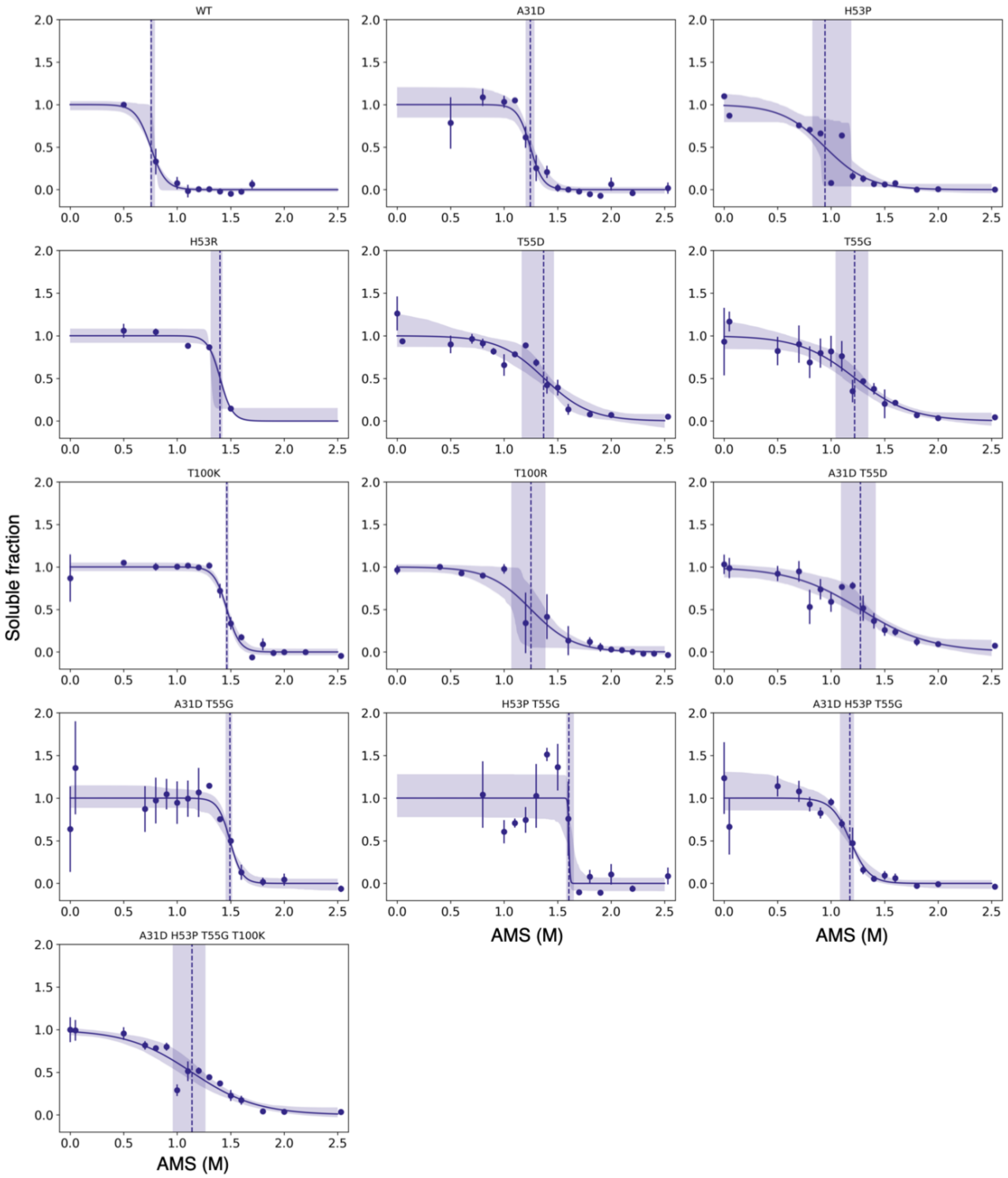


**Figure S7. AMS-precipitation.** The assay was carried out following the procedure described in detail in Ref. [16] using AMS instead of PEG as a precipitant, and 1 mg/mL final protein concentration in each well. The y-axis reports the normalised soluble concentration of the nanobody measured by absorbance at 280nm from the supernatant transferred to a fresh UV-transparent plate following centrifugation. The variant H53R has very few data point because we did not have enough purified protein material.

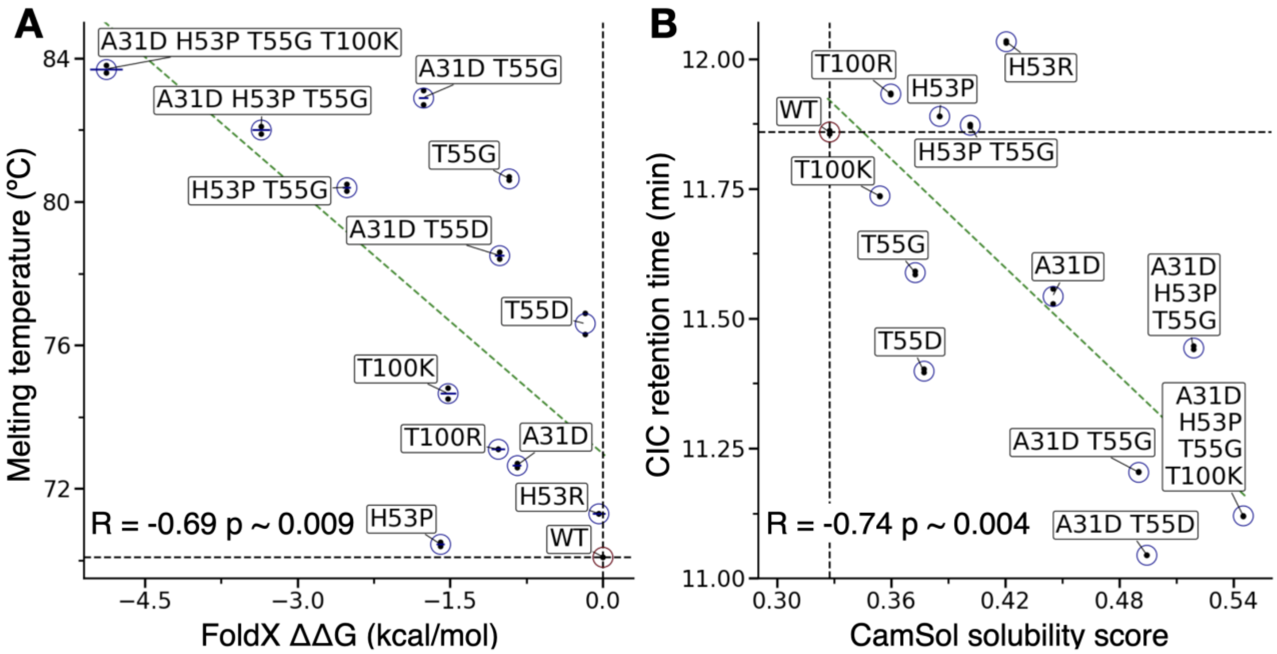


**Figure S8. Correlations between in silico predictions and experimental data for Nb.b201.** (**A**) Scatter plot of the measured melting temperature as a function of the calculated FoldX ΔΔG. (**B**) Scatter plot of the measured cross interaction chromatography (CIC) retention time as a function of the CamSol score. Pearson's coefficient of correlations (R) and corresponding p-values are reported on each panel. Experimental data are the same as **Fig. 4** of the main text, dashed black lines correspond to the WT values, the green line is the best linear trendline. If one excludes the two mutations to arginine (as cases where the known correlation between CIC and solubility breaks down, see main text) the correlation with CamSol becomes R = -0.79 p = 0.004. A correlation between CIC and CamSol on a library of 17 mAbs was previously reported, also with R = -0.79 (Ref. [17], figure 4 therein). A quantitative comparison between CamSol predictions and AMS-precipitation results was not carried out in this work because, given the rather low accuracy of this experimental assay, many variants had undistinguishable $AMS_{50\%}$ within the broad confidence intervals on this parameter (see **Fig. 4** and **Table 1**in the main text).

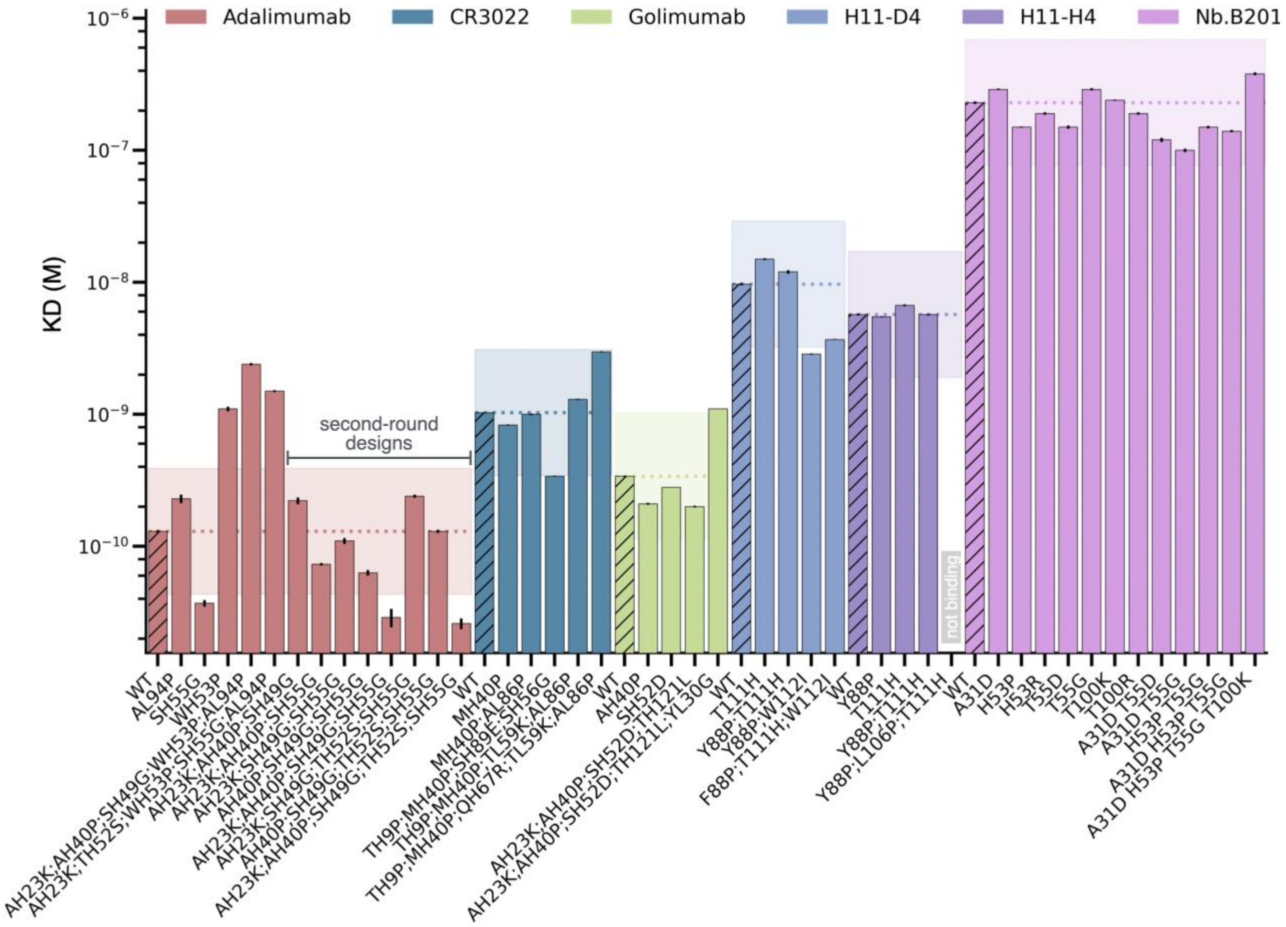


**Figure S9. Measured KD values of all antibody variants.** Bar plot of the KD values measured with BLI for the six WT antibodies (see legend) and all their designed variants (x-axis labels). Bars are coloured in groups according to the WT scFv or nanobody of each variant, and the bar corresponding to the WT is hatched. A coloured horizontal dotted line demarks the measurement of the WT in each group and serves as a guide to the eye. The semi-transparent band represents a confidence interval of +/- 3 folds over the KD value of the WT. We have used this boundary as cut-off to conclude on whether a design KD value is different from that of the WT or not. This cut-off, while arbitrary like any cut-off value, is based on our experience in estimating KDs using the BLI with relatively low molecular weight analytes, such as scFvs and Nbs, as it reflects the typical variability of repeated independent measurements across different batches. We also note that the BLI sensorgrams for adalimumab (see Supplementary Appendix) are characterised by an extremely slow dissociation. This slow dissociation makes the fitting of biding parameters rather unstable, which is likely affecting the fitted KD of those designs that appear to have extremely tight binding (KD below the WT confidence interval). Based on these measurements, we can conclude that these designs bind at least as well as the WT, but concluding that they bind better would require additional experiments beyond the scope of this work.

**Supplementary references**


(1) Frenz, B., Lewis, S. M., King, I., DiMaio, F., Park, H., and Song, Y. (2020) Prediction of Protein Mutational Free Energy: Benchmark and Sampling Improvements Increase Classification Accuracy. *Front. Bioeng. Biotechnol. 8*, 558247.

(2) Fariselli, P., Martelli, P. L., Savojardo, C., and Casadio, R. (2015) INPS: predicting the impact of non-synonymous variations on protein stability from sequence. *Bioinformatics 31*, 2816–2821.

(3) Schymkowitz, J., Borg, J., Stricher, F., Nys, R., Rousseau, F., and Serrano, L. (2005) The FoldX web server: an online force field. *Nucleic Acids Res. 33*, W382-8.

(4) Pei, J., and Grishin, N. V. (2001) AL2CO: calculation of positional conservation in a protein sequence alignment. *Bioinformatics 17*, 700–712.

(5) Dunbar, J., and Deane, C. M. (2015) ANARCI: antigen receptor numbering and receptor classification. *Bioinformatics* btv552.

(6) Honegger, A., and Plückthun, A. (2001) Yet Another Numbering Scheme for Immunoglobulin Variable Domains: An Automatic Modeling and Analysis Tool. *J. Mol. Biol. 309*, 657–670.

(7) Raybould, M. I. J., Marks, C., Krawczyk, K., Taddese, B., Nowak, J., Lewis, A. P., Bujotzek, A., Shi, J., and Deane, C. M. (2019) Five computational developability guidelines for therapeutic antibody profiling. *Proc. Natl. Acad. Sci. 33*, 201810576.

(8) Raybould, M. I. J., Marks, C., Lewis, A. P., Shi, J., Bujotzek, A., Taddese, B., and Deane, C. M. (2020) Thera-SAbDab: the Therapeutic Structural Antibody Database. *Nucleic Acids Res. 48*, D383–D388.

(9) Kovaltsuk, A., Leem, J., Kelm, S., Snowden, J., Deane, C. M., and Krawczyk, K. (2018) Observed Antibody Space: A Resource for Data Mining Next-Generation Sequencing of Antibody Repertoires. *J. Immunol. 201*, 2502–2509.

(10) Prihoda, D., Maamary, J., Waight, A., Juan, V., Fayadat-Dilman, L., Svozil, D., and Bitton, D. A. BioPhi: A platform for antibody design, humanization and humanness evaluation based on natural antibody repertoires and deep learning 19.

(11) Swindells, M. B., Porter, C. T., Couch, M., Hurst, J., Abhinandan, K. R., Nielsen, J. H., Macindoe, G., Hetherington, J., and Martin, A. C. R. (2017) abYsis: Integrated Antibody Sequence and Structure—Management, Analysis, and Prediction. *J. Mol. Biol. 429*, 356–364.

(12) Wilton, E. E., Opyr, M. P., Kailasam, S., Kothe, R. F., and Wieden, H.-J. (2018) sdAb-DB: The Single Domain Antibody Database. *ACS Synth. Biol. 7*, 2480–2484.

(13) Dunbar, J., Krawczyk, K., Leem, J., Baker, T., Fuchs, A., Georges, G., Shi, J., and Deane, C. M. (2014) SAbDab: the structural antibody database. *Nucleic Acids Res. 42*, D1140-6.

(14) Pettersen, E. F., Goddard, T. D., Huang, C. C., Couch, G. S., Greenblatt, D. M., Meng, E. C., and Ferrin, T. E. (2004) UCSF Chimera - A visualization system for exploratory research and analysis. *J. Comput. Chem. 25*, 1605–1612.

(15) Wolf Pérez, A.-M., Sormanni, P., Andersen, J. S., Sakhnini, L. I., Rodriguez-Leon, I., Bjelke, J. R., Gajhede, A. J., De Maria, L., Otzen, D. E., Vendruscolo, M., and Lorenzen, N. (2018) In vitro and in silico assessment of the developability of a designed monoclonal antibody library. *mAbs* 19420862.2018.1556082.

(16) Oeller, M., Sormanni, P., and Vendruscolo, M. (2021) An open-source automated PEG precipitation assay to measure the relative solubility of proteins with low material requirement. *Sci. Rep. 11*, 21932.

(17) Wolf Pérez, A.-M., Sormanni, P., Andersen, J. S., Sakhnini, L. I., Rodriguez-Leon, I., Bjelke, J. R., Gajhede, A. J., De Maria, L., Otzen, D. E., Vendruscolo, M., and Lorenzen, N. (2019) In vitro and in silico assessment of the developability of a designed monoclonal antibody library. *mAbs 11*, 388–400.

### SUPPLEMENTARY APPENDIX

#### BLI sensorgrams and fitted binding rates

##### Nb.b201 variants

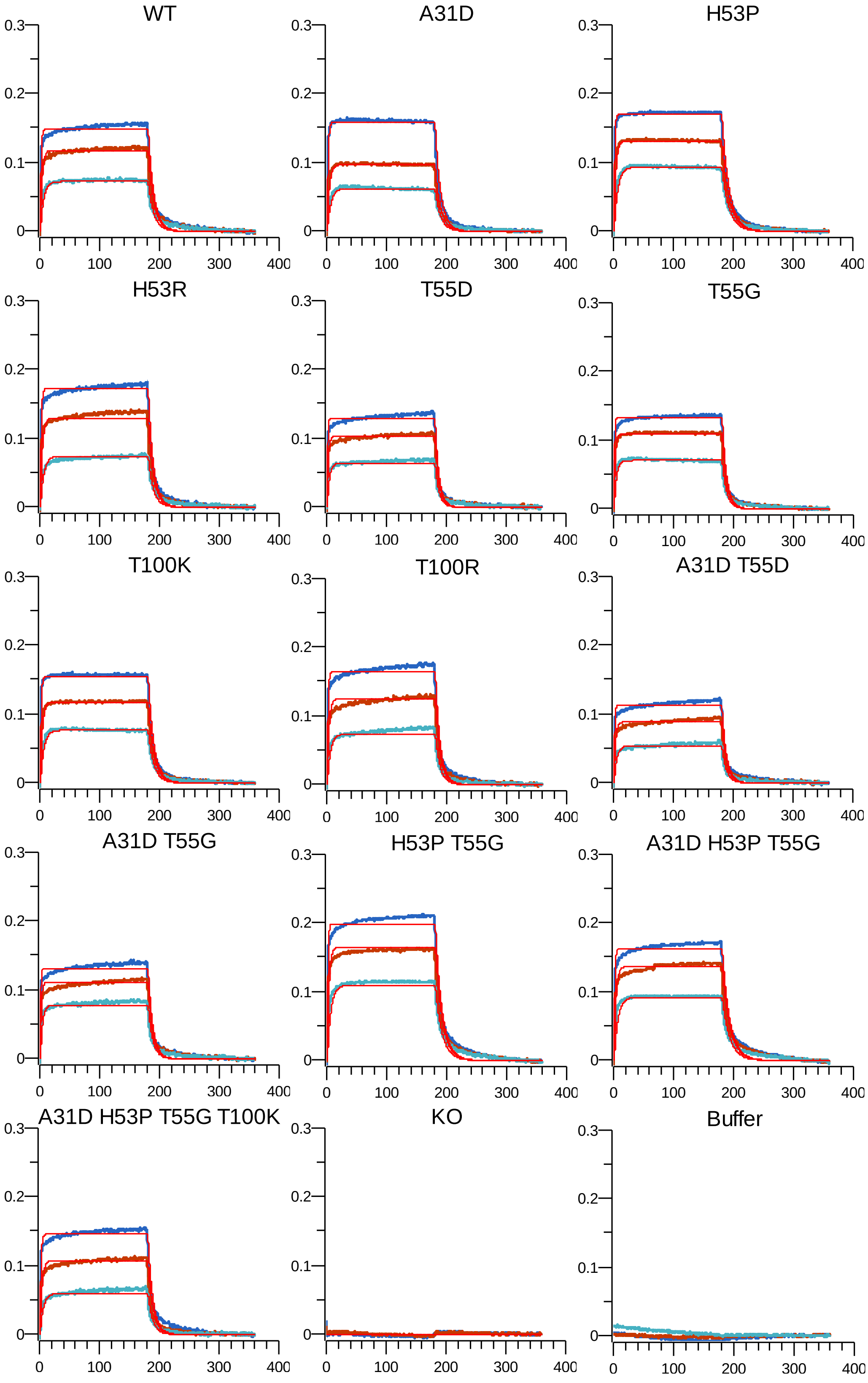


Y-axis is BLI binding signal in nm, x-axis is time in seconds. The KO variant (**Table S2**) contains 4 mutations in paratope residues that abolish binding and was employed as a negative control. Biotinylated HSA is immobilized on the sensors, and nanobody concentrations employed as analyte were 1500 (blue), 500 (red), and 167 (cyan) nM.

**ADALIMUMAB variants**

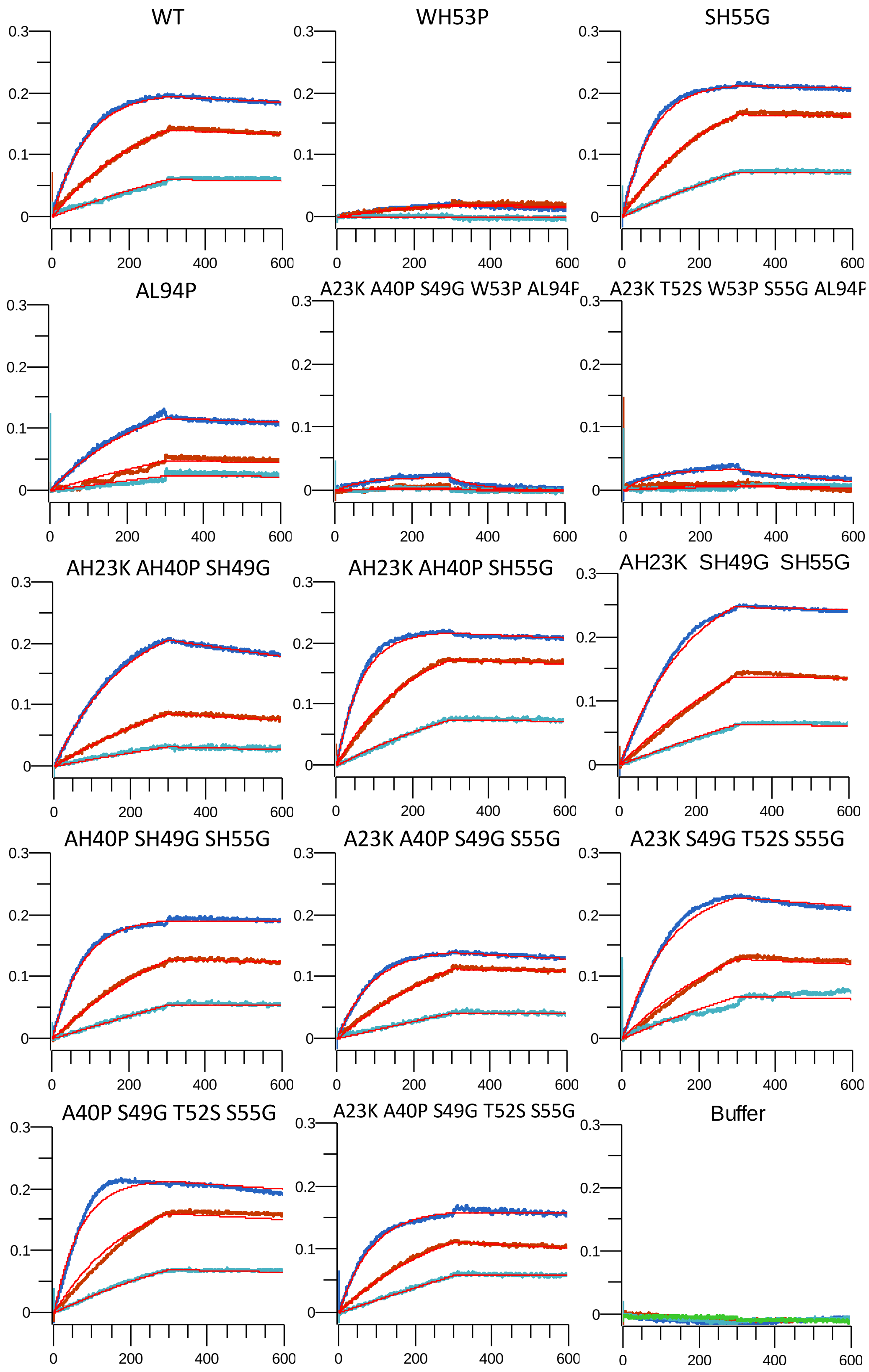


Y-axis is BLI binding signal in nm, x-axis is time in seconds. Biotinylated TNF-α is immobilized on the sensors, and scFv concentrations employed as analyte were 9 (blue), 3 (red), and 1 (cyan) nM.

**GOLIMUMAB variants**

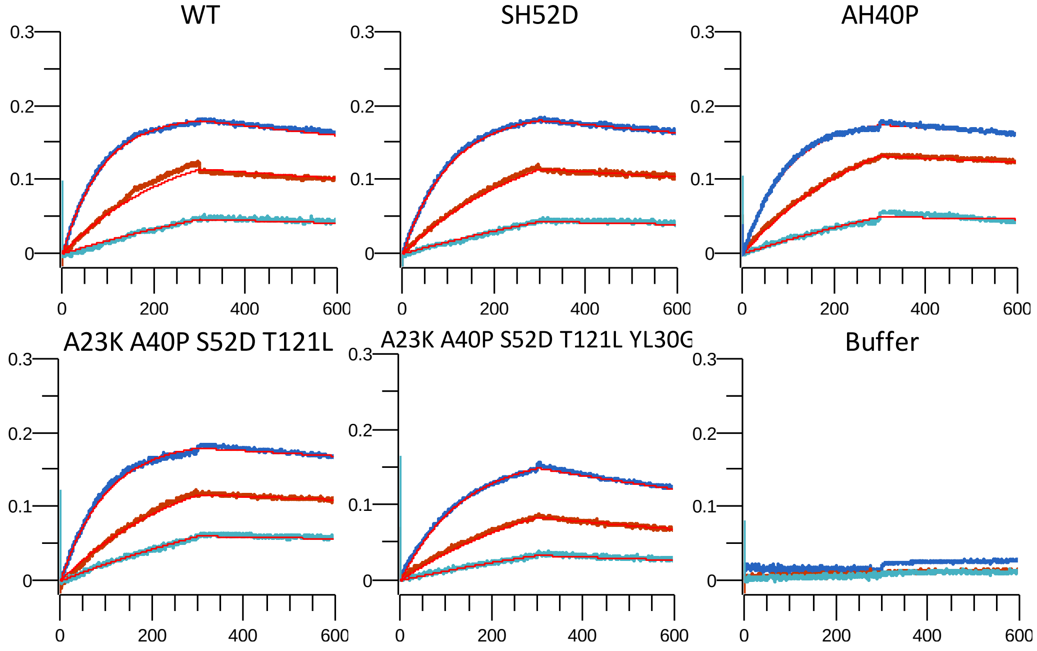


Y-axis is BLI binding signal in nm, x-axis is time in seconds. Biotinylated TNF-α is immobilized on the sensors, and scFv concentrations employed as analyte were 9 (blue), 3 (red), and 1 (cyan) nM.

**CR3022 variants**

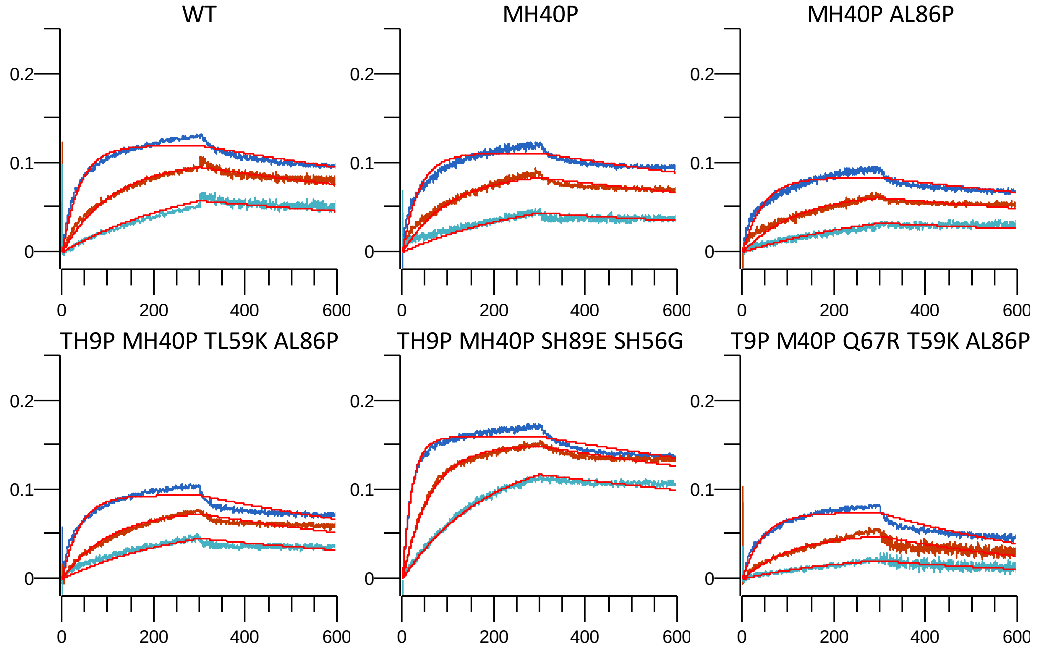


Y-axis is BLI binding signal in nm, x-axis is time in seconds. Biotinylated RBD is immobilized on the sensors, and scFv concentrations employed as analyte were 90 (blue), 30 (red), and 10 (cyan) nM.

**H11 variants:**

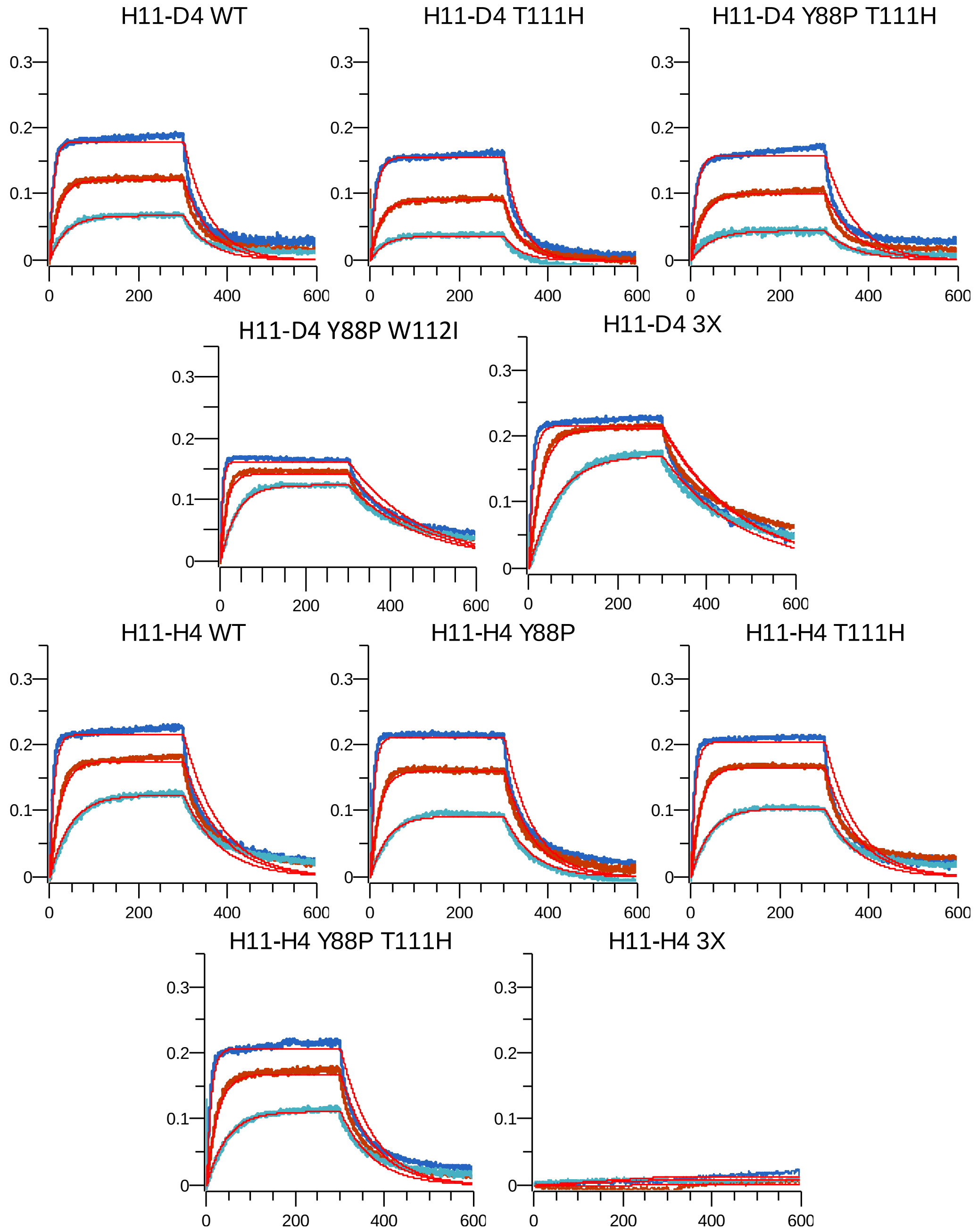


Y-axis is BLI binding signal in nm, x-axis is time in seconds. Biotinylated RBD is immobilized on the sensors, and nanobody concentrations employed as analyte were 90 (blue), 30 (red), and 10 (cyan) nM.

| **Antibody variant** | **KD (M)** | KD Error | **ka (1/Ms)** | ka Error | **kdiss (1/s)** | kdiss Error | Fit R^2 |
|---|---|---|---|---|---|---|---|
| Nb.B201 WT | **2.7E-07** | 1.6E-09 | **4.0E+05** | 1.7E+03 | **1.1E-01** | 4.1E-04 | 1.00 |
| Nb.B201 A31D | **2.8E-07** | 3.8E-09 | **3.7E+05** | 4.9E+03 | **1.0E-01** | 4.5E-04 | 1.00 |
| Nb.B201 H53P | **1.7E-07** | 1.1E-09 | **4.6E+05** | 2.3E+03 | **8.0E-02** | 3.3E-04 | 1.00 |
| Nb.B201 H53R | **3.0E-07** | 3.0E-09 | **3.8E+05** | 2.9E+03 | **1.1E-01** | 7.4E-04 | 0.99 |
| Nb.B201 T55D | **2.1E-07** | 2.5E-09 | **6.9E+05** | 6.0E+03 | **1.5E-01** | 1.1E-03 | 0.99 |
| Nb.B201 T55G | **1.7E-07** | 2.0E-09 | **6.2E+05** | 5.9E+03 | **1.0E-01** | 8.0E-04 | 0.99 |
| Nb.B201 T100K | **2.4E-07** | 3.4E-09 | **4.3E+05** | 5.7E+03 | **1.0E-01** | 4.5E-04 | 1.00 |
| Nb.B201 T100R | **2.7E-07** | 3.1E-09 | **4.4E+05** | 3.8E+03 | **1.2E-01** | 8.8E-04 | 0.99 |
| Nb.B201 A31D T55D | **2.5E-07** | 3.2E-09 | **5.1E+05** | 4.9E+03 | **1.3E-01** | 1.1E-03 | 0.99 |
| Nb.B201 A31D T55G | **1.4E-07** | 1.9E-09 | **8.7E+05** | 9.0E+03 | **1.2E-01** | 1.0E-03 | 0.99 |
| Nb.B201 H53P T55G | **1.9E-07** | 2.1E-09 | **4.4E+05** | 3.8E+03 | **8.2E-02** | 5.7E-04 | 0.99 |
| Nb.B201 A31D H53P T55G | **1.7E-07** | 2.0E-09 | **4.2E+05** | 4.0E+03 | **7.0E-02** | 5.1E-04 | 0.99 |
| Nb.B201 A31D H53P T55G T100K | **3.3E-07** | 3.8E-09 | **3.4E+05** | 2.9E+03 | **1.1E-01** | 8.4E-04 | 0.99 |
| Adalimumab WT | **1.3E-10** | 3.2E-12 | **1.2E+06** | 3.9E+03 | **1.5E-04** | 3.7E-06 | 1.00 |
| Adalimumab AH23K AH40P SH49G WH53P AL94P | **2.4E-09** | 4.9E-11 | **1.9E+06** | 3.6E+04 | **4.7E-03** | 3.6E-05 | 0.92 |
| Adalimumab AH23K TH52S WH53P SH55G AL94P | **1.5E-09** | 2.7E-11 | **1.6E+06** | 2.5E+04 | **2.3E-03** | 2.5E-05 | 0.95 |
| Adalimumab AL94P | **2.3E-10** | 1.7E-11 | **6.7E+05** | 8.1E+03 | **1.5E-04** | 1.1E-05 | 0.98 |
| Adalimumab SH55G | **3.7E-11** | 2.1E-12 | **1.4E+06** | 3.7E+03 | **5.2E-05** | 3.0E-06 | 1.00 |
| Adalimumab WH53P | **1.1E-09** | 4.5E-11 | **7.2E+05** | 2.0E+04 | **7.9E-04** | 2.4E-05 | 0.86 |
| Adalimumab2 AH23K AH40P SH49G | **2.2E-10** | 1.3E-11 | **1.7E+06** | 7.4E+04 | **3.7E-04** | 1.4E-05 | 0.97 |
| Adalimumab2 AH23K AH40P SH49G SH55G | **2.9E-11** | 4.6E-12 | **1.3E+06** | 3.6E+03 | **3.7E-05** | 6.0E-06 | 0.98 |
| Adalimumab2 AH23K AH40P SH49G TH52S SH55G | **2.6E-11** | 2.4E-12 | **1.4E+06** | 2.1E+03 | **3.7E-05** | 3.4E-06 | 0.99 |
| Adalimumab2 AH23K AH40P SH55G | **7.3E-11** | 1.6E-12 | **1.7E+06** | 4.1E+03 | **1.3E-04** | 2.8E-06 | 1.00 |
| Adalimumab2 AH23K SH49G SH55G | **1.1E-10** | 5.8E-12 | **6.1E+05** | 2.7E+03 | **6.8E-05** | 3.5E-06 | 1.00 |
| Adalimumab2 AH23K SH49G TH52S SH55G | **2.4E-10** | 6.4E-12 | **9.1E+05** | 5.2E+03 | **2.2E-04** | 5.8E-06 | 0.99 |
| Adalimumab2 AH40P SH49G SH55G | **6.3E-11** | 3.1E-12 | **1.2E+06** | 1.9E+03 | **7.3E-05** | 3.6E-06 | 0.99 |
| Adalimumab2 AH40P SH49G TH52S SH55G | **1.3E-10** | 3.8E-12 | **1.6E+06** | 8.3E+03 | **2.1E-04** | 6.0E-06 | 0.99 |
| Golimumab WT | **3.4E-10** | 3.8E-12 | **1.3E+06** | 5.0E+03 | **4.3E-04** | 4.5E-06 | 0.99 |
| Golimumab SH52D | **2.8E-10** | 2.1E-12 | **1.1E+06** | 2.3E+03 | **3.2E-04** | 2.2E-06 | 1.00 |
| Golimumab AH40P | **2.1E-10** | 3.5E-12 | **1.1E+06** | 3.6E+03 | **2.2E-04** | 3.6E-06 | 1.00 |
| Golimumab AH23K AH40P SH52D TH121L YL30G | **1.1E-09** | 5.8E-12 | **6.7E+05** | 2.3E+03 | **7.2E-04** | 3.0E-06 | 1.00 |
| Golimumab AH23K AH40P SH52D TH121L | **2.0E-10** | 2.7E-12 | **1.2E+06** | 3.3E+03 | **2.3E-04** | 3.0E-06 | 1.00 |
| CR3022 WT | **1.0E-09** | 8.9E-12 | **8.9E+05** | 4.6E+03 | **9.1E-04** | 6.3E-06 | 0.98 |
| CR3022 TH9P MH40P SH89E SH56G | **3.4E-10** | 2.7E-12 | **1.6E+06** | 5.4E+03 | **5.4E-04** | 3.9E-06 | 0.99 |
| CR3022 TH9P MH40P QH67R TL59K AL86P | **3.0E-09** | 3.3E-11 | **7.0E+05** | 6.4E+03 | **2.1E-03** | 1.3E-05 | 0.97 |
| CR3022 TH9P MH40P TL59K AL86P | **1.3E-09** | 1.5E-11 | **8.4E+05** | 6.1E+03 | **1.1E-03** | 9.1E-06 | 0.96 |
| CR3022 MH40P AL86P | **1.0E-09** | 1.4E-11 | **7.3E+05** | 5.1E+03 | **7.4E-04** | 8.6E-06 | 0.97 |
| CR3022 MH40P | **8.3E-10** | 9.7E-12 | **8.9E+05** | 5.4E+03 | **7.4E-04** | 7.4E-06 | 0.98 |
| H11-D4 WT | **9.7E-09** | 1.9E-10 | **1.2E+06** | 2.1E+04 | **1.1E-02** | 6.3E-05 | 0.94 |
| H11-D4 F88P T111H W112I | **3.7E-09** | 2.4E-11 | **1.1E+06** | 6.8E+03 | **4.1E-03** | 9.6E-06 | 0.98 |
| H11-D4 T111H | **1.5E-08** | 2.4E-10 | **9.3E+05** | 1.4E+04 | **1.4E-02** | 6.9E-05 | 0.97 |
| H11-D4 Y88P T111H | **1.2E-08** | 3.5E-10 | **6.7E+05** | 1.9E+04 | **7.8E-03** | 7.4E-05 | 0.81 |
| H11-D4 Y88P W112I | **2.9E-09** | 2.7E-11 | **2.1E+06** | 1.9E+04 | **6.0E-03** | 1.7E-05 | 0.972 |
| H11-H4 WT | **5.7E-09** | 1.0E-10 | **2.3E+06** | 4.1E+04 | **1.3E-02** | 7.3E-05 | 0.95 |
| H11-H4 T111H | **6.7E-09** | 1.2E-10 | **1.6E+06** | 2.7E+04 | **1.1E-02** | 5.8E-05 | 0.96 |
| H11-H4 Y88P | **5.5E-09** | 6.4E-11 | **2.1E+06** | 2.3E+04 | **1.1E-02** | 4.1E-05 | 0.98 |
| H11-H4 Y88P L106P T111H | - | - | - | - | - | - | - |
| H11-H4 Y88P T111H | **5.7E-09** | 7.1E-11 | **1.5E+06** | 1.8E+04 | **8.6E-03** | 3.5E-05 | 0.98 |

**Table with the fitted biding parameters from the BLI sensorgrams.** The association and dissociation rates, and affinities of the antibodies were evaluated in the Data analysis HT 12.0 software from FORTÉBIO, where the data was fitted to a 1:1 binding model using unlinked Rmax values. The 2 in adalimumab2 denotes second round designs, the WT is always adalimumab.
