## Supplementary files 1 to 9 for "Automated optimisation of solubility and conformational stability of antibodies and proteins": SF1 Amylase 1bli report.pdf

### CamSol design report for input 1bli

**General remarks.** This CamSol design procedure is aimed at optimising the solubility of the target protein while retaining or improving its native state stability. This task is achieved by combining four approaches. (1) The CamSol structurally-corrected prediction is carried out to identify potential aggregation hotspots on the surface of the target protein, whose presence may elicit aggregation from the native state. Those residues contributing most to the hotspots aggregation propensity are flagged as candidate mutation sites. (2) The CamSol intrinsic profile is used to identify further mutation sites that contribute strongly to the poor solubility of the unfolded state. In fact, thermal fluctuations may lead to the transient exposure of otherwise buried aggregation-promoting regions leading to aggregation from partially or fully unfolded states. After (1) & (2) the CamSol intrinsic algorithm is used to quickly screen all possible point mutations at these identified sites to create a long-list of candidate mutations that would in principle increase solubility. (3) Sequences homologous to the target protein are automatically identified, and a multiple-sequence alignment (MSA) is performed between them. A position-specific scoring-matrix (PSSM) is calculated from this alignment to identify those residues that are most likely to be found at each position. Mutations to residues more conserved (i.e. with higher PSSM frequency) in the alignments of homologous sequences have been shown to correlate with increased native-state stability (albeit the correlation is not perfect), and more generally mutations to such residues are much better tolerated, and highly unlikely to be deleterious for the native fold. This PSSM is used to filter the long-list of candidate solubilising mutations by selecting only those mutations that increase the frequency at the position under scrutiny. (5) A structure-based atomistic prediction of the stability change upon mutation is carried out with the energy function Fold-X for each shortlisted mutations. Mutations with calculated DDG<0 are predicted to be stabilising. The correlation between measured stability and Fold-X predicted DDG is statistically significant but far from perfect. However, the atomistic calculations of Fold-X are fully independent from, and therefore highly complementary to, the evolutionary frequency difference calculated from the PSSM. Therefore mutations with increased evolutionary frequency and with a negative calculated DDG should be stabilising, or at least should not negatively impact stability. Conversely, the CamSol methods has been shown to be highly quantitative in recapitulating the effect of mutations on measured solubility in different contexts (R~0.9). Consequently mutations that (i) increase the CamSol solubility score, (ii) have a FoldX DDG smaller than 0 and (iii) increase the PSSM frequency are expected to increase protein solubility, while not affecting or even improving native fold stability.

**CamSol analysis of input pdb file: 1bli.pdb**

These amino acids are excluded from the list of potential substitution targets: C, M

**Sequences extracted from input pdb and used for analysis:**
lower case residues (if any) are residues of missing coordinates but present in the SEQRES field of the input pdb file. Such residues are considered for the solubility calculation, but are never targeted for mutations and their potential impact on stability is not considered.

```
> 1bli:A
anLNGTLMQYFEWYMPNDGQHWKRLQNDSAYLAEHGITAVWIPPAYKGTSQADVGYGAYDLYDLGEFHQKGTVRTKYGTKGELQSAIKSLHSRDINVYGDVVINHKGGADATEDVTAVEV
DPADRNRVISGEHLIKAWTHFHFPGRGSTYSDFKWHWYHFDGTDWDESRKLNRIYKFQGKAWDWEVSNEFGNYDYLMYADIDYDHPDVAAEIKRWGTWYANELQLDGFRLDAVKHIKFSF
LRDWVNHVREKTGKEMFTVAEYWSYDLGALENYLNKTNFNHSVFDVPLHYQFHAASTQGGGYDMRKLLNGTVVSKHPLKSVTFVDNHDTQPGQSLESTVQTWFKPLAYAFILTRESGYPQ
VFYGDMYGTKGDSQREIPALKHKIEPILKARKQYAYGAQHDYFDHHDIVGWTREGDSSVANSGLAALITDGPGGAKRMYVGRQNAGETWHDITGNRSEPVVINSAGWGEFHVNGGSVSIY
VQR
```

Using log-likelihood pssm. Considering only candidate mutations with positive enrichment (log-likelihood > 0), and further restricting the space of candidate substitutions at each position to those residues that are more likely than the WT one

#### PSSM used to pick candidate mutations

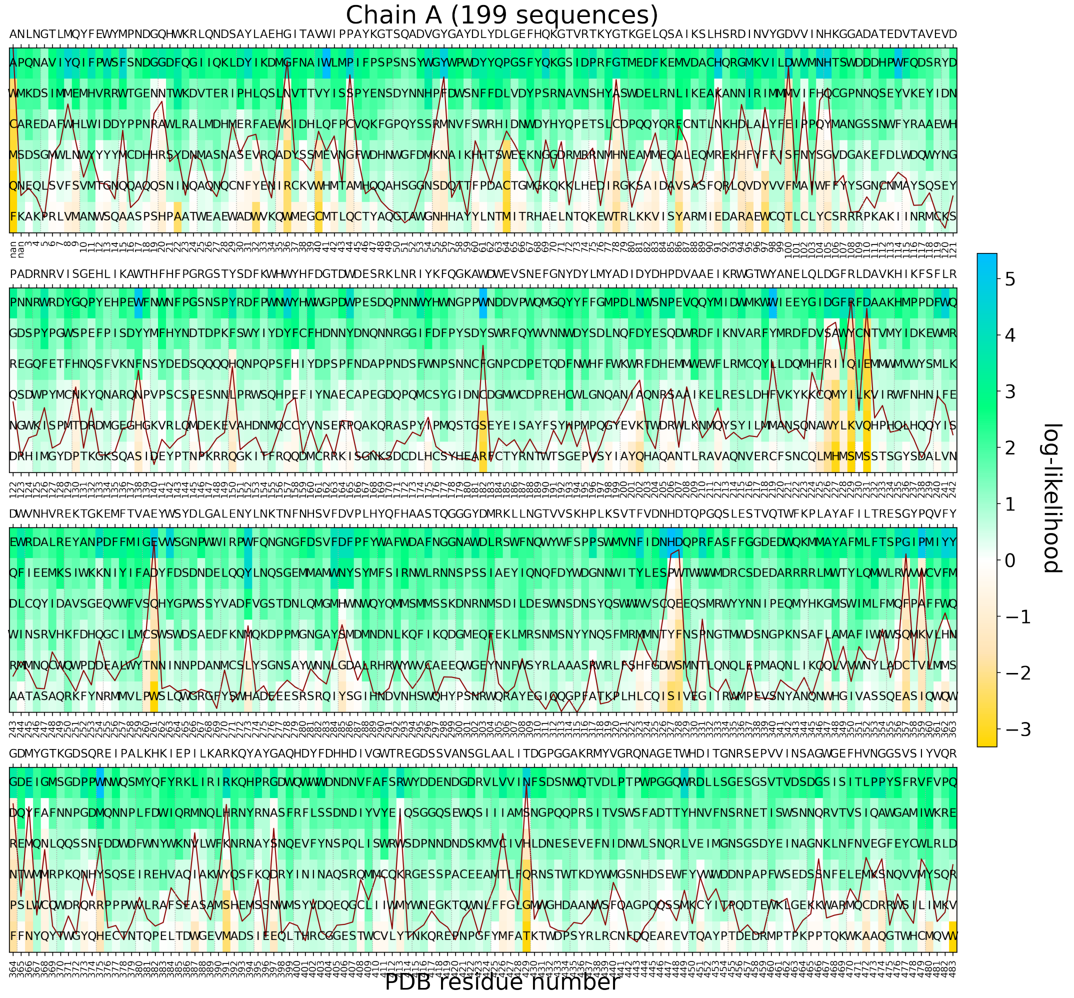


Position-specific scoring matrix (PSSM), as calculated from a multiple-sequence alignment (MSA) of similar sequences. The observed residue frequency (color-bar) is used to select candidate amino acid substitutions. The sequence above the panels is the wild-type (input) sequence as read from the alignment. The red line (if present) is the conservation index of each position (high means position highly conserved). PSSM of chain A read from file hhblits_fullQT_3iteration1kseqs_1bli_chain_A.hhm, computed from 199 aligned sequences (Neff=8).

#### CamSol intrinsic and structurally corrected profiles

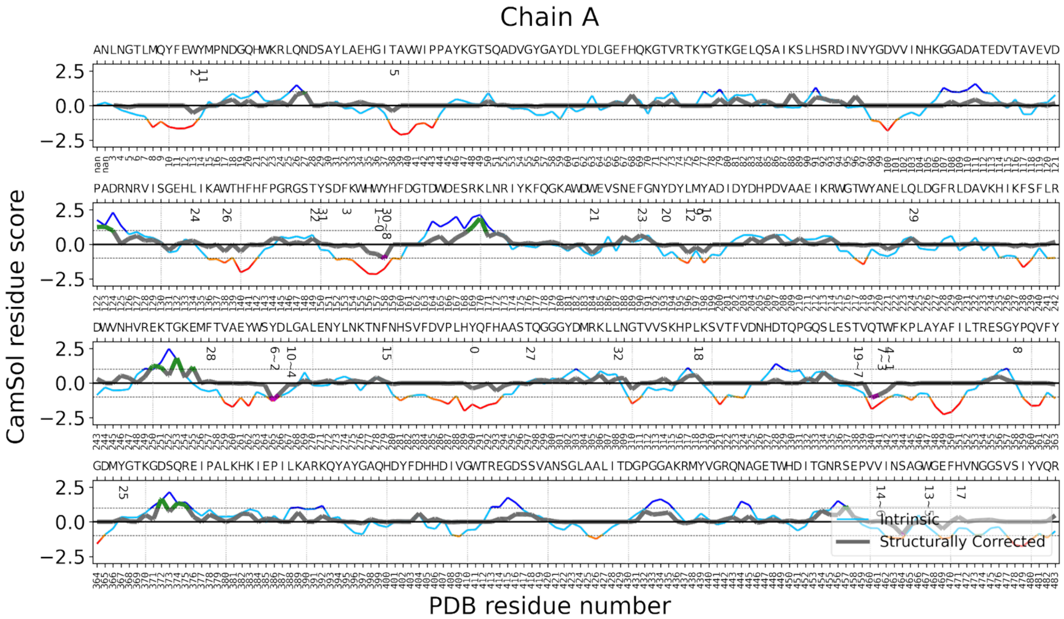


The CamSol intrinsic profile is colour-coded red to blue, where red means aggregation-prone and blue aggregation-resistant. It is common for folded proteins to have large aggregation-prone regions in their intrinsic profile that typically drive the hydrophobic collapse during folding. The structurally corrected profile is color-coded in gray/green/magenta, regions of low negative scores (magenta) are potential aggregation hotspots, regions of high score (green) are solubility promoting. Numbers below the amino acid sequence at the top denote potential mutation sites, identified according to their contribution to the solubility, as well as their accessibility to the solvent.

### DESIGN PIPELINE RESULTS: Best Models

##### Table with identified best combinations of mutations

| Design Name | Number of Mutations | Mutations in Combination | Stability Rank | Solubility Rank | Theoretica PI |
|---|---|---|---|---|---|
| model_18 | 9 | AA45P,GA81E,NA96R,QA224G,TA338P,TA341P,SA356R,NA473P,GA474P | 2 | 5 | 6.112 |
| model_7 | 9 | AA45P,GA81E,YA198G,QA224G,TA338P,TA341P,SA356R,NA473P,GA474P | 2 | 4 | 6.049 |
| model_5 | 9 | AA45P,GA81E,AA209R,QA224G,TA338P,TA341P,SA356R,NA473P,GA474P | 2 | 5 | 6.112 |
| model_12 | 8 | AA45P,GA81E,QA224G,TA338P,TA341P,SA356R,NA473P,GA474P | 3 | 6 | 6.049 |
| model_9 | 8 | AA45P,GA81E,NA96R,QA224G,TA338P,TA341P,NA473P,GA474P | 4 | 6 | 6.049 |
| model_13 | 8 | AA45P,GA81E,NA96R,TA338P,TA341P,SA356R,NA473P,GA474P | 3 | 6 | 6.112 |
| model_20 | 6 | AA45P,GA81E,TA338P,TA341P,NA473P,GA474P | 5 | 7 | 5.987 |
| model_22 | 6 | AA45P,GA81E,QA224G,TA338P,TA341P,GA474P | 5 | 8 | 5.987 |
| model_21 | 6 | AA45P,GA81E,TA338P,TA341P,SA356R,GA474P | 5 | 7 | 6.049 |
| model_3 | 2 | AA45P,GA81E | 10 | 11 | 5.987 |
| model_15 | 2 | AA45P,GA474P | 10 | 12 | 6.047 |
| model_8 | 2 | GA81E,GA474P | 10 | 11 | 5.987 |
| model_6 | 1 | AA45P | 11 | 12 | 6.047 |
| model_17 | 1 | GA81E | 11 | 12 | 5.987 |
| model_1 | 1 | GA474P | 11 | 12 | 6.047 |
| model_4 | 12 | AA45P,GA81E,NA96R,YA198G,AA209R,QA224G,TA338P,TA341P,SA356R,VA461K,NA473P,GA474P | 1 | 1 | 6.247 |
| model_11 | 11 | AA45P,GA81E,NA96R,YA198G,AA209R,QA224G,TA338P,TA341P,SA356R,NA473P,GA474P | 1 | 3 | 6.178 |
| model_2 | 10 | AA45P,GA81E,NA96R,YA198G,QA224G,TA338P,TA341P,SA356R,NA473P,GA474P | 2 | 3 | 6.112 |
| model_19 | 7 | AA45P,GA81E,QA224G,TA338P,TA341P,NA473P,GA474P | 4 | 7 | 5.987 |
| model_16 | 5 | AA45P,GA81E,TA338P,TA341P,GA474P | 6 | 8 | 5.987 |
| model_14 | 4 | AA45P,GA81E,TA341P,GA474P | 7 | 9 | 5.987 |
| model_10 | 3 | AA45P,GA81E,GA474P | 8 | 11 | 5.987 |
| WT | 0 | WT | 12 | 12 | 6.047 |

This table contains information on the identified mutation combinations, selected on the basis of their contribution to the overall solubility and stability of the protein. The column 'design name' identifies the pdb file bearing the mutations listed in the column 'mutations in combination'. Combinations belonging to the best combination groups (i.e., combinations containing 1, 2, 6, 8, 9 mutants) are listed first and sorted according to their mutation score. The remaining combinations, not identified as best mutation combinations, are listed afterwards and are sorted according to their mutation score. Mutation combinations are given a solubility and stability rank according to their delta CamSol score and DDG value respectively (i.e. they are binned starting from 1 corresponding the best predicted). The top ranking combination is usually one with the highest 'Number of Mutations'. However, because with each added mutation the chances of introducing a false positive increase, the best combination groups (top rows in the table) embody the best tradeoff between low number of mutations and high gain in solubility and stability. All combinations listed are expected to increase both solubility and stability (or one without compromising the other) and contain single point mutants that are enriched in the input PSSM.

##### 9 Simultaneous Mutations:

**Model Name: Model_18**

Mutations by chain

Chain A : A45P,G81E,N96R,Q224G,T338P,T341P,S356R,N473P,G474P

Solubility Ranking: 5 | Delta CamSol score: 0.199

Stability Ranking: 2 | FoldX DDG: -15.199 kcal/mol

Mutation Score: 2.239

> Model_18 chain A Optimized_1bli_Repair | A45P G81E N96R Q224G T338P T341P S356R N473P G474P

LNGTLMQYFEWYMPNDGQHWKRLQNDSAYLAEHGITAVWIPP**P**YKGTSQADVGYGAYDLYDLGEFHQKGTVRTKYGTK**E**ELQSAIKSLHSRDI**R**VYGDVVINHKGGADATEDVTAVEVDPA

DRNRVISGEHLIKAWTHFHFPGRGSTYSDFKWHWYHFDGTDWDESRKLNRIYKFQGKAWDWEVSNEFGNYDYLMYADIDYDHPDVAAEIKRWGTWYANEL**G**LDGFRLDAVKHIKFSFLRD

WVNHVREKTGKEMFTVAEYWSYDLGALENYLNKTNFNHSVFDVPLHYQFHAASTQGGGYDMRKLLNGTVVSKHPLKSVTFVDNHDTQPGQSLES**P**VQ**P**WFKPLAYAFILTRE**R**GYPQVFY

GDMYGTKGDSQREIPALKHKIEPILKARKQYAYGAQHDYFDHHDIVGWTREGDSSVANSGLAALITDGPGGAKRMYVGRQNAGETWHDITGNRSEPVVINSAGWGEFHV**PP**GSVSIYVQR

**Model Name: Model_7**

Mutations by chain

Chain A : A45P,G81E,Y198G,Q224G,T338P,T341P,S356R,N473P,G474P

Solubility Ranking: 4 | Delta CamSol score: 0.242

Stability Ranking: 2 | FoldX DDG: -15.351 kcal/mol

Mutation Score: 2.235

> Model_7 chain A Optimized_1bli_Repair | A45P G81E Y198G Q224G T338P T341P S356R N473P G474P

LNGTLMQYFEWYMPNDGQHWKRLQNDSAYLAEHGITAVWIPP**P**YKGTSQADVGYGAYDLYDLGEFHQKGTVRTKYGTK**E**ELQSAIKSLHSRDINVYGDVVINHKGGADATEDVTAVEVDPA

DRNRVISGEHLIKAWTHFHFPGRGSTYSDFKWHWYHFDGTDWDESRKLNRIYKFQGKAWDWEVSNEFGNYDYLM**G**ADIDYDHPDVAAEIKRWGTWYANEL**G**LDGFRLDAVKHIKFSFLRD

WVNHVREKTGKEMFTVAEYWSYDLGALENYLNKTNFNHSVFDVPLHYQFHAASTQGGGYDMRKLLNGTVVSKHPLKSVTFVDNHDTQPGQSLES**P**VQ**P**WFKPLAYAFILTRE**R**GYPQVFY

GDMYGTKGDSQREIPALKHKIEPILKARKQYAYGAQHDYFDHHDIVGWTREGDSSVANSGLAALITDGPGGAKRMYVGRQNAGETWHDITGNRSEPVVINSAGWGEFHV**PP**GSVSIYVQR

**Model Name: Model_5**

Mutations by chain

Chain A : A45P,G81E,A209R,Q224G,T338P,T341P,S356R,N473P,G474P

Solubility Ranking: 5 | Delta CamSol score: 0.209

Stability Ranking: 2 | FoldX DDG: -15.156 kcal/mol

Mutation Score: 2.234

> Model_5 chain A Optimized_1bli_Repair | A45P G81E A209R Q224G T338P T341P S356R N473P G474P

LNGTLMQYFEWYMPNDGQHWKRLQNDSAYLAEHGITAVWIPP**P**YKGTSQADVGYGAYDLYDLGEFHQKGTVRTKYGTK**E**ELQSAIKSLHSRDINVYGDVVINHKGGADATEDVTAVEVDPA

DRNRVISGEHLIKAWTHFHFPGRGSTYSDFKWHWYHFDGTDWDESRKLNRIYKFQGKAWDWEVSNEFGNYDYLMYADIDYDHPDV**R**AEIKRWGTWYANEL**G**LDGFRLDAVKHIKFSFLRD

WVNHVREKTGKEMFTVAEYWSYDLGALENYLNKTNFNHSVFDVPLHYQFHAASTQGGGYDMRKLLNGTVVSKHPLKSVTFVDNHDTQPGQSLES**P**VQ**P**WFKPLAYAFILTRE**R**GYPQVFY

GDMYGTKGDSQREIPALKHKIEPILKARKQYAYGAQHDYFDHHDIVGWTREGDSSVANSGLAALITDGPGGAKRMYVGRQNAGETWHDITGNRSEPVVINSAGWGEFHV**PP**GSVSIYVQR

#### 8 Simultaneous Mutations:

**Model Name: Model_12**

Mutations by chain

Chain A : A45P,G81E,Q224G,T338P,T341P,S356R,N473P,G474P

Solubility Ranking: 6 | Delta CamSol score: 0.188

Stability Ranking: 3 | FoldX DDG: -14.27 kcal/mol

Mutation Score: 2.057

> Model_12 chain A Optimized_1bli_Repair | A45P G81E Q224G T338P T341P S356R N473P G474P

LNGTLMQYFEWYMPNDGQHWKRLQNDSAYLAEHGITAVWIPP**P**YKGTSQADVGYGAYDLYDLGEFHQKGTVRTKYGTK**E**ELQSAIKSLHSRDINVYGDVVINHKGGADATEDVTAVEVDPA

DRNRVISGEHLIKAWTHFHFPGRGSTYSDFKWHWYHFDGTDWDESRKLNRIYKFQGKAWDWEVSNEFGNYDYLMYADIDYDHPDVAAEIKRWGTWYANEL**G**LDGFRLDAVKHIKFSFLRD

WVNHVREKTGKEMFTVAEYWSYDLGALENYLNKTNFNHSVFDVPLHYQFHAASTQGGGYDMRKLLNGTVVSKHPLKSVTFVDNHDTQPGQSLES**P**VQ**P**WFKPLAYAFILTRE**R**GYPQVFY

GDMYGTKGDSQREIPALKHKIEPILKARKQYAYGAQHDYFDHHDIVGWTREGDSSVANSGLAALITDGPGGAKRMYVGRQNAGETWHDITGNRSEPVVINSAGWGEFHV**PP**GSVSIYVQR

**Model Name: Model_9**

Mutations by chain

Chain A : A45P,G81E,N96R,Q224G,T338P,T341P,N473P,G474P

Solubility Ranking: 6 | Delta CamSol score: 0.173

Stability Ranking: 4 | FoldX DDG: -13.43 kcal/mol

Mutation Score: 2.017

> Model_9 chain A Optimized_1bli_Repair | A45P G81E N96R Q224G T338P T341P N473P G474P

LNGTLMQYFEWYMPNDGQHWKRLQNDSAYLAEHGITAVWIPP**P**YKGTSQADVGYGAYDLYDLGEFHQKGTVRTKYGTK**E**ELQSAIKSLHSRDI**R**VYGDVVINHKGGADATEDVTAVEVDPA

DRNRVISGEHLIKAWTHFHFPGRGSTYSDFKWHWYHFDGTDWDESRKLNRIYKFQGKAWDWEVSNEFGNYDYLMYADIDYDHPDVAAEIKRWGTWYANEL**G**LDGFRLDAVKHIKFSFLRD

WVNHVREKTGKEMFTVAEYWSYDLGALENYLNKTNFNHSVFDVPLHYQFHAASTQGGGYDMRKLLNGTVVSKHPLKSVTFVDNHDTQPGQSLES**P**VQ**P**WFKPLAYAFILTRESGYPQVFY

GDMYGTKGDSQREIPALKHKIEPILKARKQYAYGAQHDYFDHHDIVGWTREGDSSVANSGLAALITDGPGGAKRMYVGRQNAGETWHDITGNRSEPVVINSAGWGEFHV**PP**GSVSIYVQR

**Model Name: Model_13**

Mutations by chain

Chain A : A45P,G81E,N96R,T338P,T341P,S356R,N473P,G474P

Solubility Ranking: 6 | Delta CamSol score: 0.188

Stability Ranking: 3 | FoldX DDG: -13.763 kcal/mol

Mutation Score: 2.015

> Model_13 chain A Optimized_1bli_Repair | A45P G81E N96R T338P T341P S356R N473P G474P

LNGTLMQYFEWYMPNDGQHWKRLQNDSAYLAEHGITAVWIPP**P**YKGTSQADVGYGAYDLYDLGEFHQKGTVRTKYGTK**E**ELQSAIKSLHSRDI**R**VYGDVVINHKGGADATEDVTAVEVDPA

DRNRVISGEHLIKAWTHFHFPGRGSTYSDFKWHWYHFDGTDWDESRKLNRIYKFQGKAWDWEVSNEFGNYDYLMYADIDYDHPDVAAEIKRWGTWYANELQLDGFRLDAVKHIKFSFLRD

WVNHVREKTGKEMFTVAEYWSYDLGALENYLNKTNFNHSVFDVPLHYQFHAASTQGGGYDMRKLLNGTVVSKHPLKSVTFVDNHDTQPGQSLES**P**VQ**P**WFKPLAYAFILTRE**R**GYPQVFY

GDMYGTKGDSQREIPALKHKIEPILKARKQYAYGAQHDYFDHHDIVGWTREGDSSVANSGLAALITDGPGGAKRMYVGRQNAGETWHDITGNRSEPVVINSAGWGEFHV**PP**GSVSIYVQR

#### 6 Simultaneous Mutations:

**Model Name: Model_20**

Mutations by chain

Chain A : A45P,G81E,T338P,T341P,N473P,G474P

Solubility Ranking: 7 | Delta CamSol score: 0.151

Stability Ranking: 5 | FoldX DDG: -11.064 kcal/mol

Mutation Score: 1.61

> Model_20 chain A Optimized_1bli_Repair | A45P G81E T338P T341P N473P G474P

LNGTLMQYFEWYMPNDGQHWKRLQNDSAYLAEHGITAVWIPP**P**YKGTSQADVGYGAYDLYDLGEFHQKGTVRTKYGTK**E**ELQSAIKSLHSRDINVYGDVVINHKGGADATEDVTAVEVDPA

DRNRVISGEHLIKAWTHFHFPGRGSTYSDFKWHWYHFDGTDWDESRKLNRIYKFQGKAWDWEVSNEFGNYDYLMYADIDYDHPDVAAEIKRWGTWYANELQLDGFRLDAVKHIKFSFLRD

WVNHVREKTGKEMFTVAEYWSYDLGALENYLNKTNFNHSVFDVPLHYQFHAASTQGGGYDMRKLLNGTVVSKHPLKSVTFVDNHDTQPGQSLES**P**VQ**P**WFKPLAYAFILTRESGYPQVFY

GDMYGTKGDSQREIPALKHKIEPILKARKQYAYGAQHDYFDHHDIVGWTREGDSSVANSGLAALITDGPGGAKRMYVGRQNAGETWHDITGNRSEPVVINSAGWGEFHV**PP**GSVSIYVQR

**Model Name: Model_22**

Mutations by chain

Chain A : A45P,G81E,Q224G,T338P,T341P,G474P

Solubility Ranking: 8 | Delta CamSol score: 0.126

Stability Ranking: 5 | FoldX DDG: -11.271 kcal/mol

Mutation Score: 1.603

> Model_22 chain A Optimized_1bli_Repair | A45P G81E Q224G T338P T341P G474P

LNGTLMQYFEWYMPNDGQHWKRLQNDSAYLAEHGITAVWIPP**P**YKGTSQADVGYGAYDLYDLGEFHQKGTVRTKYGTK**E**ELQSAIKSLHSRDINVYGDVVINHKGGADATEDVTAVEVDPA

DRNRVISGEHLIKAWTHFHFPGRGSTYSDFKWHWYHFDGTDWDESRKLNRIYKFQGKAWDWEVSNEFGNYDYLMYADIDYDHPDVAAEIKRWGTWYANEL**G**LDGFRLDAVKHIKFSFLRD

WVNHVREKTGKEMFTVAEYWSYDLGALENYLNKTNFNHSVFDVPLHYQFHAASTQGGGYDMRKLLNGTVVSKHPLKSVTFVDNHDTQPGQSLES**P**VQ**P**WFKPLAYAFILTRESGYPQVFY

GDMYGTKGDSQREIPALKHKIEPILKARKQYAYGAQHDYFDHHDIVGWTREGDSSVANSGLAALITDGPGGAKRMYVGRQNAGETWHDITGNRSEPVVINSAGWGEFHVN**P**GSVSIYVQR

**Model Name: Model_21**

Mutations by chain

Chain A : A45P,G81E,T338P,T341P,S356R,G474P

Solubility Ranking: 7 | Delta CamSol score: 0.141

Stability Ranking: 5 | FoldX DDG: -11.604 kcal/mol

Mutation Score: 1.601

> Model_21 chain A Optimized_1bli_Repair | A45P G81E T338P T341P S356R G474P

LNGTLMQYFEWYMPNDGQHWKRLQNDSAYLAEHGITAVWIPP**P**YKGTSQADVGYGAYDLYDLGEFHQKGTVRTKYGTK**E**ELQSAIKSLHSRDINVYGDVVINHKGGADATEDVTAVEVDPA

DRNRVISGEHLIKAWTHFHFPGRGSTYSDFKWHWYHFDGTDWDESRKLNRIYKFQGKAWDWEVSNEFGNYDYLMYADIDYDHPDVAAEIKRWGTWYANELQLDGFRLDAVKHIKFSFLRD

WVNHVREKTGKEMFTVAEYWSYDLGALENYLNKTNFNHSVFDVPLHYQFHAASTQGGGYDMRKLLNGTVVSKHPLKSVTFVDNHDTQPGQSLES**P**VQ**P**WFKPLAYAFILTRE**R**GYPQVFY

GDMYGTKGDSQREIPALKHKIEPILKARKQYAYGAQHDYFDHHDIVGWTREGDSSVANSGLAALITDGPGGAKRMYVGRQNAGETWHDITGNRSEPVVINSAGWGEFHVN**P**GSVSIYVQR

#### 2 Simultaneous Mutations:

##### Model Name: Model_3

Mutations by chain

Chain A : A45P,G81E

Solubility Ranking: 11 | Delta CamSol score: 0.038

Stability Ranking: 10 | FoldX DDG: -4.413 kcal/mol

Mutation Score: 0.627

> Model_3 chain A Optimized_1bli_Repair | A45P G81E

LNGTLMQYFEWYMPNDGQHWKRLQNDSAYLAEHGITAVWIPP**P**YKGTSQADVGYGAYDLYDLGEFHQKGTVRTKYGTK**E**ELQSAIKSLHSRDINVYGDVVINHKGGADATEDVTAVEVDPA

DRNRVISGEHLIKAWTHFHFPGRGSTYSDFKWHWYHFDGTDWDESRKLNRIYKFQGKAWDWEVSNEFGNYDYLMYADIDYDHPDVAAEIKRWGTWYANELQLDGFRLDAVKHIKFSFLRD

WVNHVREKTGKEMFTVAEYWSYDLGALENYLNKTNFNHSVFDVPLHYQFHAASTQGGGYDMRKLLNGTVVSKHPLKSVTFVDNHDTQPGQSLESTVQTWFKPLAYAFILTRESGYPQVFY

GDMYGTKGDSQREIPALKHKIEPILKARKQYAYGAQHDYFDHHDIVGWTREGDSSVANSGLAALITDGPGGAKRMYVGRQNAGETWHDITGNRSEPVVINSAGWGEFHVNGGSVSIYVQR

##### Model Name: Model_15

Mutations by chain

Chain A : A45P,G474P

Solubility Ranking: 12 | Delta CamSol score: 0.023

Stability Ranking: 10 | FoldX DDG: -4.176 kcal/mol

Mutation Score: 0.579

> Model_15 chain A Optimized_1bli_Repair | A45P G474P

LNGTLMQYFEWYMPNDGQHWKRLQNDSAYLAEHGITAVWIPP**P**YKGTSQADVGYGAYDLYDLGEFHQKGTVRTKYGTKGELQSAIKSLHSRDINVYGDVVINHKGGADATEDVTAVEVDPA

DRNRVISGEHLIKAWTHFHFPGRGSTYSDFKWHWYHFDGTDWDESRKLNRIYKFQGKAWDWEVSNEFGNYDYLMYADIDYDHPDVAAEIKRWGTWYANELQLDGFRLDAVKHIKFSFLRD

WVNHVREKTGKEMFTVAEYWSYDLGALENYLNKTNFNHSVFDVPLHYQFHAASTQGGGYDMRKLLNGTVVSKHPLKSVTFVDNHDTQPGQSLESTVQTWFKPLAYAFILTRESGYPQVFY

GDMYGTKGDSQREIPALKHKIEPILKARKQYAYGAQHDYFDHHDIVGWTREGDSSVANSGLAALITDGPGGAKRMYVGRQNAGETWHDITGNRSEPVVINSAGWGEFHVN**P**GSVSIYVQR

##### Model Name: Model_8

Mutations by chain

Chain A : G81E,G474P

Solubility Ranking: 11 | Delta CamSol score: 0.029

Stability Ranking: 10 | FoldX DDG: -4.144 kcal/mol

Mutation Score: 0.572

> Model_8 chain A Optimized_1bli_Repair | G81E G474P

LNGTLMQYFEWYMPNDGQHWKRLQNDSAYLAEHGITAVWIPPAYKGTSQADVGYGAYDLYDLGEFHQKGTVRTKYGTK**E**ELQSAIKSLHSRDINVYGDVVINHKGGADATEDVTAVEVDPA

DRNRVISGEHLIKAWTHFHFPGRGSTYSDFKWHWYHFDGTDWDESRKLNRIYKFQGKAWDWEVSNEFGNYDYLMYADIDYDHPDVAAEIKRWGTWYANELQLDGFRLDAVKHIKFSFLRD

WVNHVREKTGKEMFTVAEYWSYDLGALENYLNKTNFNHSVFDVPLHYQFHAASTQGGGYDMRKLLNGTVVSKHPLKSVTFVDNHDTQPGQSLESTVQTWFKPLAYAFILTRESGYPQVFY

GDMYGTKGDSQREIPALKHKIEPILKARKQYAYGAQHDYFDHHDIVGWTREGDSSVANSGLAALITDGPGGAKRMYVGRQNAGETWHDITGNRSEPVVINSAGWGEFHVN**P**GSVSIYVQR

#### 1 Single Mutation:

##### Model Name: Model_6

Mutations by chain

Chain A : A45P

Solubility Ranking: 12 | Delta CamSol score: 0.016

Stability Ranking: 11 | FoldX DDG: -2.222 kcal/mol

Mutation Score: 0.317

> Model_6 chain A Optimized_1bli_Repair | A45P

LNGTLMQYFEWYMPNDGQHWKRLQNDSAYLAEHGITAVWIPPPYKGTSQADVGYGAYDLYDLGEFHQKGTVRTKYGTKGELQSAIKSLHSRDINVYGDVVINHKGGADATEDVTAVEVDPA

DRNRVISGEHLIKAWTHFHFPGRGSTYSDFKWHWYHFDGTDWDESRKLNRIYKFQGKAWDWEVSNEFGNYDYLMYADIDYDHPDVAAEIKRWGTWYANELQLDGFRLDAVKHIKFSFLRD

WVNHVREKTGKEMFTVAEYWSYDLGALENYLNKTNFNHSVFDVPLHYQFHAASTQGGGYDMRKLLNGTVVSKHPLKSVTFVDNHDTQPGQSLESTVQTWFKPLAYAFILTRESGYPQVFY

GDMYGTKGDSQREIPALKHKIEPILKARKQYAYGAQHDYFDHHDIVGWTREGDSSVANSGLAALITDGPGGAKRMYVGRQNAGETWHDITGNRSEPVVINSAGWGEFHVNGGSVSIYVQR

**Model Name: Model_17**

Mutations by chain

Chain A : G81E

Solubility Ranking: 12 | Delta CamSol score: 0.022

Stability Ranking: 11 | FoldX DDG: -2.19 kcal/mol

Mutation Score: 0.31

> Model_17 chain A Optimized_1bli_Repair | G81E

LNGTLMQYFEWYMPNDGQHWKRLQNDSAYLAEHGITAVWIPPAYKGTSQADVGYGAYDLYDLGEFHQKGTVRTKYGTKEELQSAIKSLHSRDINVYGDVVINHKGGADATEDVTAVEVDPA

DRNRVISGEHLIKAWTHFHFPGRGSTYSDFKWHWYHFDGTDWDESRKLNRIYKFQGKAWDWEVSNEFGNYDYLMYADIDYDHPDVAAEIKRWGTWYANELQLDGFRLDAVKHIKFSFLRD

WVNHVREKTGKEMFTVAEYWSYDLGALENYLNKTNFNHSVFDVPLHYQFHAASTQGGGYDMRKLLNGTVVSKHPLKSVTFVDNHDTQPGQSLESTVQTWFKPLAYAFILTRESGYPQVFY

GDMYGTKGDSQREIPALKHKIEPILKARKQYAYGAQHDYFDHHDIVGWTREGDSSVANSGLAALITDGPGGAKRMYVGRQNAGETWHDITGNRSEPVVINSAGWGEFHVNGGSVSIYVQR

**Model Name: Model_1**

Mutations by chain

Chain A : G474P

Solubility Ranking: 12 | Delta CamSol score: 0.007

Stability Ranking: 11 | FoldX DDG: -1.953 kcal/mol

Mutation Score: 0.262

> Model_1 chain A Optimized_1bli_Repair | G474P

LNGTLMQYFEWYMPNDGQHWKRLQNDSAYLAEHGITAVWIPPAYKGTSQADVGYGAYDLYDLGEFHQKGTVRTKYGTKGELQSAIKSLHSRDINVYGDVVINHKGGADATEDVTAVEVDPA

DRNRVISGEHLIKAWTHFHFPGRGSTYSDFKWHWYHFDGTDWDESRKLNRIYKFQGKAWDWEVSNEFGNYDYLMYADIDYDHPDVAAEIKRWGTWYANELQLDGFRLDAVKHIKFSFLRD

WVNHVREKTGKEMFTVAEYWSYDLGALENYLNKTNFNHSVFDVPLHYQFHAASTQGGGYDMRKLLNGTVVSKHPLKSVTFVDNHDTQPGQSLESTVQTWFKPLAYAFILTRESGYPQVFY

GDMYGTKGDSQREIPALKHKIEPILKARKQYAYGAQHDYFDHHDIVGWTREGDSSVANSGLAALITDGPGGAKRMYVGRQNAGETWHDITGNRSEPVVINSAGWGEFHVNPGSVSIYVQR

**Solubility and Stability enhancing combinations of mutations**

In the second part of the pipeline the single-point mutaions are combined to produce combination of mutations that are predicted to enhance both solubility and stability of the input protein. The combination process does not rely on running all the computational methods above mentioned for each possible combination since such a process would be too computationally costly. The single point mutants are combined by means of the sum of two of their main characteristics, namely DDG and Delta frequency. The CamSol score is then computed for each combination. Owing to its high computation speed, re-running the CamSol method for each combination does not slow down the algorithm. Once the three metrics are collected the mutation score for the combinations is calculated. Combinations are being produced until the limit of simultaneously occurring mutations set by the user is reached. Of all combination groups only some are selected as "best" groups. The best combination groups are those ones whose maximum mutation score marks a change in the behavior of the increment of the mutation score thorughout the different combination grous. Such a change in behavior is interpreted as a point in the combination proces from which the contribution of an additional mutant is not as favorable as it has been until that point. For all those groups label as "best" up to 3 combinations are modeled, while for the other groups just one model is returned. In generating the protein model bearing the mutation combination the DDG path of its mutations is checked. For DDG path we mean the contribution that each single mutant in the combination gives, in terms of stability, to the protein. If in applying one mutation after the other we register a non negative DDG value the algorithm tries to correct for it by swapping the single point mutation under scrutiny with another taken from the list of the other mutation combinations of the same combination group and that happens at the same point in the DDG path. If no viable alternative is found after three retries the mutation is simply skipped.

**Best mutant combination groups identified for 1,2,6,8,9 simultaneous mutations in combination**

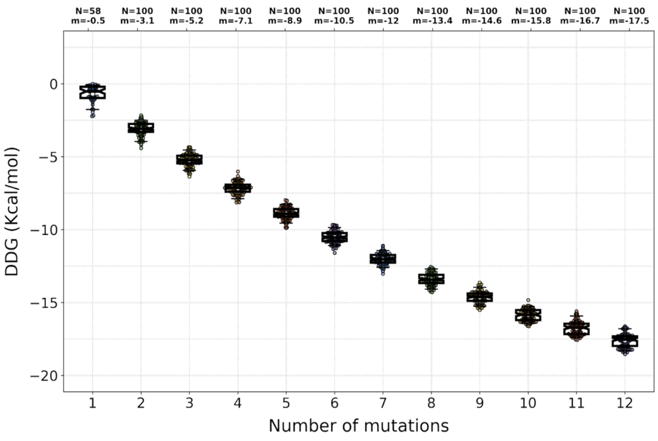


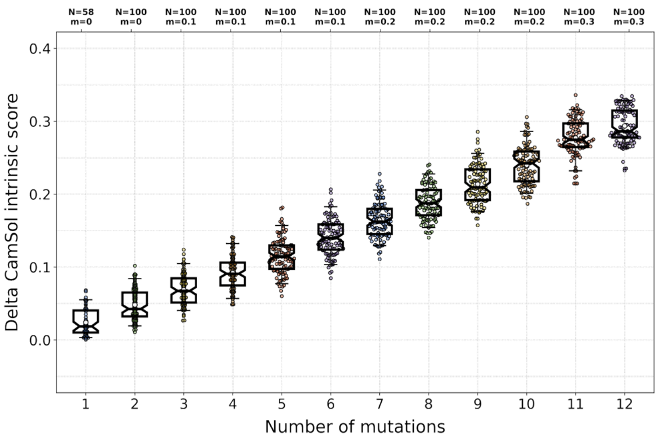


**Table with identified candidate mutation sites (105 sites)**

| Mutation site (seq index) | Mutation site (pdb number) | PSSM score > 0 | Delta PSSM score > 0 | site conservation | solvent exposure | solubilization potential | Problematic region+site score | region size | residue stru. corr. score | residue intrinsic score | wt residue frequency score | identified from |
|---|---|---|---|---|---|---|---|---|---|---|---|---|
| W156.A | WA157 | Y F P | | 0.403 | 0.318 | 0.124 | 6.671 | 9 | -0.687 | -2.155 | 4.703 | Solub. Seq. |
| W341.A | WA342 | R Q K N Y | | 0.204 | 0.5 | 0.106 | 6.012 | 13 | -0.522 | -1.07 | 2.432 | Solub. Seq. |
| Y264.A | YA265 | G S P D N W H | G P S D N H W | 0.189 | 1.0 | 0.074 | 3.31 | 3 | -1.254 | -1.304 | -0.012 | Solub. Seq. |
| T340.A | TA341 | D A E P Q N G | D E A P N Q G | 0.234 | 0.58 | 0.068 | 6.356 | 13 | -0.828 | -1.478 | -0.725 | Solub. Seq. |
| L266.A | LA267 | P N S A D R E | P N S D E A R | 0.189 | 0.144 | 0.064 | 2.612 | 3 | -0.105 | -0.678 | -0.468 | Solub. Seq. |
| W466.A | WA467 | S R K N T Q | S R K N | 0.243 | 0.495 | 0.06 | 2.24 | 3 | 0.05 | -0.278 | 0.323 | Solub. Seq. |
| V460.A | VA461 | T S E W K P | T S E K P W | 0.313 | 0.889 | 0.058 | 2.622 | 3 | 0.058 | -0.465 | 0.024 | Solub. Seq. |
| T337.A | TA338 | G D N S P E R K | D G N S P E R K | 0.15 | 0.22 | 0.049 | 5.309 | 13 | -0.075 | -0.432 | -1.235 | Solub. Seq. |
| Y157.A | YA158 | F | | 0.436 | 0.606 | 0.0 | 6.251 | 9 | -1.02 | -1.755 | 3.263 | Solub. Seq. |
| Y289.A | YA290 | W F D H N S Q E | W F | 0.162 | 0.228 | 0.182 | 6.63 | 16 | -0.394 | -1.531 | 1.474 | Solub. Seq. |
| W12.A | WA13 | R D Y | | 0.311 | 0.402 | 0.123 | 5.354 | 8 | -0.536 | -1.448 | 3.671 | Solub. Seq. |

| | | | | | | | | | | | | |
|---|---|---|---|---|---|---|---|---|---|---|---|---|
| F152.A | FA153 | Y P W D I G | | 0.248 | 0.228 | 0.119 | 5.545 | 9 | -0.048 | -1.024 | 2.739 | Solub. Seq. |
| T37.A | TA38 | N D | N | 0.422 | 0.301 | 0.077 | 5.865 | 6 | -0.408 | -1.67 | 2.308 | Solub. Seq. |
| Y357.A | YA358 | I V P T S | I P V S T | 0.324 | 0.242 | 0.068 | 2.643 | 3 | 0.024 | 0.026 | 0.248 | Solub. Seq. |
| M196.A | MA197 | G D F N Q S H | G D N Q S H F | 0.212 | 0.538 | 0.064 | 2.298 | 2 | -0.193 | -0.433 | -1.033 | Solub. Seq. |
| Y13.A | YA14 | S R D N Q A | S D R | 0.241 | 0.407 | 0.064 | 4.671 | 8 | -0.201 | -0.785 | 0.589 | Solub. Seq. |
| L195.A | LA196 | F S H G P V N E | G P S H E N F V | 0.129 | 0.477 | 0.061 | 3.247 | 2 | -0.632 | -1.37 | -0.605 | Solub. Seq. |
| F278.A | FA279 | G E N A S Q | G E N S A Q | 0.201 | 0.473 | 0.054 | 4.834 | 16 | 0.133 | 0.233 | -1.743 | Solub. Seq. |
| Y197.A | YA198 | L F Q G | L Q F G | 0.326 | 0.454 | 0.054 | 3.158 | 3 | -0.578 | -1.344 | -0.394 | Solub. Seq. & Conservation |
| H470.A | HA471 | T S N E Q K | T E N S K Q | 0.318 | 0.441 | 0.053 | 2.383 | 3 | 0.361 | -0.449 | -0.115 | Solub. Seq. & Conservation |
| L317.A | LA318 | S N Y Q W T D F E A H K | S N D E K Q Y T A H W F | 0.12 | 1.0 | 0.052 | 2.003 | 5 | 0.574 | 0.467 | -1.239 | Solub. Seq. |
| Y192.A | YA193 | W F H G | | 0.317 | 0.16 | 0.044 | 1.875 | 1 | -0.105 | -0.863 | 2.816 | Solub. Seq. |
| W183.A | WA184 | D N G Y | D | 0.244 | 0.835 | 0.043 | 1.869 | 1 | -0.665 | -0.824 | 1.436 | Solub. Seq. |
| T148.A | TA149 | P K Q N E R D | P K E R Q N D | 0.27 | 0.582 | 0.042 | 3.763 | 9 | 0.424 | 0.688 | -0.373 | Solub. Seq. & Conservation |
| F189.A | FA190 | V T R W Y L | R T V | 0.173 | 0.671 | 0.041 | 0.764 | 1 | 0.653 | 0.271 | 0.485 | Solub. Seq. |
| L133.A | LA134 | E P F N K T G D | E P N K D G T F | 0.164 | 0.814 | 0.037 | 3.45 | 7 | 0.271 | 0.181 | -1.469 | Solub. Seq. |
| Y366.A | YA367 | I F Q W L V | Q I F W | 0.257 | 0.393 | 0.03 | 1.881 | 3 | 0.203 | 0.357 | 0.77 | Solub. Seq. |
| W137.A | WA138 | Y F | | 0.565 | 0.601 | 0.021 | 5.003 | 7 | -0.517 | -1.337 | 5.428 | Solub. Seq. |
| T296.A | TA297 | N R S K A Q G | N R S K Q A G | 0.159 | 0.584 | 0.021 | 4.901 | 16 | -0.084 | 0.099 | -1.201 | Solub. Seq. |
| F256.A | FA257 | Y W I L V | | 0.301 | 0.222 | 0.016 | 3.715 | 4 | 0.148 | -0.606 | 2.674 | Solub. Seq. |
| Q223.A | QA224 | G D H | G D H | 0.556 | 0.953 | 0.015 | 0.707 | 2 | 0.462 | 0.549 | -0.364 | Solub. Seq. & Conservation |
| Y149.A | YA150 | F | | 0.583 | 0.4 | 0.0 | 4.885 | 9 | 0.035 | -0.389 | 3.928 | Solub. Seq. |
| L307.A | LA308 | F I W Y V | I F | 0.335 | 0.429 | 0.0 | 1.297 | 2 | 0.307 | 0.657 | 1.73 | Solub. Seq. |
| T428.A | TA429 | N | N | 0.876 | 0.0 | 0.023 | | | 0.0 | -0.065 | -3.2 | Conservation |
| S28.A | SA29 | L I A | L I A | 0.6 | 0.0 | 0.0 | | | 0.0 | 0.068 | -2.571 | Conservation |
| K105.A | KA106 | T V F S I R | T R S V F I | 0.309 | 0.0 | 0.004 | | | 0.0 | 0.422 | -2.941 | Conservation |

| | | | | | | | | | | | | |
|---|---|---|---|---|---|---|---|---|---|---|---|---|
| L2.A | LA3 | Q K R D E | Q K R D E | 0.292 | 1.0 | 0.028 | | | 0.033 | 0.009 | -2.664 | Conservatio |
| G98.A | GA99 | L F I V Q | L Q I F V | 0.406 | 0.0 | 0.0 | | | -0.0 | -1.157 | -2.215 | Conservatio |
| Q19.A | QA20 | G | G | 0.704 | 0.529 | 0.003 | | | 0.389 | 0.657 | -1.672 | Conservatio |
| R23.A | RA24 | G D A N | G D A N | 0.452 | 0.475 | 0.0 | | | 0.254 | 0.452 | -2.004 | Conservatio |
| A44.A | AA45 | I P V F | P I V F | 0.442 | 0.0 | 0.016 | | | 0.0 | 0.146 | -1.666 | Conservatio |
| R374.A | RA375 | W F Y | W F Y | 0.586 | 0.839 | 0.0 | | | 1.213 | 1.415 | -1.45 | Conservatio |
| A425.A | AA426 | V I | V I | 0.643 | 0.0 | 0.0 | | | -0.0 | -1.228 | -1.284 | Conservatio |
| N95.A | NA96 | K R A Y | K R A Y | 0.388 | 0.516 | 0.011 | | | 0.685 | -0.081 | -1.657 | Conservatio |
| A85.A | AA86 | L F | L F | 0.588 | 0.0 | 0.0 | | | 0.0 | -0.123 | -1.345 | Conservatio |
| V53.A | VA54 | W H S N | W H S N | 0.378 | 0.065 | 0.006 | | | -0.018 | -0.184 | -1.603 | Conservatio |
| H34.A | HA35 | L W A | L A W | 0.58 | 0.127 | 0.0 | | | 0.084 | -0.278 | -1.293 | Conservatio |
| P385.A | PA386 | K R N Q E D A | R K N E Q D A | 0.294 | 0.267 | 0.02 | | | 0.165 | 0.444 | -1.704 | Conservatio |
| D27.A | DA28 | K R H N Q E | K R H N Q E | 0.369 | 0.157 | 0.009 | | | 0.015 | 0.027 | -1.159 | Conservatio |
| L224.A | LA225 | I V F A | I V F A | 0.446 | 0.0 | 0.015 | | | -0.0 | -0.293 | -1.012 | Conservatio |
| M365.A | MA366 | E Y | E Y | 0.771 | 0.0 | 0.064 | | | -0.0 | -0.48 | -0.509 | Conservatio |
| H20.A | HA21 | D T W S N | D T W S N | 0.418 | 0.0 | 0.024 | | | 0.0 | 1.063 | -0.946 | Conservatio |
| G80.A | GA81 | E D Q A S | E D Q A S | 0.339 | 0.669 | 0.022 | | | 0.547 | 0.777 | -0.951 | Conservatio |
| T111.A | TA112 | H S N F | H S N F | 0.316 | 0.297 | 0.02 | | | 0.16 | 0.948 | -0.978 | Conservatio |
| A464.A | AA465 | D N G S | D N G S | 0.262 | 0.752 | 0.055 | | | -0.181 | -0.512 | -0.945 | Conservatio |
| S295.A | SA296 | F L I W V | F L I W V | 0.315 | 0.0 | 0.0 | | | -0.0 | -0.075 | -0.857 | Conservatio |
| T5.A | TA6 | V I A | V I A | 0.449 | 0.0 | 0.008 | | | -0.0 | -0.674 | -0.645 | Conservatio |
| E11.A | EA12 | P V I Y T | P V I T Y | 0.316 | 0.0 | 0.0 | | | -0.0 | -1.645 | -0.835 | Conservatio |
| M437.A | MA438 | V I F Y W L | V I F Y W L | 0.329 | 0.0 | 0.0 | | | -0.0 | 0.029 | -0.808 | Conservatio |
| A29.A | AA30 | D P E | D P E | 0.459 | 0.63 | 0.011 | | | 0.142 | -0.008 | -0.58 | Conservatio |
| A208.A | AA209 | Q R W I | R Q W I | 0.349 | 0.169 | 0.02 | | | 0.066 | 0.243 | -0.728 | Conservatio |
| V285.A | VA286 | F Y W D | F Y D W | 0.352 | 0.0 | 0.112 | | | -0.0 | -1.261 | -0.714 | Conservatio |
| R353.A | RA354 | T W F L S | T S W F L | 0.475 | 0.092 | 0.0 | | | 0.028 | 0.296 | -0.469 | Conservatio |
| D113.A | DA114 | W Y F L | W Y L F | 0.367 | 0.737 | 0.0 | | | 0.149 | 0.294 | -0.628 | Conservatio |
| A389.A | AA390 | I L F W V | L I F W V | 0.436 | 0.0 | 0.0 | | | 0.0 | 0.907 | -0.495 | Conservatio |

| | | | | | | | | | | | | |
|---|---|---|---|---|---|---|---|---|---|---|---|---|
| G215.A | GA216 | A L I V Y F | A L I V Y F | 0.254 | 0.0 | 0.0 | | | 0.0 | -0.499 | -0.752 | Conservatio |
| G409.A | GA410 | A V S I | A S V I | 0.443 | 0.0 | 0.0 | | | -0.0 | -0.585 | -0.459 | Conservatio |
| N125.A | NA126 | W Y F P I | W Y F P I | 0.388 | 0.931 | 0.0 | | | 0.411 | 0.721 | -0.478 | Conservatio |
| V117.A | VA118 | S Y A W Q | S A Y W Q | 0.27 | 0.099 | 0.016 | | | -0.048 | -0.625 | -0.645 | Conservatio |
| R172.A | RA173 | N G S Q P D | N G S P Q D | 0.272 | 0.632 | 0.0 | | | 0.616 | 0.678 | -0.576 | Conservatio |
| E384.A | EA385 | R Q K A S | R K Q A S | 0.332 | 0.307 | 0.01 | | | 0.249 | -0.051 | -0.48 | Conservatio |
| Y9.A | YA10 | I L W V A | I A L V W | 0.344 | 0.0 | 0.067 | | | -0.0 | -1.541 | -0.442 | Conservatio |
| A424.A | AA425 | L I V F | L I V F | 0.399 | 0.0 | 0.0 | | | -0.0 | -1.013 | -0.35 | Conservatio |
| L6.A | LA7 | I F W V | I F W V | 0.501 | 0.0 | 0.0 | | | -0.0 | -0.781 | -0.12 | Conservatio |
| T138.A | TA139 | F Y N P | N P F Y | 0.275 | 0.0 | 0.069 | | | -0.0 | -0.949 | -0.411 | Conservatio |
| N472.A | NA473 | P Q E K D A | P E D K Q A | 0.435 | 0.489 | 0.041 | | | 0.094 | -0.951 | -0.171 | Conservatio |
| L60.A | LA61 | Y F W H P | Y F H W P | 0.421 | 0.0 | 0.016 | | | -0.0 | -0.437 | -0.164 | Conservatio |
| T352.A | TA353 | F L W Y S A | F L S A W Y | 0.324 | 0.0 | 0.015 | | | 0.0 | -0.42 | -0.291 | Conservatio |
| T251.A | TA252 | A W S D P F E | A S D W P E F | 0.279 | 0.216 | 0.038 | | | 0.535 | 2.498 | -0.357 | Conservatio |
| Q373.A | QA374 | P D N | P D N | 0.348 | 1.0 | 0.038 | | | 1.355 | 1.334 | -0.167 | Conservatio |
| T257.A | TA258 | I F L W V | L I F W V | 0.324 | 0.0 | 0.0 | | | 0.0 | -0.464 | -0.198 | Conservatio |
| V244.A | VA245 | R I N T K Q W F | R I N K Q T W F | 0.275 | 0.0 | 0.033 | | | 0.0 | -0.526 | -0.23 | Conservatio |
| G467.A | GA468 | I V L F | L I V F | 0.256 | 0.0 | 0.0 | | | -0.0 | -0.741 | -0.258 | Conservatio |
| H280.A | HA281 | D Q N W S | D N Q S W | 0.287 | 0.0 | 0.049 | | | -0.0 | -1.265 | -0.184 | Conservatio |
| S416.A | SA417 | D G N E | D N G | 0.276 | 0.929 | 0.031 | | | 0.548 | 0.823 | 0.521 | Exposed Solub. |
| S355.A | SA356 | P R Q | P R Q | 0.384 | 0.295 | 0.026 | | | 0.475 | 0.86 | 0.067 | Exposed Solub. |
| G453.A | GA454 | S N D T | S N | 0.274 | 0.59 | 0.026 | | | 0.82 | 0.79 | 1.483 | Exposed Solub. |
| D242.A | DA243 | E Q W R A K Y N | E Q | 0.253 | 0.33 | 0.026 | | | 0.305 | -0.861 | 1.017 | Exposed Solub. |
| N220.A | NA221 | E R D K S T | E R D K | 0.269 | 0.493 | 0.022 | | | -0.37 | -0.867 | 0.738 | Exposed Solub. |
| Q83.A | QA84 | K R E | K R | 0.373 | 0.323 | 0.021 | | | 0.316 | -0.05 | 1.435 | Exposed Solub. |
| Q25.A | QA26 | I T A E | I T A | 0.305 | 0.315 | 0.017 | | | 0.775 | 1.488 | 0.235 | Exposed Solub. |

| | | | | | | | | | | | | |
|---|---|---|---|---|---|---|---|---|---|---|---|---|
| H67.A | HA68 | Y P D N G L | Y P D N G | 0.267 | 0.606 | 0.016 | | | 0.776 | 0.605 | 0.677 | Exposed Solub. |
| S91.A | SA92 | Q A K E R D N | Q K E A R D | 0.264 | 0.639 | 0.015 | | | 0.511 | 0.357 | 0.569 | Exposed Solub. |
| S84.A | SA85 | E N R Q A D K | E R N Q A | 0.26 | 0.39 | 0.013 | | | 0.291 | 0.232 | 0.572 | Exposed Solub. |
| S372.A | SA373 | P G N | P G | 0.462 | 0.234 | 0.01 | | | 0.73 | 2.166 | 0.405 | Exposed Solub. |
| N26.A | NA27 | Q E D S A R K | Q E D S | 0.252 | 0.686 | 0.01 | | | 0.964 | 0.908 | 0.959 | Exposed Solub. |
| K135.A | KA136 | P S R E Q D | P S | 0.29 | 0.335 | 0.009 | | | -0.197 | -1.084 | 1.073 | Exposed Solub. |
| G473.A | GA474 | P A S | P A | 0.523 | 0.437 | 0.007 | | | 0.121 | -0.917 | 0.664 | Exposed Solub. |
| K391.A | KA392 | R N Q | | 0.365 | 0.255 | 0.003 | | | 0.427 | 1.162 | 2.294 | Exposed Solub. |
| K253.A | KA254 | P E Q D | P | 0.42 | 0.329 | 0.002 | | | 0.402 | 0.579 | 0.475 | Exposed Solub. |
| V72.A | VA73 | I P L | I | 0.434 | 0.364 | 0.001 | | | 0.453 | 0.953 | 2.119 | Exposed Solub. |
| S129.A | SA130 | G H | G | 0.486 | 0.264 | 0.0 | | | 0.079 | 0.194 | 1.632 | Exposed Solub. |
| R482.A | RA483 | Q E D | E Q D | 0.495 | 0.433 | 0.0 | | | 0.466 | -0.674 | 0.802 | Exposed Solub. |
| I128.A | IA129 | Y W F D N | Y W F D | 0.313 | 0.387 | 0.0 | | | 0.326 | 0.48 | 0.097 | Exposed Solub. |
| K87.A | KA88 | D N E A R S Q | D | 0.302 | 0.663 | 0.0 | | | 0.872 | -0.152 | 1.332 | Exposed Solub. |

This table contains information on the identified mutation sites, selected on the basis of their contribution to the overall solubility and their solvent exposure, or on the basis of their conservation in the PSSM. The column 'Problematic region+site score' is an indicator (the higher the more aggregation-promoting) of how much a site is expected to contribute to the aggregation propensity accounting for both its own CamSol score and that of the broader sequence region that contains it. Sites identified from the structurally corrected profile are listed first, then those identified from the intrinsic solubility profile, which are also solvent-exposed but don't have a particularly negative structurally-corrected score, and finally those identified from the Conservation (see column 'identified from'). Mutations sites identified from solubility are ranked according to the column 'solubilization potential', which indicates how much the solubility can be improved by mutating that site according to those mutations allowed by the PSSM. As some problematic sites may be highly conserved, the 'solubilization potential' does not necessarily correlate with 'Problematic region+site score'. The 'sovlent exposure' is a score that ranges from 0 (non-exposed) to 1 (as exposed as in the context of a Gly-AminoAcidUnderScrutiny-Gly 3-peptide in an extended conformation), note that a solvent exposure of 0.5 would already offer the largest area to a potential aggregation partner. Sites identifide from 'Conservation' are those where the WT residue has negative log2(enrichment ratio) score from the input PSSM, and the site is relatively well conserved (conservation index > 0.25). The conservation index ranges in 0 to 1 and indicates how conserved a position in the alignment is (red line in the plot of the PSSM). Additional sites may be identified as sites that are solvent exposed, different from the consensus, and whose solubility can be improved with a substitution to a residue that has higher PSSM frequency than the WT residue (sites labelled as Exposed Solub.).

##### Results of the single-mutation scanning at all suitable sites

| mut_id_seqIndex | mut_id_pdb | Mutation Score | Delta CamSol intrinsic score | delta_frequency | DDG (kcal/mol) | Mutation type | CamSol intrinsic score | Mutation frequency |
|---|---|---|---|---|---|---|---|---|
| WT | WT | 0 | 0.0 | 0.0 | | n/a | 0.129 | n/a |
| LA2R | LA3R | 0.325 | 0.025 | 4.38 | -0.371 | Conservation | 0.155 | 1.716 |
| AA29P | AA30P | 0.28 | 0.003 | 3.004 | -0.961 | Conservation | 0.133 | 2.424 |
| AA44P | AA45P | 0.468 | 0.016 | 3.837 | -2.222 | Conservation | 0.145 | 2.171 |
| HA67P | HA68P | 0.195 | 0.003 | 1.105 | -1.255 | Exposed Solub. | 0.133 | 1.782 |
| HA67N | HA68N | 0.107 | 0.003 | 0.824 | -0.543 | Exposed Solub. | 0.133 | 1.501 |
| GA80E | GA81E | 0.442 | 0.022 | 3.356 | -2.19 | Conservation | 0.151 | 2.405 |

| | | | | | | | | |
|---|---|---|---|---|---|---|---|---|
| GA80D | GA81D | 0.276 | 0.019 | 2.775 | -0.906 | Conservation | 0.148 | 1.824 |
| QA83K | QA84K | 0.169 | 0.018 | 1.067 | -0.869 | Exposed Solub. | 0.147 | 2.502 |
| QA83R | QA84R | 0.166 | 0.021 | 0.58 | -1.108 | Exposed Solub. | 0.15 | 2.015 |
| SA84E | SA85E | 0.125 | 0.012 | 1.315 | -0.336 | Exposed Solub. | 0.142 | 1.887 |
| SA84R | SA85R | 0.107 | 0.013 | 0.848 | -0.431 | Exposed Solub. | 0.142 | 1.42 |
| SA84N | SA85N | 0.07 | 0.003 | 0.904 | -0.132 | Exposed Solub. | 0.132 | 1.476 |
| SA91K | SA92K | 0.079 | 0.012 | 0.815 | -0.185 | Exposed Solub. | 0.141 | 1.384 |
| SA91E | SA92E | 0.057 | 0.009 | 0.749 | -0.025 | Exposed Solub. | 0.139 | 1.318 |
| NA95K | NA96K | 0.341 | 0.006 | 4.191 | -0.835 | Conservation | 0.136 | 2.534 |
| NA95R | NA96R | 0.334 | 0.011 | 3.842 | -0.929 | Conservation | 0.14 | 2.185 |
| KA105R | KA106R | 0.258 | 0.004 | 3.068 | -0.702 | Conservation | 0.134 | 0.127 |
| NA125P | NA126P | 0.136 | 0.0 | 1.674 | -0.356 | Conservation | 0.13 | 1.196 |
| TA148K | TA149K | 0.174 | 0.039 | 1.36 | -0.535 | Solub. Seq. | 0.169 | 0.987 |
| TA148E | TA149E | 0.111 | 0.035 | 0.935 | -0.195 | Solub. Seq. | 0.164 | 0.562 |
| TA148R | TA149R | 0.112 | 0.042 | 0.731 | -0.257 | Solub. Seq. | 0.172 | 0.358 |
| TA148N | TA149N | 0.084 | 0.018 | 0.969 | -0.084 | Solub. Seq. | 0.147 | 0.596 |
| YA197G | YA198G | 0.211 | 0.054 | 0.817 | -1.081 | Solub. Seq. | 0.183 | 0.423 |
| AA208R | AA209R | 0.308 | 0.02 | 3.318 | -0.886 | Conservation | 0.15 | 2.59 |
| NA220R | NA221R | 0.181 | 0.02 | 0.864 | -1.097 | Exposed Solub. | 0.149 | 1.602 |
| NA220K | NA221K | 0.086 | 0.015 | 0.462 | -0.433 | Exposed Solub. | 0.145 | 1.2 |
| QA223G | QA224G | 0.358 | 0.011 | 3.377 | -1.436 | Solub. Seq. | 0.141 | 3.013 |
| DA242E | DA243E | 0.108 | 0.004 | 1.078 | -0.394 | Exposed Solub. | 0.133 | 2.095 |
| VA244I | VA245I | 0.083 | 0.005 | 1.099 | -0.119 | Conservation | 0.134 | 0.869 |
| TA251A | TA252A | 0.157 | 0.005 | 2.461 | -0.05 | Conservation | 0.134 | 2.104 |
| YA264P | YA265P | 0.146 | 0.067 | 1.136 | -0.114 | Solub. Stru. | 0.196 | 1.124 |
| TA296R | TA297R | 0.219 | 0.021 | 2.413 | -0.537 | Solub. Seq. | 0.15 | 1.212 |
| TA296K | TA297K | 0.22 | 0.018 | 1.92 | -0.864 | Solub. Seq. | 0.148 | 0.719 |
| LA317D | LA318D | 0.143 | 0.042 | 1.678 | -0.001 | Solub. Seq. | 0.172 | 0.439 |
| TA337P | TA338P | 0.321 | 0.025 | 2.011 | -1.76 | Solub. Stru. | 0.154 | 0.776 |
| TA337R | TA338R | 0.239 | 0.049 | 1.438 | -1.042 | Solub. Stru. | 0.178 | 0.203 |
| TA337K | TA338K | 0.157 | 0.044 | 1.297 | -0.35 | Solub. Stru. | 0.174 | 0.062 |
| TA340E | TA341E | 0.201 | 0.068 | 1.699 | -0.307 | Solub. Stru. | 0.197 | 0.974 |
| TA340P | TA341P | 0.311 | 0.048 | 1.541 | -1.709 | Solub. Stru. | 0.177 | 0.816 |
| TA340Q | TA341Q | 0.119 | 0.028 | 1.337 | -0.114 | Solub. Stru. | 0.157 | 0.612 |
| SA355P | SA356P | 0.28 | 0.007 | 3.135 | -0.849 | Exposed Solub. | 0.136 | 3.202 |
| SA355R | SA356R | 0.259 | 0.026 | 0.935 | -1.77 | Exposed Solub. | 0.156 | 1.002 |
| | | | | | | Exposed | | |

| | | | | | | | | |
|---|---|---|---|---|---|---|---|---|
| SA372P | SA373P | 0.203 | 0.01 | 3.017 | -0.124 | Solub. | 0.139 | 3.422 |
| QA373P | QA374P | 0.223 | 0.013 | 2.943 | -0.338 | Conservation | 0.142 | 2.776 |
| QA373N | QA374N | 0.084 | 0.012 | 0.972 | -0.133 | Conservation | 0.141 | 0.805 |
| SA416N | SA417N | 0.081 | 0.006 | 1.088 | -0.1 | Exposed Solub. | 0.135 | 1.609 |
| SA416G | SA417G | 0.089 | 0.002 | 1.15 | -0.179 | Exposed Solub. | 0.131 | 1.671 |
| VA460E | VA461E | 0.174 | 0.058 | 1.072 | -0.513 | Solub. Stru. | 0.188 | 1.096 |
| VA460K | VA461K | 0.173 | 0.055 | 0.394 | -0.944 | Solub. Stru. | 0.184 | 0.418 |
| VA460P | VA461P | 0.099 | 0.05 | 0.283 | -0.322 | Solub. Stru. | 0.179 | 0.307 |
| AA464D | AA465D | 0.265 | 0.055 | 3.177 | -0.201 | Conservation | 0.184 | 2.232 |
| AA464N | AA465N | 0.196 | 0.027 | 2.499 | -0.192 | Conservation | 0.156 | 1.554 |
| HA470E | HA471E | 0.157 | 0.053 | 0.938 | -0.477 | Solub. Seq. | 0.182 | 0.823 |
| HA470N | HA471N | 0.113 | 0.031 | 0.997 | -0.219 | Solub. Seq. | 0.16 | 0.882 |
| HA470S | HA471S | 0.186 | 0.017 | 1.103 | -1.027 | Solub. Seq. | 0.146 | 0.988 |
| HA470K | HA471K | 0.155 | 0.041 | 0.247 | -0.991 | Solub. Seq. | 0.171 | 0.132 |
| HA470Q | HA471Q | 0.14 | 0.01 | 0.644 | -0.915 | Solub. Seq. | 0.14 | 0.529 |
| NA472P | NA473P | 0.375 | 0.041 | 3.515 | -1.23 | Conservation | 0.17 | 3.344 |
| GA473P | GA474P | 0.375 | 0.007 | 2.863 | -1.953 | Exposed Solub. | 0.137 | 3.527 |

This table contains information on all possible single mutations at sites identified on the basis of their contribution to the overall solubility and their solvent exposure, as well as to their conservation in the MSA (see previous table). The column 'Mutation type' describes how a site has been identified: Solub. Stru. denotes that it was a site contributing to poor local solubility in the structurally corrected profile; Solub. seq. is the same but for the sequence-based intrinsic profile; Conservation indicates that the wild-type amino acid had log-likelihood<0 in the PSSM; Exposed Solub. are additional solvent-exposed sites where the WT residue is not strongly conserved (other residues with high PSSM score exist at that position) that may be mutated to further increase solubility (albeit unlike Solub. Seq. and Solub. Stru. these are typically not close to or within candidate aggregation hotspots). The column 'mut_id_seqIndex' contains candidate mutations numbered according to the index of the mutation site along the input sequence (from index 0), while in that 'mut_id_pdb' mutations are numbered according to the residue number in the input pdb file. This table has been filtered to contain only point-mutations predicted to increase both predicted solubility and stability - and if the strict-PSSM cutoff is selected as input - also PSSM frequency (see file 1blisingle_mutation_scanning.csv for a full table with all mutations tested, including those predicted destabilising).

##### Result of single-mutation scanning at all suitable sites

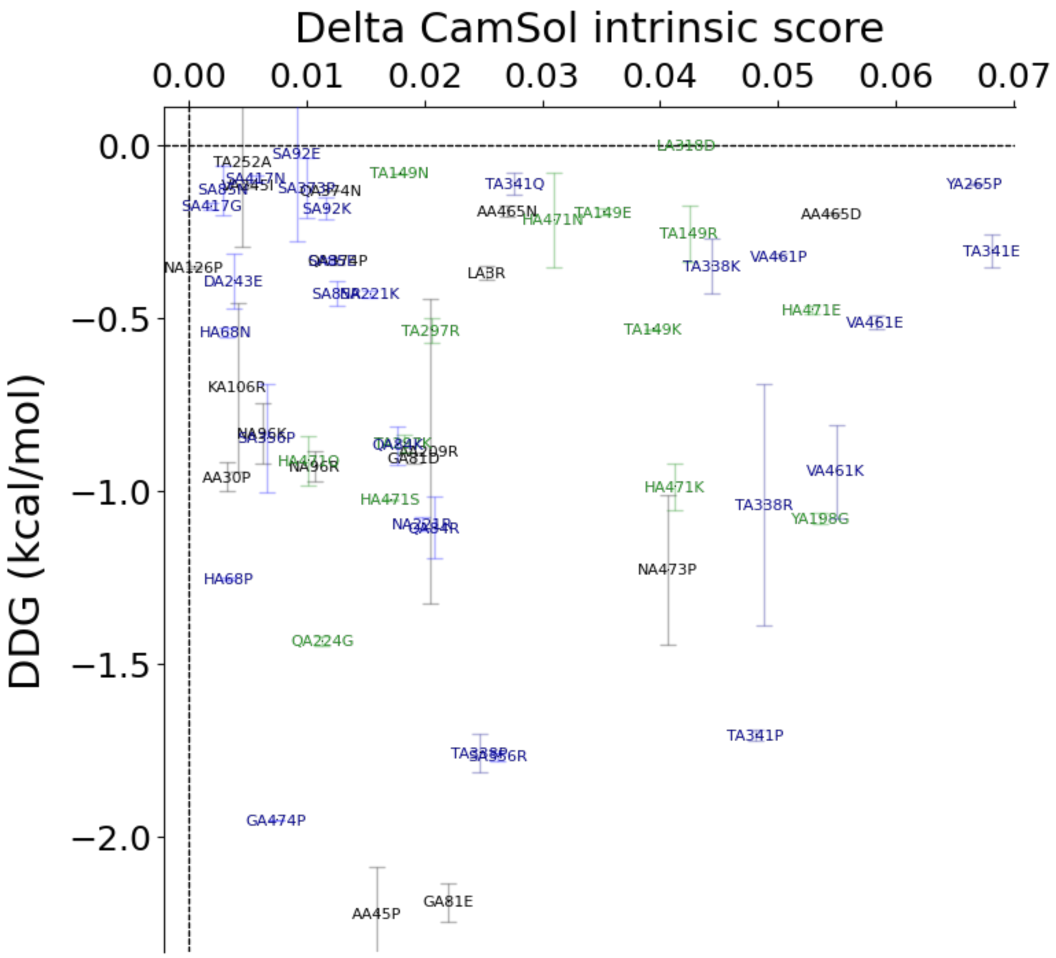
Delta CamSol intrinsic score
0.00
0.01
0.02
0.03
0.04
0.05
0.06
0.07
DDG (kcal/mol)
0.0
−0.5
−1.0
−1.5
−2.0
TA252A
SA92E
SA417N
SA85N
SA417G
SA373P
QA374N
SA92K
TA149N
TA341Q
LA318D
AA465N
HA471N
TA149E
AA465D
YA265P
TA149R
VA461P
TA338K
TA341E
NA126P
DA243E
LA3R
SA85P
NA221K
HA68N
TA297R
TA149K
HA471E
VA461E
KA106R
NA96K
SA356P
HA471Q
NA96R
AA30P
GA81D
AA209R
HA471K
TA338R
YA198G
VA461K
HA471S
NA221R
QA84R
HA68P
NA473P
QA224G
TA341P
TA338P
SA356R
GA474P
AA45P
GA81E

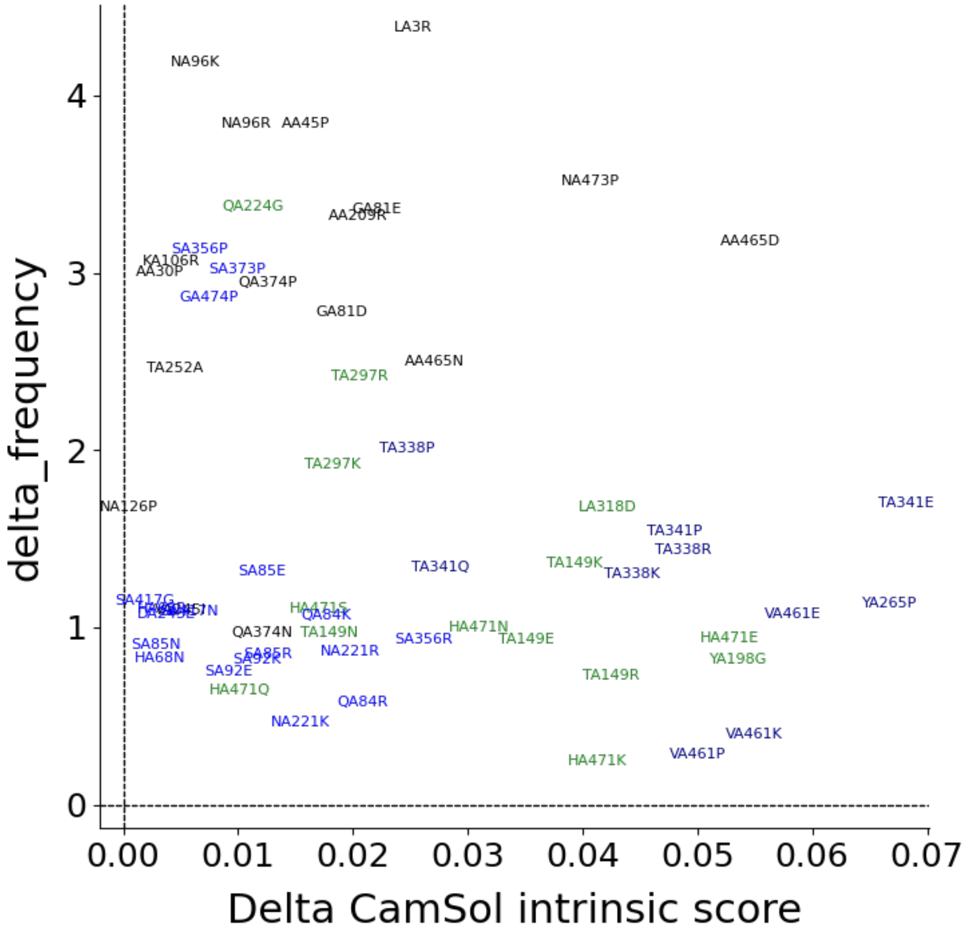


Plot of the data in the previous table, points are candidate mutations numbered according to the residue number within the input pdb file. The best mutations are those with large Delta CamSol score and large negative predicted DDG (if FoldX was run) and/or large positive Delta Frequency in MSA. Points in black correspond to mutation sites selected according to their conservation (see previous table), in green according to the sequence-based intrinsic solubility prediction, and in blue according to the structurally corrected one. Mutations at sites selected according to the solubility profiles (blue and green) are expected to have more impact on solubility than mutations at sites selected from conservation (black).

##### Mutation score of results of single-mutation scanning at all suitable sites (before normalisation)

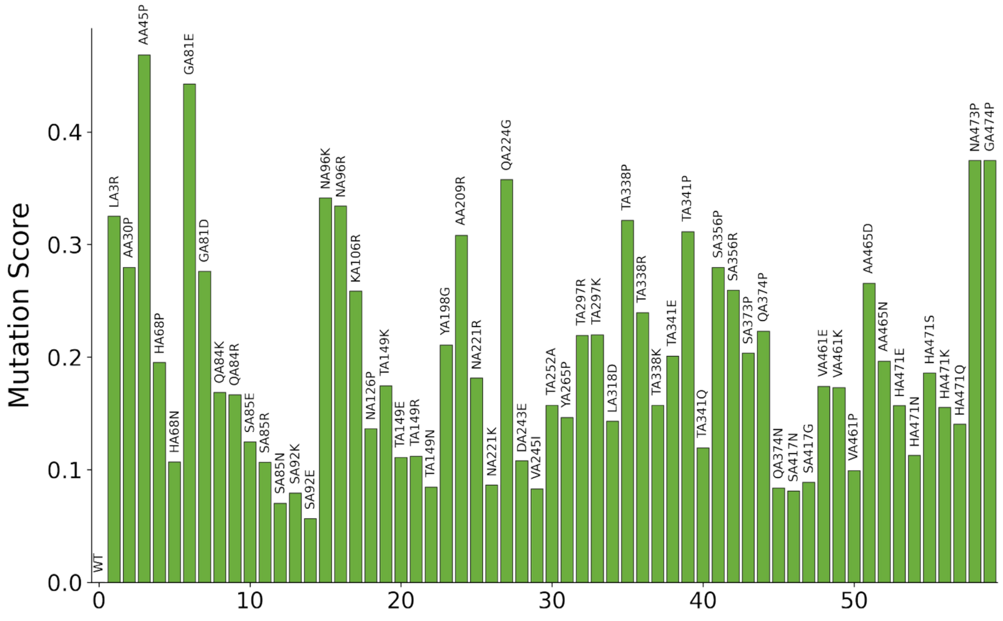


The plotted score is a rather arbitrary combination of delta solubility score, delta frequency, and predicted DDG (if FoldX was run). The highest this score the best the mutation. This score provides a nice visual ranking of mutations, but in practice one should refer to actual delta solubility score, delta frequency, and predicted DDG values to choose suitable mutations. This is for the single mutational scanning, then for combination of mutations, especially when across multiple chains, the mutation score is normalised usually resulting in a decreased contribution of the PSSM frequency (see paper).
