## Supplementary files 1 to 9 for "Automated optimisation of solubility and conformational stability of antibodies and proteins": SF4 Adalimumab_humanMSA_5mut report.pdf

#### PSSM used to pick candidate mutations

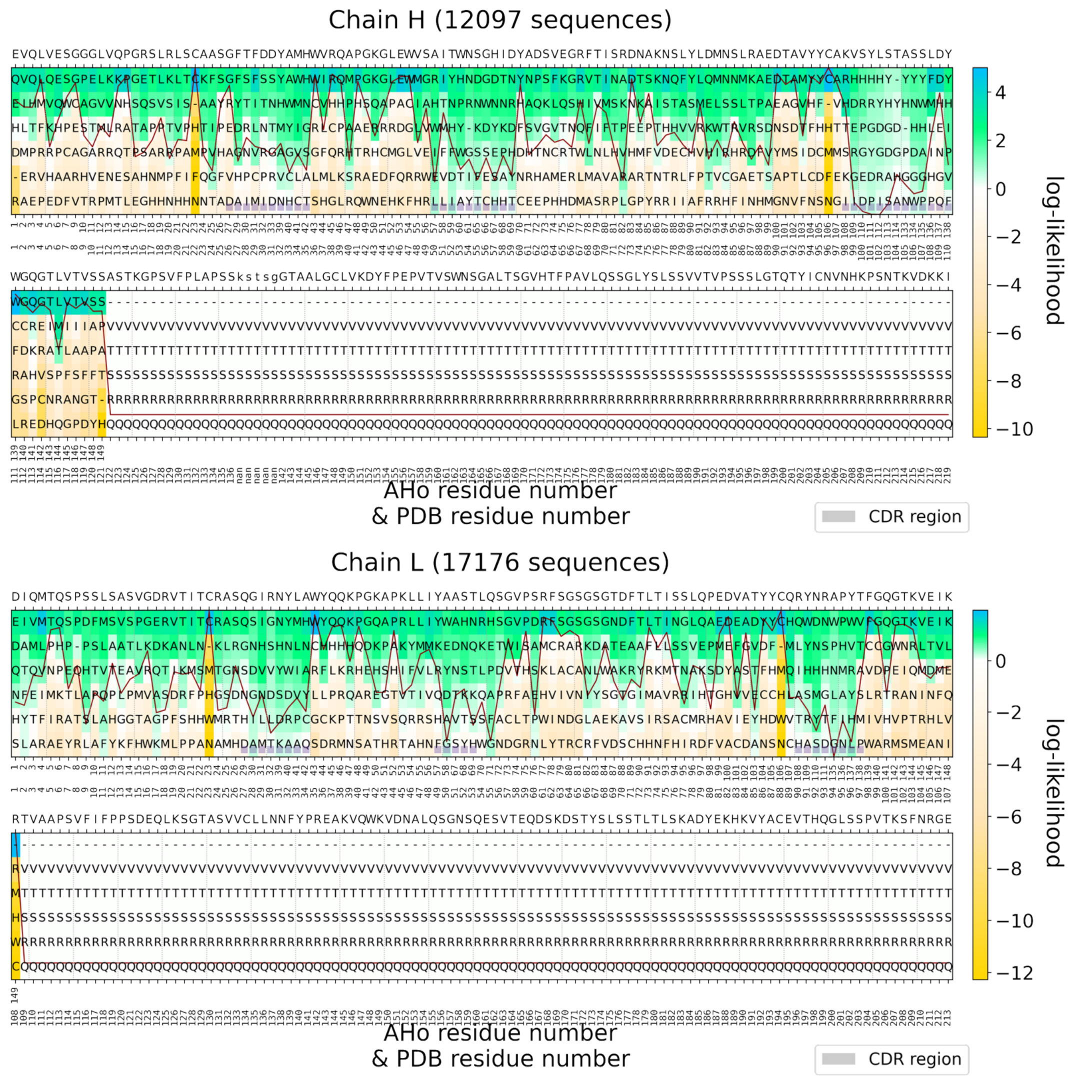


Position-specific scoring matrix (PSSM), as calculated from a multiple-sequence alignment (MSA) of similar sequences. The observed residue frequency (color-bar) is used to select candidate amino acid substitutions. The sequence above the panels is the wild-type (input) sequence as read from the alignment. The red line (if present) is the conservation index of each position (high means position highly conserved). Starting from the top panel: PSSM obtained from MSA of chain H containing 12097 Fv sequences (Fv region only). PSSM obtained from MSA of chain L containing 17176 Fv sequences (Fv region only).

#### CamSol intrinsic and structurally corrected profiles

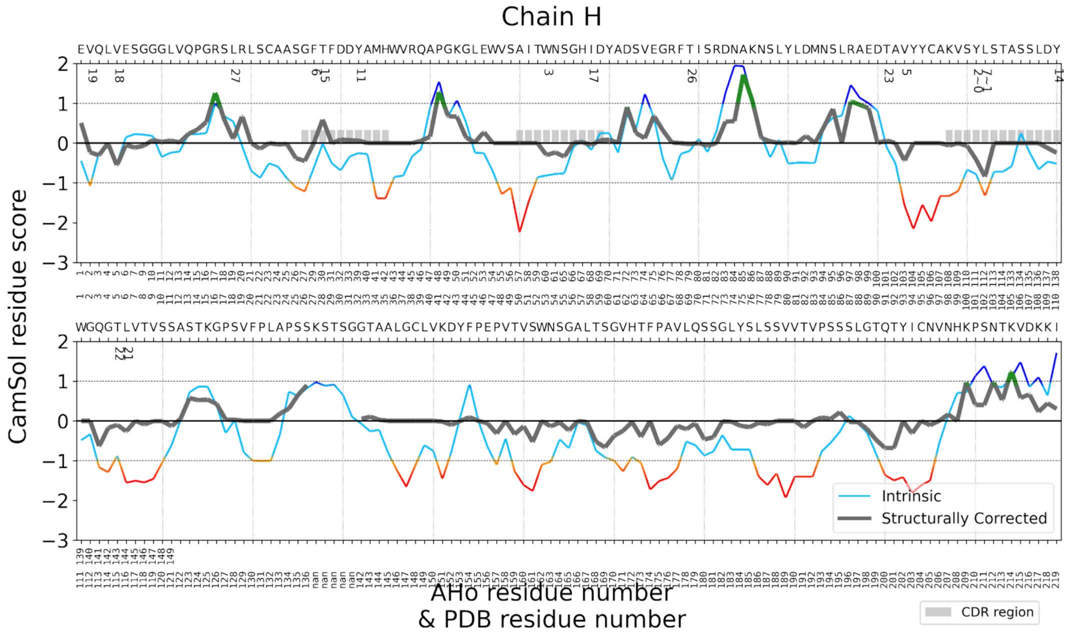

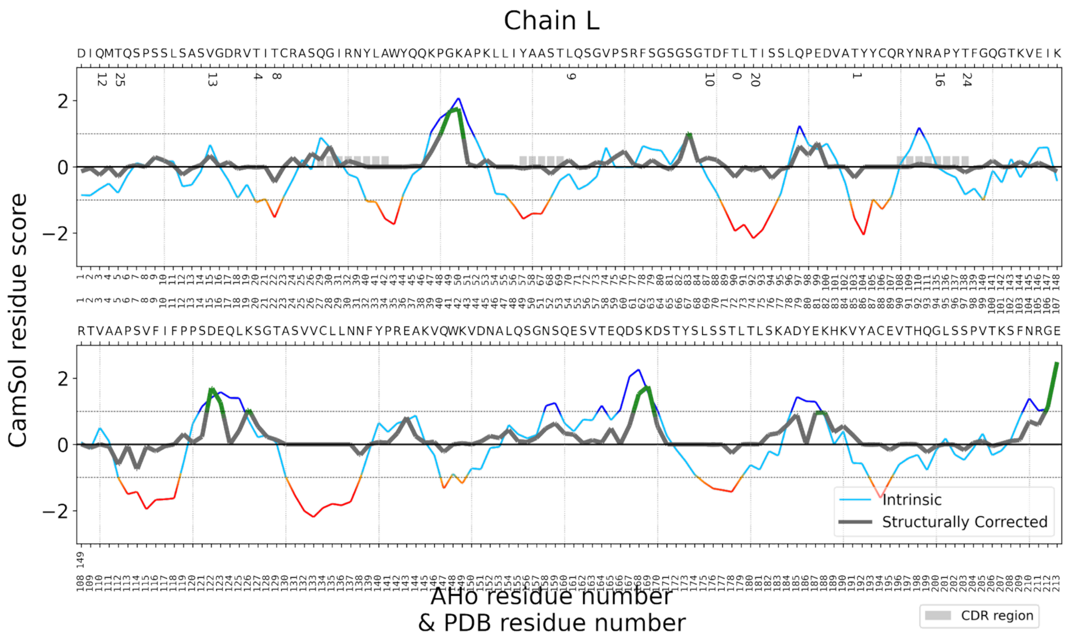


For each chain (see titles of the panels) the CamSol intrinsic profile is colour-coded red to blue, where red means aggregation-prone and blue aggregation-resistant. It is common for folded proteins to have large aggregation-prone regions in their intrinsic profile that typically drive the hydrophobic collapse during folding. The structurally corrected profile is color-coded in gray/green/magenta, regions of low negative scores (magenta) are potential aggregation hotspots, regions of high score (green) are solubility promoting. Numbers below the amino acid sequence at the top denote potential mutation sites, identified according to their contribution to the solubility, as well as their accessibility to the solvent.

### DESIGN PIPELINE RESULTS: Best Models

#### Table with identified best combinations of mutations

| Design Name | Number of Mutations | Mutations in Combination | Stability Rank | Solubility Rank | Theoretical PI | Mutation Score |
|---|---|---|---|---|---|---|
| model_7 | 1 | AH40P | 4 | 5 | 7.955 | 0.303 |
| model_6 | 1 | SH49G | 5 | 5 | 7.955 | 0.266 |
| model_3 | 1 | WH53P | 5 | 3 | 7.955 | 0.26 |
| model_2 | 5 | AH23K,AH40P,SH49G,WH53P,AL94P | 1 | 1 | 8.187 | 1.263 |
| model_1 | 4 | AH40P,SH49G,WH53P,AL94P | 1 | 2 | 7.955 | 1.081 |
| model_5 | 3 | AH40P,SH49G,WH53P | 2 | 2 | 7.955 | 0.828 |
| model_4 | 2 | AH40P,SH49G | 3 | 5 | 7.955 | 0.569 |
| WT | 0 | WT | 5 | 5 | 7.955 | 0 |

This table contains information on the identified mutation combinations, selected on the basis of their contribution to the overall solubility and stability of the protein. The column 'design name' identifies the pdb file bearing the mutations listed in the column 'mutations in combination'. Combinations belonging to the best combination groups (i.e., combinations containing 1 mutants) are listed first and sorted according to their mutation score. The remaining combinations, not identified as best mutation combinations, are listed afterwards and are sorted according to their mutation score. Mutation combinations are given a solubility and stability rank according to their delta CamSol score and DDG value respectively (i.e. they are binned starting from 1 corresponding the best predicted). The top ranking combination is usually one with the highest 'Number of Mutations'. However, because with each added mutation the chances of introducing a false positive increase, the best combination groups (top rows in the table) embody the best tradeoff between low number of mutations and high gain in solubility and stability. All combinations listed are expected to increase both solubility and stability (or one without compromising the other) and contain single point mutants that are enriched in the input PSSM.

#### 1 Single Mutation:

**Model Name: Model_7**

Mutations by chain

Chain H : A40P

Solubility Ranking: 5 | Delta CamSol score: 0.017

Stability Ranking: 4 | FoldX DDG: -2.782 kcal/mol

Mutation Score: 0.303

> Model_7 chain A Optimized_3wd5_Repair

RTPSDKPVAHVVANPQAEGQLQWLNDRANALLANGVELRDNQLVVPSEGLYLIYSQVLFKGQGCPSTHVLLTHTISRIAVSYQTKVNLLSAIKSPCQRETPEGAEAKPWYEPIYLGGVFQL

EKGDRLSAEINRPDYLDFAESGQVYFGIIAL

> Model_7 chain H Optimized_3wd5_Repair | A40P

EVQLVESGGGLVQPGRSLRLSCAASGFTFDDYAMHWVRQPPGKGLEWVSAITWNSGHIDYADSVEGRFTISRDNAKNSLYLDMNSLRAEDTAVYYCAKVSYLSTASSLDYWGQGTLVTVSS

ASTKGPSVFPLAPSS-----GTAALGCLVKDYFPEPVTVSWNSGALTSGVHTFPAVLQSSGLYSLSSVVTVPSSSLGTQTYICNVNHKPSNTKVDKKI

> Model_7 chain L Optimized_3wd5_Repair

DIQMTQSPSSLSASVGDRVTITCRASQGIRNYLAWYQQKPGKAPKLLIYAASTLQSGVPSRFSGSGSGTDFTLTISSLQPEDVATYYCQRYNRAPYTFGQGTKVEIKRTVAAPSVFIFPPS

DEQLKSGTASVVCLLNNFYPREAKVQWKVDNALQSGNSQESVTEQDSKDSTYSLSSTLTLSKADYEKHKVYACEVTHQGLSSPVTKSFNRGE

**Model Name: Model_6**

Mutations by chain

Chain H : S49G

Solubility Ranking: 5 | Delta CamSol score: 0.006

Stability Ranking: 5 | FoldX DDG: -1.694 kcal/mol

Mutation Score: 0.266

> Model_6 chain A Optimized_3wd5_Repair

RTPSDKPVAHVVANPQAEGQLQWLNDRANALLANGVELRDNQLVVPSEGLYLIYSQVLFKGQGCPSTHVLLTHTISRIAVSYQTKVNLLSAIKSPCQRETPEGAEAKPWYEPIYLGGVFQL

EKGDRLSAEINRPDYLDFAESGQVYFGIIAL

> Model_6 chain H Optimized_3wd5_Repair | S49G

EVQLVESGGGLVQPGRSLRLSCAASGFTFDDYAMHWVRQAPGKGLEWVGAITWNSGHIDYADSVEGRFTISRDNAKNSLYLDMNSLRAEDTAVYYCAKVSYLSTASSLDYWGQGTLVTVSS

ASTKGPSVFPLAPSS-----GTAALGCLVKDYFPEPVTVSWNSGALTSGVHTFPAVLQSSGLYSLSSVVTVPSSSLGTQTYICNVNHKPSNTKVDKKI

> Model_6 chain L Optimized_3wd5_Repair

DIQMTQSPSSLSASVGDRVTITCRASQGIRNYLAWYQQKPGKAPKLLIYAASTLQSGVPSRFSGSGSGTDFTLTISSLQPEDVATYYCQRYNRAPYTFGQGTKVEIKRTVAAPSVFIFPPS

DEQLKSGTASVVCLLNNFYPREAKVQWKVDNALQSGNSQESVTEQDSKDSTYSLSSTLTLSKADYEKHKVYACEVTHQGLSSPVTKSFNRGE

**Model Name: Model_3**

Mutations by chain

Chain H : W53P

Solubility Ranking: 3 | Delta CamSol score: 0.058

Stability Ranking: 5 | FoldX DDG: -1.91 kcal/mol

Mutation Score: 0.26

> Model_3 chain A Optimized_3wd5_Repair

RTPSDKPVAHVVANPQAEGQLQWLNDRANALLANGVELRDNQLVVPSEGLYLIYSQVLFKGQGCPSTHVLLTHTISRIAVSYQTKVNLLSAIKSPCQRETPEGAEAKPWYEPIYLGGVFQL

**Best mutant combination groups identified for 1 simultaneous mutations in combination**

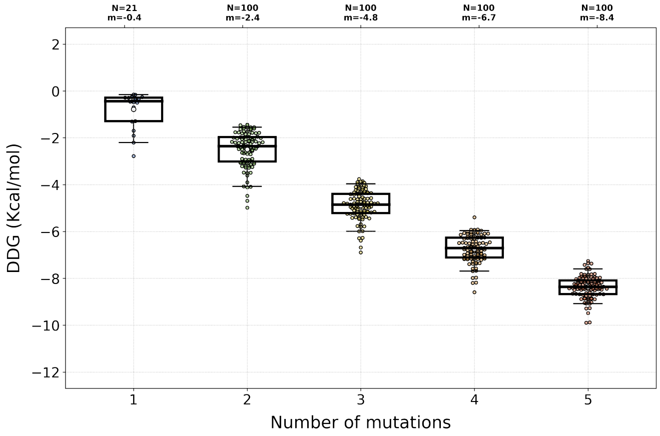


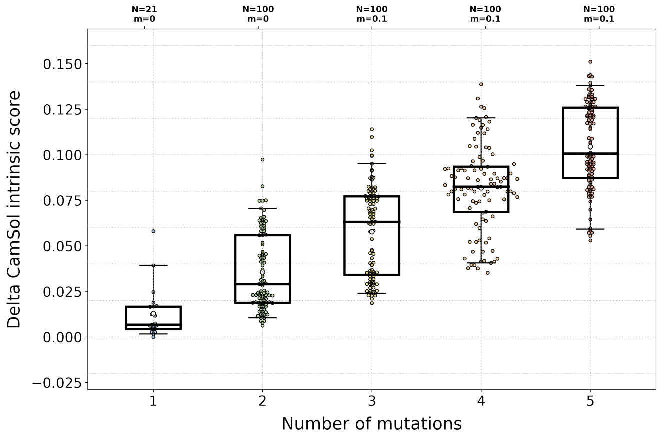


#### Table with identified candidate mutation sites (73 sites)

| Mutation site (seq index) | Mutation site (pdb number) | PSSM score > 0 | Delta PSSM score > 0 | site conservation | solvent exposure | solubilization potential | Problematic region+site score | region size | residue stru. corr. score | residue intrinsic score | wt residue frequency score | identified from |
|---|---|---|---|---|---|---|---|---|---|---|---|---|
| Y100.H | YH101 | H R G D P E F A N I | H R G | 0.122 | 0.623 | 0.063 | 2.702 | 4 | -0.41 | -0.779 | 0.84 | Solub. Seq. |
| L101.H | LH102 | H Y D G R I A N F S E P | G D H R Y A P E N S I F | 0.121 | 0.696 | 0.038 | 3.236 | 4 | -0.851 | -1.33 | -0.381 | Solub. Seq. |
| T71.L | TL72 | A Y I S | | 0.568 | 0.344 | 0.111 | 6.386 | 8 | -0.31 | -1.938 | 3.267 | Solub. Seq. |
| T84.L | TL85 | D V E Y A | D V | 0.438 | 0.256 | 0.07 | 4.716 | 5 | -0.331 | -1.558 | 1.684 | Solub. Seq. |
| W52.H | WH53 | H P Y T | H P Y | 0.395 | 0.716 | 0.058 | 4.33 | 9 | -0.3 | -0.814 | 2.061 | Solub. Seq. |
| T19.L | TL20 | N K R | | 0.646 | 0.606 | 0.055 | 3.51 | 4 | -0.013 | -1.074 | 3.434 | Solub. Seq. |
| V92.H | VH93 | T I | | 0.599 | 0.3 | 0.043 | 5.927 | 7 | -0.45 | -1.564 | 3.261 | Solub. Seq. |
| F26.H | FH27 | Y D G | | 0.446 | 0.195 | 0.041 | 2.647 | 3 | -0.001 | -0.621 | 3.187 | Solub. Seq. |
| T21.L | TL22 | N S | | 0.55 | 0.503 | 0.033 | 3.904 | 4 | -0.466 | -1.537 | 3.118 | Solub. Seq. |

| | | | | | | | | | | | | |
|---|---|---|---|---|---|---|---|---|---|---|---|---|
| L53.L | LL54 | R K Q S | R K | 0.484 | 0.403 | 0.032 | 3.453 | 7 | 0.204 | -0.209 | 1.8 | Solub. Seq. |
| T68.L | TL69 | N A | N | 0.605 | 0.452 | 0.027 | 4.695 | 8 | 0.276 | -0.314 | 3.25 | Solub. Seq. |
| Y31.H | YH32 | H A V | | 0.553 | 0.269 | 0.022 | 2.617 | 4 | 0.062 | -0.25 | 3.471 | Solub. Seq. |
| Q2.L | QL3 | V E | V | 0.64 | 0.867 | 0.019 | 2.039 | 2 | -0.249 | -0.667 | 1.261 | Solub. Seq. |
| V14.L | VL15 | P L | P L | 0.648 | 0.435 | 0.017 | 0.477 | 1 | 0.341 | 0.679 | 1.719 | Solub. Seq. |
| Y109.H | YH110 | H I P V F | | 0.425 | 0.473 | 0.015 | 4.356 | 8 | -0.242 | -0.517 | 2.963 | Solub. Seq. |
| T27.H | TH28 | S | S | 0.589 | 0.91 | 0.014 | 1.968 | 3 | 0.6 | -0.013 | 1.785 | Solub. Seq. |
| A93.L | AL94 | W P N L F G T S H | P N W L G T F S H | 0.164 | 0.17 | 0.012 | 0.967 | 1 | 0.009 | -0.014 | -0.331 | Solub. Seq. |
| I57.H | IH58 | T N K P A | N T K P A | 0.48 | 0.443 | 0.012 | 3.673 | 9 | -0.049 | -0.179 | -1.477 | Solub. Seq. & Conservation |
| V4.H | VH5 | Q K | Q | 0.533 | 0.784 | 0.008 | 1.752 | 1 | -0.566 | -0.511 | 2.77 | Solub. Seq. |
| V1.H | VH2 | I | | 0.775 | 0.283 | 0.003 | 2.317 | 1 | -0.23 | -1.076 | 3.675 | Solub. Seq. |
| T73.L | TL74 | | | 0.8 | 0.078 | 0.0 | 6.612 | 8 | -0.111 | -2.162 | 3.726 | Solub. Seq. |
| L115.H | LH116 | T | | 0.651 | 0.229 | 0.0 | 5.379 | 8 | -0.265 | -1.555 | 3.53 | Solub. Seq. |
| T114.H | TH115 | | | 0.894 | 0.131 | 0.0 | 4.67 | 8 | -0.102 | -0.874 | 3.67 | Solub. Seq. |
| T90.H | TH91 | | | 0.912 | 0.199 | 0.0 | 4.402 | 7 | 0.066 | -0.103 | 3.697 | Solub. Seq. |
| T96.L | TL97 | V | V | 0.595 | 0.28 | 0.0 | 1.81 | 1 | -0.17 | -0.839 | 3.007 | Solub. Seq. |
| T4.L | TL5 | | | 0.904 | 0.519 | 0.0 | 1.716 | 1 | -0.302 | -0.792 | 3.888 | Solub. Seq. |
| T68.H | TH69 | V | | 0.789 | 0.657 | 0.0 | 1.304 | 1 | -0.022 | -0.197 | 3.504 | Solub. Seq. |
| L17.H | LH18 | V | | 0.708 | 0.154 | 0.0 | 0.459 | 1 | 0.096 | 0.544 | 3.623 | Solub. Seq. |
| R107.L | RL108 | | | 1.019 | 0.36 | 0.0 | | | 0.005 | 0.069 | -8.654 | Conservation |
| S48.H | SH49 | G A | G A | 0.768 | 0.0 | 0.006 | | | -0.0 | -1.12 | -5.555 | Conservation |
| S77.H | SH78 | Q T H | Q T H | 0.62 | 0.076 | 0.0 | | | 0.043 | -0.368 | -5.718 | Conservation |
| A49.H | AH50 | R H W I E L Y | R E H I W L Y | 0.32 | 0.0 | 0.041 | | | -0.0 | -2.244 | -4.762 | Conservation |
| T51.H | TH52 | Y N H F D I | N Y D H F I | 0.538 | 0.0 | 0.069 | | | -0.0 | -0.856 | -3.943 | Conservation |
| R15.H | RH16 | E Q A | E Q A | 0.499 | 0.939 | 0.0 | | | 1.273 | 1.007 | -3.84 | Conservation |
| D61.H | DH62 | P Q V | P Q V | 0.54 | 1.0 | 0.0 | | | 0.91 | 0.724 | -3.588 | Conservation |
| D29.H | DH30 | S T N | S T N | 0.52 | 0.576 | 0.0 | | | 0.086 | -0.688 | -3.428 | Conservation |
| R89.L | RL90 | Q L I A T H | Q A L T H I | 0.513 | 0.0 | 0.0 | | | -0.0 | 0.104 | -2.968 | Conservation |
| G8.H | GH9 | P A | P A | 0.601 | 0.462 | 0.0 | | | 0.08 | 0.181 | -2.771 | Conservation |
| N73.H | NH74 | T K E | T K E | 0.679 | 0.344 | 0.014 | | | 0.551 | 1.946 | -2.131 | Conservation |
| D81.H | DH82 | Q E K H T | E Q K H T | 0.458 | 0.411 | 0.0 | | | 0.182 | -0.494 | -2.527 | Conservation |
| L78.H | LH79 | F A V | F A V | 0.546 | 0.0 | 0.003 | | | 0.0 | 0.098 | -2.192 | Conservation |
| A74.H | AH75 | S | S | 0.857 | 1.0 | 0.006 | | | 1.738 | 1.934 | -1.513 | Conservation |
| K97.H | KH98 | R H | R H | 0.77 | 0.041 | 0.012 | | | -0.059 | -1.326 | -1.66 | Conservation |
| Q12.H | QH13 | K | K | 0.902 | 0.686 | 0.007 | | | 0.231 | 0.224 | -1.046 | Conservation |
| V47.H | VH48 | I L | I L | 0.558 | 0.0 | 0.024 | | | -0.0 | -1.282 | -1.593 | Conservation |
| R92.L | RL93 | N S H G T D | N S H G D T | 0.466 | 0.064 | 0.0 | | | 0.041 | 0.685 | -1.366 | Conservation |

| | | | | | | | | | | | | |
|---|---|---|---|---|---|---|---|---|---|---|---|---|
| S106.H | SH107 | Y W H A G P | H A G Y P W | 0.262 | 0.0 | 0.004 | | | -0.0 | -0.242 | -1.423 | Conservatio |
| T103.H | TH104 | Y D G H A | D G H A Y | 0.364 | 0.0 | 0.028 | | | -0.0 | -0.719 | -1.116 | Conservatio |
| D30.H | DH31 | S N T G | S N T G | 0.487 | 0.769 | 0.0 | | | 0.072 | -0.339 | -0.58 | Conservatio |
| E64.H | EH65 | K Q T R | K Q R T | 0.578 | 0.879 | 0.004 | | | 0.689 | 0.637 | -0.311 | Conservatio |
| R29.L | RL30 | G S V D L T | G S D L T V | 0.303 | 0.276 | 0.0 | | | 0.184 | -0.223 | -0.595 | Conservatio |
| A104.H | AH105 | Y H P G N R | H Y P R G N | 0.302 | 0.0 | 0.025 | | | -0.0 | -0.577 | -0.485 | Conservatio |
| S76.L | SL77 | G N R | G | 0.473 | 0.331 | 0.044 | | | -0.07 | -0.8 | 2.2 | Exposed Solub. |
| Q78.L | QL79 | E K | | 0.672 | 0.285 | 0.043 | | | 0.631 | 1.261 | 3.648 | Exposed Solub. |
| A22.H | AH23 | K T | K | 0.48 | 0.375 | 0.039 | | | -0.032 | -0.592 | 2.206 | Exposed Solub. |
| N53.H | NH54 | R G I | | 0.457 | 0.462 | 0.026 | | | -0.233 | -0.776 | 2.46 | Exposed Solub. |
| S54.H | SH55 | D N K F T | D N K | 0.332 | 0.373 | 0.021 | | | -0.344 | -0.759 | 1.132 | Exposed Solub. |
| A87.H | AH88 | P S | | 0.542 | 0.519 | 0.017 | | | 0.966 | 1.138 | 3.303 | Exposed Solub. |
| A39.H | AH40 | P H S | P | 0.451 | 0.144 | 0.017 | | | 0.163 | 0.897 | 2.548 | Exposed Solub. |
| S75.L | SL76 | N | N | 0.647 | 0.399 | 0.017 | | | -0.35 | -1.369 | 2.779 | Exposed Solub. |
| S51.L | SL52 | H N K T | N H | 0.559 | 0.442 | 0.014 | | | -0.22 | -0.934 | 2.615 | Exposed Solub. |
| H56.H | HH57 | D N Y E S | D N E Y S | 0.327 | 0.154 | 0.014 | | | 0.026 | 0.072 | 0.537 | Exposed Solub. |
| Q54.L | QL55 | H E P A F G I V | H E P A G F I | 0.257 | 0.175 | 0.013 | | | -0.029 | -0.685 | 0.685 | Exposed Solub. |
| R86.H | RH87 | K T D | K T | 0.458 | 0.406 | 0.011 | | | 1.051 | 1.466 | 2.363 | Exposed Solub. |
| K44.L | KL45 | R I V Q | R | 0.531 | 0.579 | 0.007 | | | 0.214 | 0.127 | 3.711 | Exposed Solub. |
| S59.L | SL60 | D A E P N | D A | 0.407 | 0.732 | 0.007 | | | 0.469 | -0.003 | 1.589 | Exposed Solub. |
| Q99.L | QL100 | G P T | | 0.529 | 0.444 | 0.006 | | | 0.194 | -0.102 | 2.829 | Exposed Solub. |
| D0.L | DL1 | E Q N | E | 0.483 | 0.441 | 0.005 | | | -0.135 | -0.851 | 2.995 | Exposed Solub. |
| S84.H | SH85 | N | N | 0.686 | 0.502 | 0.004 | | | 0.866 | 0.638 | 2.813 | Exposed Solub. |
| S70.H | SH71 | N T | N | 0.51 | 0.329 | 0.003 | | | -0.066 | -0.229 | 2.303 | Exposed Solub. |
| D16.L | DL17 | E Q A G K | E | 0.422 | 0.406 | 0.003 | | | 0.149 | -0.412 | 2.725 | Exposed Solub. |
| S8.L | SL9 | D P H A | D P H A | 0.355 | 0.553 | 0.002 | | | 0.293 | 0.266 | 0.852 | Exposed Solub. |
| G27.L | GL28 | S N D I A | S N D | 0.395 | 0.465 | 0.001 | | | 0.627 | 0.588 | 0.888 | Exposed Solub. |

##### Results of the single-mutation scanning at all suitable sites

| mut_id_seqIndex | mut_id_pdb | Mutation Score | Delta CamSol intrinsic score | delta_frequency | DDG (kcal/mol) | Mutation type | CamSol intrinsic score | Mutation frequency | Chain A CamSol intrinsic score | Chain H CamSol intrinsic score | [illegible] |
|---|---|---|---|---|---|---|---|---|---|---|---|
| WT | WT | 0 | 0.0 | 0.0 | | n/a | -0.928 | n/a | 0.415 | -0.055 | [illegible] |
| QH12K | QH13K | 0.374 | 0.007 | 5.72 | -0.245 | Conservation | -0.921 | 4.674 | 0.415 | -0.044 | [illegible] |
| AH22K | AH23K | 0.245 | 0.039 | 1.248 | -1.304 | Exposed Solub. | -0.889 | 3.454 | 0.415 | 0.01 | [illegible] |
| AH39P | AH40P | 0.343 | 0.017 | 0.803 | -2.782 | Exposed Solub. | -0.911 | 3.35 | 0.415 | -0.027 | [illegible] |
| SH48G | SH49G | 0.713 | 0.006 | 8.952 | -1.694 | Conservation | -0.922 | 3.397 | 0.415 | -0.045 | [illegible] |
| WH52P | WH53P | 0.314 | 0.058 | 1.09 | -1.91 | Solub. Seq. | -0.87 | 3.15 | 0.415 | 0.042 | [illegible] |
| SH54K | SH55K | 0.201 | 0.019 | 0.895 | -1.288 | Exposed Solub. | -0.909 | 2.027 | 0.415 | -0.024 | [illegible] |
| HH56S | HH57S | 0.05 | 0.006 | 0.261 | -0.279 | Exposed Solub. | -0.922 | 0.798 | 0.415 | -0.045 | [illegible] |
| SH70N | SH71N | 0.055 | 0.003 | 0.442 | -0.257 | Exposed Solub. | -0.926 | 2.746 | 0.415 | -0.051 | [illegible] |
| AH74S | AH75S | 0.333 | 0.006 | 4.648 | -0.483 | Conservation | -0.922 | 3.135 | 0.415 | -0.045 | [illegible] |
| SH84N | SH85N | 0.058 | 0.004 | 0.127 | -0.46 | Exposed Solub. | -0.924 | 2.939 | 0.415 | -0.048 | [illegible] |
| KH97R | KH98R | 0.399 | 0.012 | 6.205 | -0.149 | Conservation | -0.916 | 4.546 | 0.415 | -0.036 | [illegible] |
| LH101R | LH102R | 0.116 | 0.025 | 0.996 | -0.311 | Solub. Stru. | -0.903 | 0.615 | 0.415 | -0.014 | [illegible] |
| TH103A | TH104A | 0.087 | 0.008 | 1.134 | -0.116 | Conservation | -0.92 | 0.017 | 0.415 | -0.042 | [illegible] |
| SH106P | SH107P | 0.168 | 0.004 | 2.005 | -0.431 | Conservation | -0.924 | 0.582 | 0.415 | -0.048 | [illegible] |
| SL8D | SL9D | 0.149 | 0.002 | 1.861 | -0.358 | Exposed Solub. | -0.927 | 2.712 | 0.415 | -0.055 | [illegible] |
| SL8P | SL9P | 0.159 | 0.0 | 1.826 | -0.495 | Exposed Solub. | -0.928 | 2.677 | 0.415 | -0.055 | [illegible] |
| VL14P | VL15P | 0.173 | 0.017 | 2.181 | -0.256 | Solub. Seq. | -0.911 | 3.899 | 0.415 | -0.055 | [illegible] |
| VL14L | VL15L | 0.042 | 0.007 | 0.04 | -0.319 | Solub. Seq. | -0.921 | 1.758 | 0.415 | -0.055 | [illegible] |
| DL16E | DL17E | 0.137 | 0.003 | 1.076 | -0.697 | Exposed Solub. | -0.926 | 3.802 | 0.415 | -0.055 | [illegible] |
| SL59D | SL60D | 0.144 | 0.005 | 1.833 | -0.29 | Exposed Solub. | -0.923 | 3.422 | 0.415 | -0.055 | [illegible] |
| SL75N | SL76N | 0.071 | 0.017 | 0.657 | -0.153 | Exposed Solub. | -0.912 | 3.436 | 0.415 | -0.055 | [illegible] |
| AL93P | AL94P | 0.345 | 0.012 | 1.876 | -2.202 | Solub. Seq. | -0.916 | 1.545 | 0.415 | -0.055 | [illegible] |

This table contains information on all possible single mutations at sites identified on the basis of their contribution to the overall solubility and their solvent exposure, as well as to their conservation in the MSA (see previous table). The column 'Mutation type' describes how a site has been identified: Solub. Stru. denotes that it was a site contributing to poor local solubility in the structurally corrected profile; Solub. seq. is the same but for the sequence-based intrinsic profile; Conservation indicates that the wild-type amino acid had log-likelihood<0 in the PSSM; Exposed Solub. are additional solvent-exposed sites where the WT residue is not strongly conserved (other residues with high PSSM score exist at that position) that may be mutated to further increase solubility (albeit unlike Solub. Seq. and Solub. Stru. these are typically not close to or within candidate aggregation hotspots). The column 'mut_id_seqIndex' contains candidate mutations numbered according to the index of the mutation site along the input sequence (from index 0), while in that 'mut_id_pdb' mutations are numbered according to the residue number in the input pdb file. This table has been filtered to contain only point-mutations predicted to increase both predicted solubility and stability - and if the strict-PSSM cutoff is selected as input - also PSSM frequency (see file 3wd5single_mutation_scanning.csv for a full table with all mutations tested, including those predicted destabilising).

##### Result of single-mutation scanning at all suitable sites

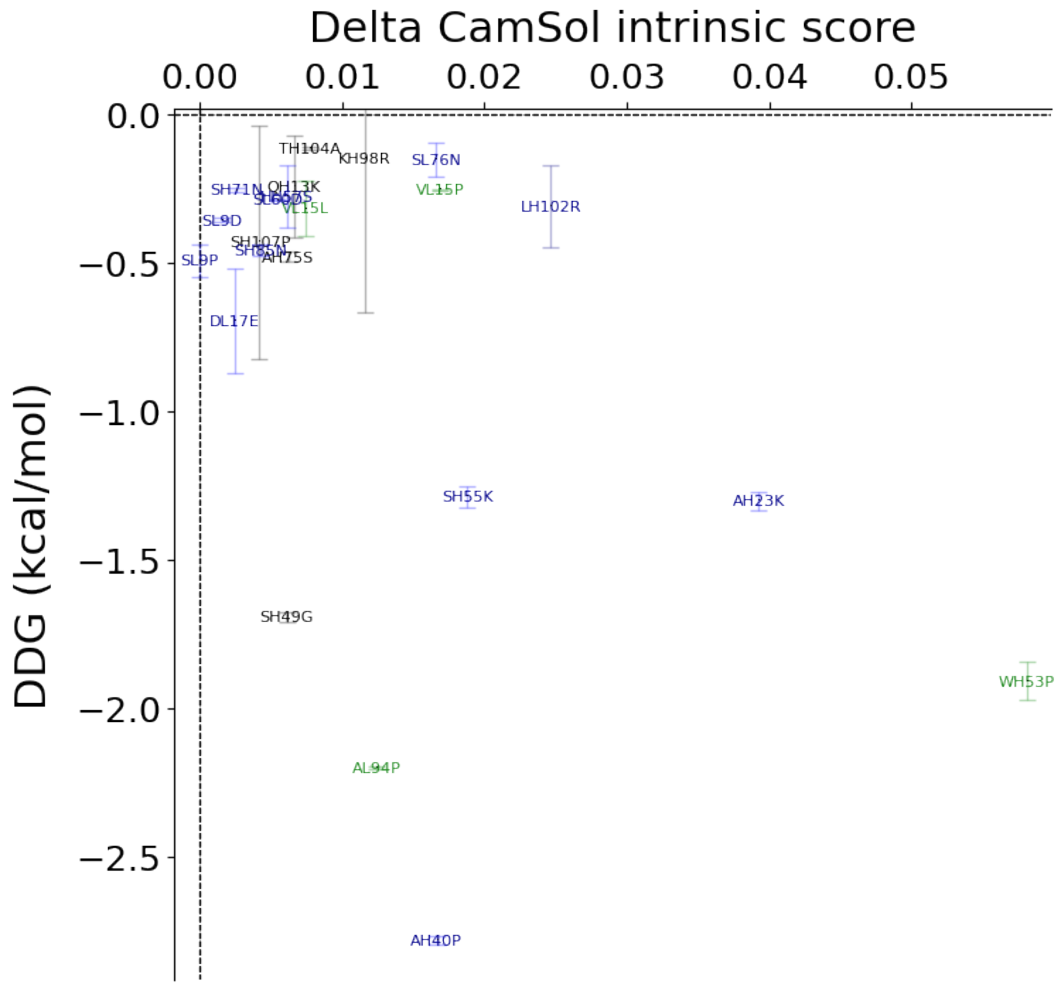

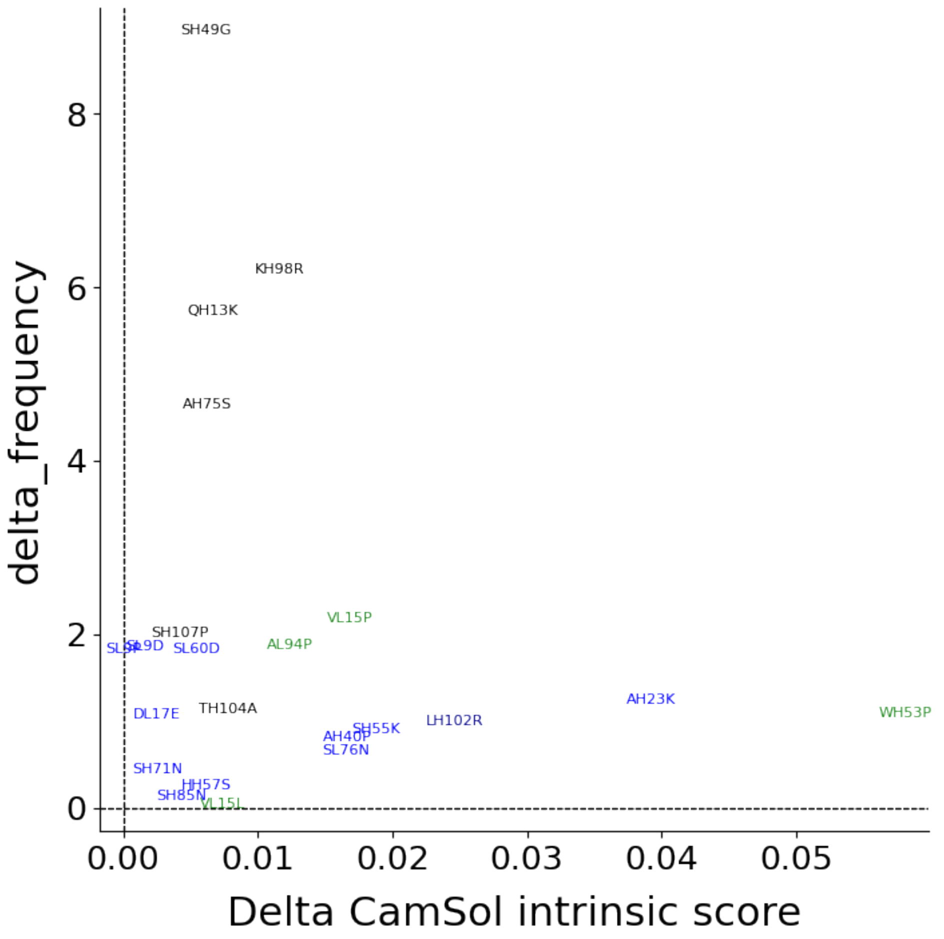


Plot of the data in the previous table, points are candidate mutations numbered according to the residue number within the input pdb file. The best mutations are those with large Delta CamSol score and large negative predicted DDG (if FoldX was run) and/or large positive Delta Frequency in MSA. Points in black correspond to mutation sites selected according to their conservation (see previous table), in green according to the sequence-based intrinsic solubility prediction, and in blue according to the structurally corrected one. Mutations at sites selected according to the solubility profiles (blue and green) are expected to have more impact on solubility than mutations at sites selected from conservation (black).

**Mutation score of results of single-mutation scanning at all suitable sites (before normalisation)**

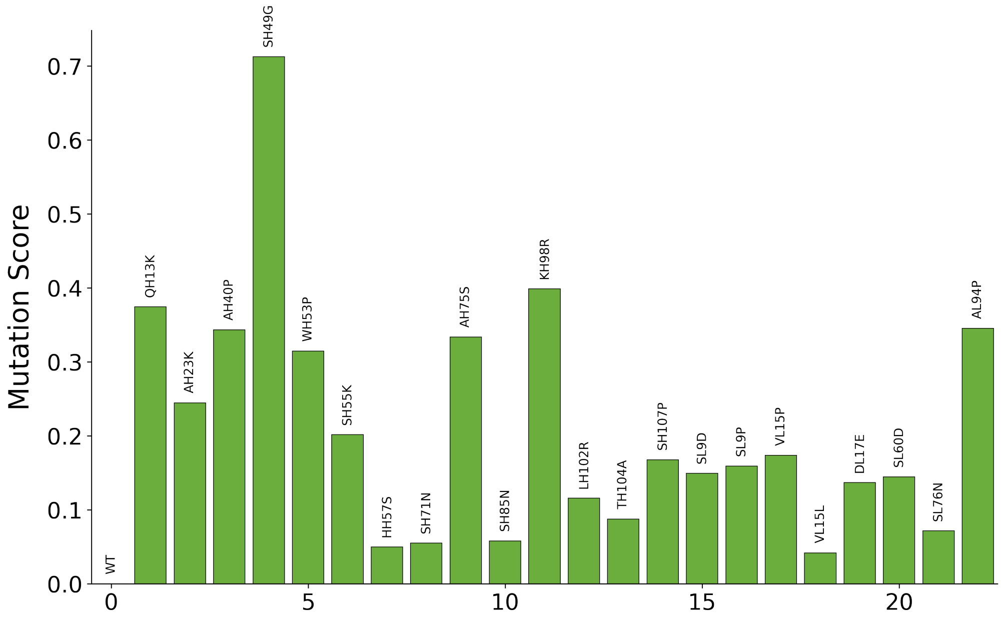


The plotted score is a rather arbitrary combination of delta solubility score, delta frequency, and predicted DDG (if FoldX was run). The highest this score the best the mutation. This score provides a nice visual ranking of mutations, but in practice one should refer to actual delta solubility score, delta frequency, and predicted DDG values to choose suitable mutations. This is for the single mutational scanning, then for combination of mutations, especially when across multiple chains, the mutation score is normalised usually resulting in a decreased contribution of the PSSM frequency (see paper).
