## Supplementary files 1 to 9 for "Automated optimisation of solubility and conformational stability of antibodies and proteins": SF7 CR3022_PDB7jn5_postPhase1MSA_5muts report.pdf

**CamSol analysis of input pdb file: 7jn5.pdb**

The following sequence positions are excluded from the design (but their presence is considered in solubility calculations - these may be excluded by the user through the 'Residues that can't be changed' field or becase of e.g. they have missing PSSM information):
Chain F: all
Chain H: R120, R121, L122, P123, P124, S125, V126, F127, P128, L129, A130, P131, S132, S133, K134, S135, T136, S137, G138, G139, T140, A141, A142, L143, G144, C145, L146, V147, K148, D149, Y150, F151, P152, E153, P154, V155, T156, V157, S158, W159, N160, S161, G162, A163, L164, T165, S166, G167, V168, H169, T170, F171, P172, A173, V174, L175, Q176, S177, S178, G179, L180, Y181, S182, L183, S184, S185, V186, V187, T188, V189, P190, S191, S192, S193, L194, G195, T196, Q197, T198, Y199, I200, C201, N202, V203, N204, H205, K206, P207, S208, N209, T210, K211, V212, D213, K214, K215, V216, E217, P218, K219, S220, C221
Chain L: T115, V116, A117, A118, P119, S120, V121, F122, I123, F124, P125, P126, S127, D128, E129, Q130, L131, K132, S133, G134, T135, A136, S137, V138, V139, C140, L141, L142, N143, N144, F145, Y146, P147, R148, E149, A150, K151, V152, Q153, W154, K155, V156, D157, N158, A159, L160, Q161, S162, G163, N164, S165, Q166, E167, S168, V169, T170, E171, Q172, D173, S174, K175, D176, S177, T178, Y179, S180, L181, S182, S183, T184, L185, T186, L187, S188, K189, A190, D191, Y192, E193, K194, H195, K196, V197, Y198, A199, C200, E201, V202, T203, H204, Q205, G206, L207, S208, S209, P210, V211, T212, K213, S214, F215, N216, R217, G218, E219, C220, S221

These amino acids are excluded from the list of potential substitution targets: C, M, N

**Sequences extracted from input pdb and used for analysis:**
lower case residues (if any) are residues of missing coordinates but present in the SEQRES field of the input pdb file. Such residues are considered for the solubility calculation, but are never targeted for mutations and their potential impact on stability is not considered.

```
> 7jn5:F
rvvpsgdvvrfpnitnlCPFGEVFNATKFPSVYAWERKKISNCVADYSVLYNSTFFSTFKCYGVSATKLNDLCFSNVYADSFVVKGDDVRQIAPGQTGVIADYNYKLPDDFMGCVLAWNT
RNidatstgnhnYKYRYLRHGKLRPFERDISNVPFSPDGKPCTPPALNCYWPLNDYGFYTTTGIGYQPYRVVVLSFEllNAPATVCGPklstdliknqcvnfsghhhhhh

> 7jn5:H
QMQLVQSGTEVKKPGESLKISCKGSGYGFITYWIGWVRQMPGKGLEWMGIIYPGDSETRYSPSFQGQVTISADKSINTAYLQWSSLKASDTAIYYCAGGSGISTPMDVWGQGTTVTVVSR
RLPPSVFPLAPSSKSTSGGTAALGCLVKDYFPEPVTVSWNSGALTSGVHTFPAVLQSSGLYSLSSVVTVPSSSLGTQTYICNVNHKPSNTKVDKKVEPKSC

> 7jn5:L
DIQLTQSPDSLAVSLGERATINCKSSQSVLYSSINKNYLAWYQQKPGQPPKLLIYWASTRESGVPDRFSGSGSGTDFTLTISSLQAEDVAVYYCQQYYSTPYTFGQGTKVEIKRTVAAPS
VFIFPPSDEQLKSGTASVVCLLNNFYPREAKVQWKVDNALQSGNSQESVTEQDSKDSTYSLSSTLTLSKADYEKHKVYACEVTHQGLSSPVTKSFNRGECs
```

Using log-likelihood pssm. Considering only candidate mutations with positive enrichment (log-likelihood > 0), and further restricting the space of candidate substitutions at each position to those residues that are more likely than the WT one

#### PSSM used to pick candidate mutations

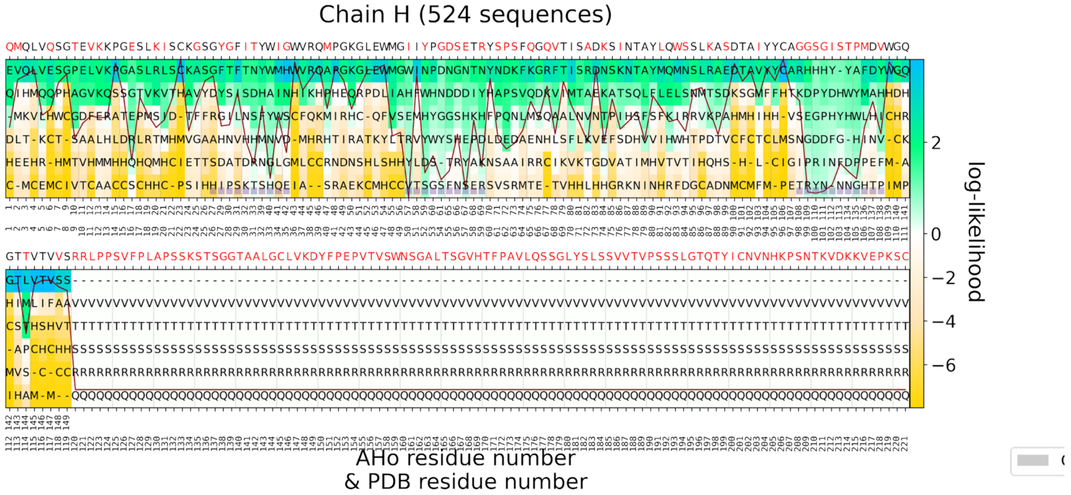


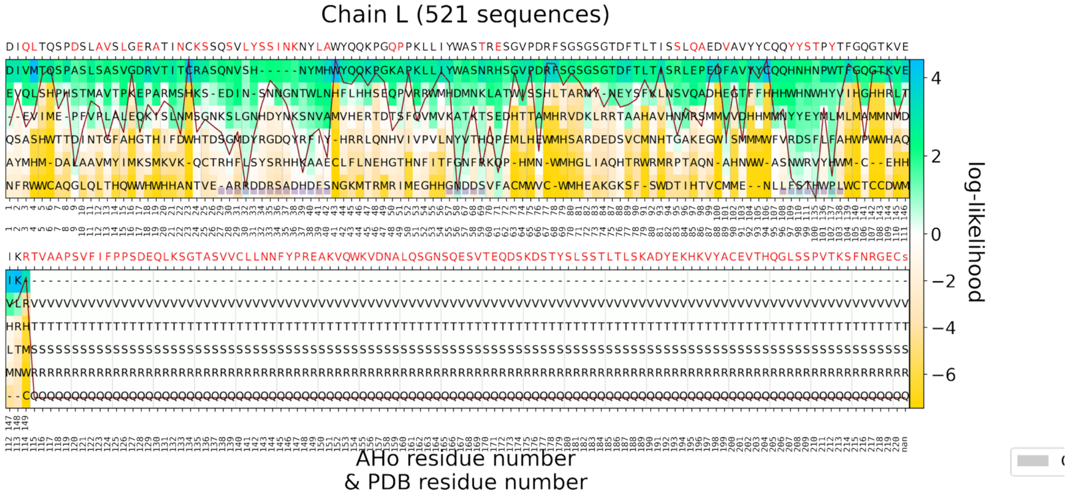


Position-specific scoring matrix (PSSM), as calculated from a multiple-sequence alignment (MSA) of similar sequences. The observed residue frequency (color-bar) is used to select candidate amino acid substitutions. The sequence above the panels is the wild-type (input) sequence as read from the alignment. The red line (if present) is the conservation index of each position (high means position highly conserved). Starting from the top panel: PSSM obtained from MSA of chain H containing 524 Fv sequences (Fv region only). PSSM obtained from MSA of chain L containing 521 Fv sequences (Fv region only).

#### CamSol intrinsic and structurally corrected profiles

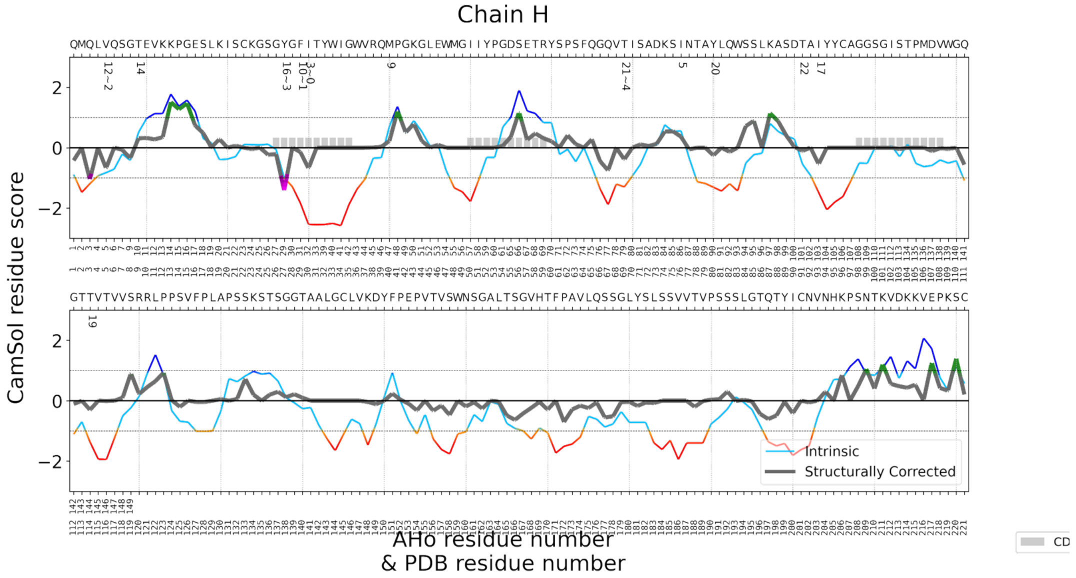

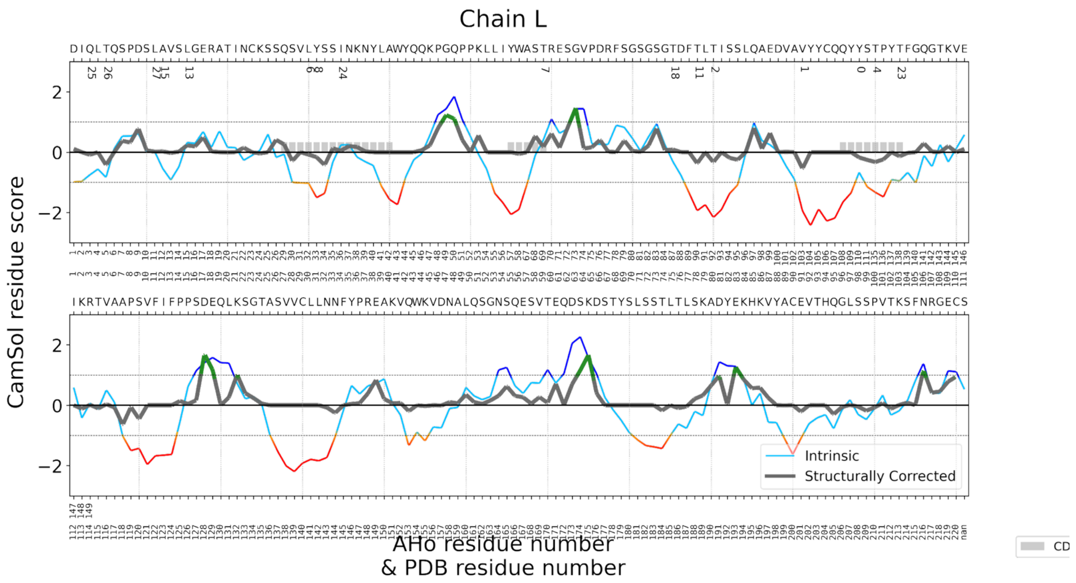


For each chain (see titles of the panels) the CamSol intrinsic profile is colour-coded red to blue, where red means aggregation-prone and blue aggregation-resistant. It is common for folded proteins to have large aggregation-prone regions in their intrinsic profile that typically drive the hydrophobic collapse during folding. The structurally corrected profile is color-coded in gray/green/magenta, regions of low negative scores (magenta) are potential aggregation hotspots, regions of high score (green) are solubility promoting. Numbers below the amino acid sequence at the top denote potential mutation sites, identified according to their contribution to the solubility, as well as their accessibility to the solvent.

### DESIGN PIPELINE RESULTS: Best Models

#### Table with identified best combinations of mutations

| Design Name | Number of Mutations | Mutations in Combination | Stability Rank | Solubility Rank | Theoretical PI | Mutation Score |
|---|---|---|---|---|---|---|
| model_7 | 2 | MH40P,QH67R | 3 | 3 | 8.643 | 0.478 |
| model_6 | 2 | MH40P,TL59K | 2 | 4 | 8.641 | 0.477 |
| model_8 | 2 | MH40P,SH56G | 2 | 5 | 8.581 | 0.476 |
| model_4 | 1 | MH40P | 3 | 5 | 8.581 | 0.315 |
| model_1 | 1 | QH67R | 5 | 4 | 8.643 | 0.163 |
| model_10 | 1 | TL59K | 5 | 4 | 8.641 | 0.162 |
| model_9 | 5 | MH40P,SH56G,QH67R,IH76K,TL59K | 1 | 1 | 8.748 | 0.958 |
| model_11 | 5 | MH40P,SH56G,QH67R,SH89E,TL59K | 1 | 2 | 8.641 | 0.931 |
| model_5 | 5 | MH40P,QH67R,IH76K,SH89E,TL59K | 1 | 1 | 8.695 | 0.926 |
| model_2 | 4 | MH40P,SH56G,QH67R,TL59K | 1 | 2 | 8.698 | 0.801 |
| model_3 | 3 | MH40P,QH67R,TL59K | 2 | 2 | 8.698 | 0.64 |
| WT | 0 | WT | 5 | 5 | 8.581 | 0 |

This table contains information on the identified mutation combinations, selected on the basis of their contribution to the overall solubility and stability of the protein. The column 'design name' identifies the pdb file bearing the mutations listed in the column 'mutations in combination'. Combinations belonging to the best combination groups (i.e., combinations containing 1, 2 mutants) are listed first and sorted according to their mutation score. The remaining combinations, not identified as best mutation combinations, are listed afterwards and are sorted according to their mutation score. Mutation combinations are given a solubility and stability rank according to their delta CamSol score and DDG value respectively (i.e. they are binned starting from 1 corresponding the best predicted). The top ranking combination is usually one with the highest 'Number of Mutations'. However, because with each added mutation the chances of introducing a false positive increase, the best combination groups (top rows in the table) embody the best tradeoff between low number of mutations and high gain in solubility and stability. All combinations listed are expected to increase both solubility and stability (or one without compromising the other) and contain single point mutants that are enriched in the input PSSM.

#### 2 Simultaneous Mutations:

**Model Name: Model_7**

Mutations by chain

Chain H : M40P,Q67R

Solubility Ranking: 3 | Delta CamSol score: 0.087

Stability Ranking: 3 | FoldX DDG: -3.157 kcal/mol

Mutation Score: 0.478

```
> Model_7 chain F Optimized_7jn5_Repair
CPFGEVFNATKFPSVYAWERKKISNCVADYSVLYNSTFFSTFKCYGVSATKLNDLCFSNVYADSFVVKGDDVRQIAPGQTGVIADYNYKLPDDFMGCVLAWNTRN----------YKYRYL
RHGKLRPFERDISNVPFSPDGKPCTPPALNCYWPLNDYGFYTTTGIGYQPYRVVVLSFE--NAPATVCGP
> Model_7 chain H Optimized_7jn5_Repair | M40P Q67R
QMQLVQSGTEVKKPGESLKISCKGSGYGFITYWIGWVRQPPGKGLEWMGIIYPGDSETRYSPSFQGRVTISADKSINTAYLQWSSLKASDTAIYYCAGGSGISTPMDVWGQGTTVTVVSRR
LPPSVFPLAPSSKSTSGGTAALGCLVKDYFPEPVTVSWNSGALTSGVHTFPAVLQSSGLYSLSSVVTVPSSSLGTQTYICNVNHKPSNTKVDKKVEPKSC
> Model_7 chain L Optimized_7jn5_Repair
DIQLTQSPDSLAVSLGERATINCKSSQSVLYSSINKNYLAWYQQKPGQPPKLLIYWASTRESGVPDRFSGSGSGTDFTLTISSLQAEDVAVYYCQQYYSTPYTFGQGTKVEIKRTVAAPSV
FIFPPSDEQLKSGTASVVCLLNNFYPREAKVQWKVDNALQSGNSQESVTEQDSKDSTYSLSSTLTLSKADYEKHKVYACEVTHQGLSSPVTKSFNRGEC
```

**Model Name: Model_6**

Mutations by chain

Chain H : M40P

Chain L : T59K

Solubility Ranking: 4 | Delta CamSol score: 0.074

Stability Ranking: 2 | FoldX DDG: -3.815 kcal/mol

Mutation Score: 0.477

```
> Model_6 chain F Optimized_7jn5_Repair
CPFGEVFNATKFPSVYAWERKKISNCVADYSVLYNSTFFSTFKCYGVSATKLNDLCFSNVYADSFVVKGDDVRQIAPGQTGVIADYNYKLPDDFMGCVLAWNTRN----------YKYRYL
RHGKLRPFERDISNVPFSPDGKPCTPPALNCYWPLNDYGFYTTTGIGYQPYRVVVLSFE--NAPATVCGP
> Model_6 chain H Optimized_7jn5_Repair | M40P
QMQLVQSGTEVKKPGESLKISCKGSGYGFITYWIGWVRQPPGKGLEWMGIIYPGDSETRYSPSFQGQVTISADKSINTAYLQWSSLKASDTAIYYCAGGSGISTPMDVWGQGTTVTVVSRR
LPPSVFPLAPSSKSTSGGTAALGCLVKDYFPEPVTVSWNSGALTSGVHTFPAVLQSSGLYSLSSVVTVPSSSLGTQTYICNVNHKPSNTKVDKKVEPKSC
> Model_6 chain L Optimized_7jn5_Repair | T59K
DIQLTQSPDSLAVSLGERATINCKSSQSVLYSSINKNYLAWYQQKPGQPPKLLIYWASKRESGVPDRFSGSGSGTDFTLTISSLQAEDVAVYYCQQYYSTPYTFGQGTKVEIKRTVAAPSV
FIFPPSDEQLKSGTASVVCLLNNFYPREAKVQWKVDNALQSGNSQESVTEQDSKDSTYSLSSTLTLSKADYEKHKVYACEVTHQGLSSPVTKSFNRGEC
```

**Model Name: Model_8**

Mutations by chain

Chain H : M40P,S56G

Solubility Ranking: 5 | Delta CamSol score: 0.028

Stability Ranking: 2 | FoldX DDG: -4.067 kcal/mol

Mutation Score: 0.476

```
> Model_8 chain F Optimized_7jn5_Repair
CPFGEVFNATKFPSVYAWERKKISNCVADYSVLYNSTFFSTFKCYGVSATKLNDLCFSNVYADSFVVKGDDVRQIAPGQTGVIADYNYKLPDDFMGCVLAWNTRN----------YKYRYL
RHGKLRPFERDISNVPFSPDGKPCTPPALNCYWPLNDYGFYTTTGIGYQPYRVVVLSFE--NAPATVCGP
> Model_8 chain H Optimized_7jn5_Repair | M40P S56G
QMQLVQSGTEVKKPGESLKISCKGSGYGFITYWIGWVRQPPGKGLEWMGIIYPGDGETRYSPSFQGQVTISADKSINTAYLQWSSLKASDTAIYYCAGGSGISTPMDVWGQGTTVTVVSRR
LPPSVFPLAPSSKSTSGGTAALGCLVKDYFPEPVTVSWNSGALTSGVHTFPAVLQSSGLYSLSSVVTVPSSSLGTQTYICNVNHKPSNTKVDKKVEPKSC
> Model_8 chain L Optimized_7jn5_Repair
DIQLTQSPDSLAVSLGERATINCKSSQSVLYSSINKNYLAWYQQKPGQPPKLLIYWASTRESGVPDRFSGSGSGTDFTLTISSLQAEDVAVYYCQQYYSTPYTFGQGTKVEIKRTVAAPSV
FIFPPSDEQLKSGTASVVCLLNNFYPREAKVQWKVDNALQSGNSQESVTEQDSKDSTYSLSSTLTLSKADYEKHKVYACEVTHQGLSSPVTKSFNRGEC
```

#### 1 Single Mutation:

**Model Name: Model_4**

Mutations by chain

Chain H : M40P

Solubility Ranking: 5 | Delta CamSol score: 0.026

Stability Ranking: 3 | FoldX DDG: -2.715 kcal/mol

Mutation Score: 0.315

```
> Model_4 chain F Optimized_7jn5_Repair
CPFGEVFNATKFPSVYAWERKKISNCVADYSVLYNSTFFSTFKCYGVSATKLNDLCFSNVYADSFVVKGDDVRQIAPGQTGVIADYNYKLPDDFMGCVLAWNTRN----------YKYRYL
RHGKLRPFERDISNVPFSPDGKPCTPPALNCYWPLNDYGFYTTTGIGYQPYRVVVLSFE--NAPATVCGP
> Model_4 chain H Optimized_7jn5_Repair | M40P
QMQLVQSGTEVKKPGESLKISCKGSGYGFITYWIGWVRQPPGKGLEWMGIIYPGDSETRYSPSFQGQVTISADKSINTAYLQWSSLKASDTAIYYCAGGSGISTPMDVWGQGTTVTVVSRR
LPPSVFPLAPSSKSTSGGTAALGCLVKDYFPEPVTVSWNSGALTSGVHTFPAVLQSSGLYSLSSVVTVPSSSLGTQTYICNVNHKPSNTKVDKKVEPKSC
> Model_4 chain L Optimized_7jn5_Repair
DIQLTQSPDSLAVSLGERATINCKSSQSVLYSSINKNYLAWYQQKPGQPPKLLIYWASTRESGVPDRFSGSGSGTDFTLTISSLQAEDVAVYYCQQYYSTPYTFGQGTKVEIKRTVAAPSV
FIFPPSDEQLKSGTASVVCLLNNFYPREAKVQWKVDNALQSGNSQESVTEQDSKDSTYSLSSTLTLSKADYEKHKVYACEVTHQGLSSPVTKSFNRGEC
```

**Model Name: Model_1**

Mutations by chain

Chain H : Q67R

Solubility Ranking: 4 | Delta CamSol score: 0.061

Stability Ranking: 5 | FoldX DDG: -0.443 kcal/mol

Mutation Score: 0.163

```
> Model_1 chain F Optimized_7jn5_Repair
CPFGEVFNATKFPSVYAWERKKISNCVADYSVLYNSTFFSTFKCYGVSATKLNDLCFSNVYADSFVVKGDDVRQIAPGQTGVIADYNYKLPDDFMGCVLAWNTRN----------YKYRYL
RHGKLRPFERDISNVPFSPDGKPCTPPALNCYWPLNDYGFYTTTGIGYQPYRVVVLSFE--NAPATVCGP
> Model_1 chain H Optimized_7jn5_Repair | Q67R
QMQLVQSGTEVKKPGESLKISCKGSGYGFITYWIGWVRQMPGKGLEWMGIIYPGDSETRYSPSFQGRVTISADKSINTAYLQWSSLKASDTAIYYCAGGSGISTPMDVWGQGTTVTVVSRR
LPPSVFPLAPSSKSTSGGTAALGCLVKDYFPEPVTVSWNSGALTSGVHTFPAVLQSSGLYSLSSVVTVPSSSLGTQTYICNVNHKPSNTKVDKKVEPKSC
> Model_1 chain L Optimized_7jn5_Repair
DIQLTQSPDSLAVSLGERATINCKSSQSVLYSSINKNYLAWYQQKPGQPPKLLIYWASTRESGVPDRFSGSGSGTDFTLTISSLQAEDVAVYYCQQYYSTPYTFGQGTKVEIKRTVAAPSV
FIFPPSDEQLKSGTASVVCLLNNFYPREAKVQWKVDNALQSGNSQESVTEQDSKDSTYSLSSTLTLSKADYEKHKVYACEVTHQGLSSPVTKSFNRGEC
```

**Model Name: Model_10**

Mutations by chain

Chain L : T59K

Solubility Ranking: 4 | Delta CamSol score: 0.048

Stability Ranking: 5 | FoldX DDG: -1.1 kcal/mol

Mutation Score: 0.162

```
> Model_10 chain F Optimized_7jn5_Repair
CPFGEVFNATKFPSVYAWERKKISNCVADYSVLYNSTFFSTFKCYGVSATKLNDLCFSNVYADSFVVKGDDVRQIAPGQTGVIADYNYKLPDDFMGCVLAWNTRN----------YKYRYL
RHGKLRPFERDISNVPFSPDGKPCTPPALNCYWPLNDYGFYTTTGIGYQPYRVVVLSFE--NAPATVCGP
> Model_10 chain H Optimized_7jn5_Repair
QMQLVQSGTEVKKPGESLKISCKGSGYGFITYWIGWVRQMPGKGLEWMGIIYPGDSETRYSPSFQGQVTISADKSINTAYLQWSSLKASDTAIYYCAGGSGISTPMDVWGQGTTVTVVSRR
LPPSVFPLAPSSKSTSGGTAALGCLVKDYFPEPVTVSWNSGALTSGVHTFPAVLQSSGLYSLSSVVTVPSSSLGTQTYICNVNHKPSNTKVDKKVEPKSC
> Model_10 chain L Optimized_7jn5_Repair | T59K
DIQLTQSPDSLAVSLGERATINCKSSQSVLYSSINKNYLAWYQQKPGQPPKLLIYWASKRESGVPDRFSGSGSGTDFTLTISSLQAEDVAVYYCQQYYSTPYTFGQGTKVEIKRTVAAPSV
FIFPPSDEQLKSGTASVVCLLNNFYPREAKVQWKVDNALQSGNSQESVTEQDSKDSTYSLSSTLTLSKADYEKHKVYACEVTHQGLSSPVTKSFNRGEC
```

**Best mutant combination groups identified for 1,2 simultaneous mutations in combination**

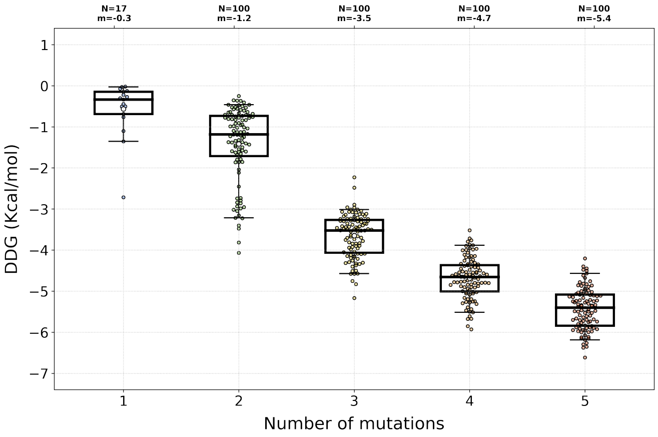


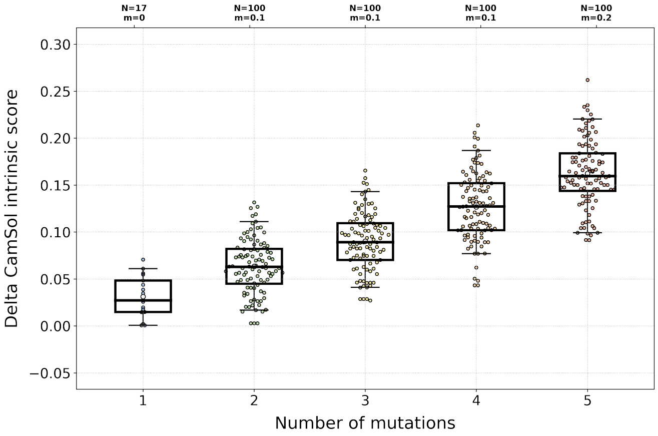


#### Table with identified candidate mutation sites (59 sites)

| Mutation site (seq index) | Mutation site (pdb number) | PSSM score > 0 | Delta PSSM score > 0 | site conservation | solvent exposure | solubilization potential | Problematic region+site score | region size | residue stru. corr. score | residue intrinsic score | wt residue frequency score | identified from |
|---|---|---|---|---|---|---|---|---|---|---|---|---|
| I29.H | IH30 | T S | S T | 0.505 | 0.253 | 0.083 | 9.066 | 11 | -0.687 | -2.541 | -1.048 | Solub. Seq. & Conservatio |
| F28.H | FH29 | I L | | 0.81 | 0.072 | 0.037 | 8.382 | 11 | -0.145 | -1.834 | 4.898 | Solub. Seq. |
| V4.H | VH5 | Q L | | 0.664 | 0.812 | 0.026 | 3.46 | 6 | -0.684 | -0.827 | 3.262 | Solub. Seq. |
| Y26.H | YH27 | F | F | 0.595 | 1.0 | 0.0 | 7.523 | 11 | -1.421 | -1.0 | 2.733 | Solub. Seq. |
| T68.H | TH69 | | | 0.875 | 0.598 | 0.0 | 4.452 | 5 | -0.393 | -1.3 | 3.714 | Solub. Seq. |
| Y97.L | YL98 | H D W S T | H | 0.245 | 0.251 | 0.116 | 6.024 | 8 | -0.174 | -0.668 | 1.877 | Solub. Seq. |
| V90.L | VL91 | T D | | 0.54 | 0.316 | 0.114 | 7.315 | 8 | -0.547 | -1.942 | 2.854 | Solub. Seq. |
| T79.L | TL80 | K | | 0.792 | 0.239 | 0.089 | 6.713 | 8 | -0.329 | -2.162 | 3.396 | Solub. Seq. |
| T99.L | TL100 | W Y F V H D L P | D Y H W L F V | 0.195 | 0.321 | 0.079 | 4.065 | 5 | -0.326 | -1.329 | 0.171 | Solub. Seq. |
| I75.H | IH76 | K T | K T | 0.575 | 0.637 | 0.056 | 2.736 | 6 | 0.363 | 0.556 | 1.264 | Solub. Seq. |
| | | S G D | | | | | | | | | | |

| | | | | | | | | | | | | |
|---|---|---|---|---|---|---|---|---|---|---|---|---|
| L29.L | LL30 | R V | G D S | 0.258 | 0.455 | 0.056 | 3.828 | 5 | -0.079 | -1.022 | 0.605 | Solub. Seq. |
| T58.L | TL59 | K H R S | K | 0.384 | 0.376 | 0.052 | 3.321 | 5 | 0.17 | 0.406 | 1.042 | Solub. Seq. |
| Y30.L | YL31 | H | H | 0.57 | 0.256 | 0.046 | 4.333 | 5 | -0.164 | -1.504 | 0.484 | Solub. Seq. |
| M39.H | MH40 | A P H R | A P | 0.705 | 0.361 | 0.042 | 5.776 | 11 | 0.224 | 0.714 | 0.647 | Solub. Seq. |
| T77.L | TL78 | S | | 0.781 | 0.359 | 0.03 | 6.487 | 8 | -0.371 | -1.938 | 3.393 | Solub. Seq. |
| L14.L | LL15 | V P | P V | 0.584 | 0.63 | 0.015 | 0.768 | 1 | 0.227 | 0.32 | 0.881 | Solub. Seq. |
| T8.H | TH9 | P A G | P A G | 0.515 | 0.63 | 0.015 | 2.068 | 6 | 0.307 | 0.503 | -4.176 | Solub. Seq. & Conservation |
| A11.L | AL12 | S P | S | 0.735 | 0.552 | 0.008 | 1.579 | 1 | 0.024 | -0.543 | 0.364 | Solub. Seq. |
| I92.H | IH93 | V | V | 0.777 | 0.411 | 0.0 | 5.3 | 6 | -0.548 | -1.487 | -0.316 | Solub. Seq. & Conservation |
| T74.L | TL75 | | | 0.822 | 0.492 | 0.0 | 4.796 | 8 | 0.086 | -0.314 | 3.448 | Solub. Seq. |
| T113.H | TH114 | L | L | 0.609 | 0.316 | 0.0 | 4.714 | 4 | -0.303 | -1.369 | 1.8 | Solub. Seq. |
| Y79.H | YH80 | F | | 0.785 | 0.269 | 0.0 | 4.602 | 6 | -0.258 | -1.311 | 3.777 | Solub. Seq. |
| T90.H | TH91 | | | 0.878 | 0.333 | 0.0 | 4.311 | 6 | -0.106 | -0.546 | 3.719 | Solub. Seq. |
| T102.L | TL103 | V | | 0.787 | 0.198 | 0.0 | 3.689 | 5 | -0.137 | -0.953 | 3.386 | Solub. Seq. |
| I33.L | IL34 | | | 0.838 | 0.238 | 0.0 | 2.589 | 5 | 0.049 | 0.24 | -3.517 | Solub. Seq. & Conservation |
| Q2.L | QL3 | V | V | 0.616 | 0.73 | 0.0 | 2.475 | 3 | -0.083 | -0.751 | 2.8 | Solub. Seq. |
| T4.L | TL5 | | | 0.924 | 0.838 | 0.0 | 1.8 | 1 | -0.429 | -0.835 | 3.612 | Solub. Seq. |
| L10.L | LL11 | V | | 0.771 | 0.153 | 0.0 | 1.058 | 1 | 0.016 | 0.027 | 3.607 | Solub. Seq. |
| V117.H | VH118 | S | S | 0.903 | 0.086 | 0.011 | | | 0.104 | -0.5 | -5.171 | Conservation |
| G27.H | GH28 | T S I | T S I | 0.602 | 0.0 | 0.0 | | | -0.0 | -1.29 | -5.573 | Conservation |
| R113.L | RL114 | | | 0.97 | 0.375 | 0.0 | | | 0.014 | 0.069 | -4.024 | Conservation |
| G97.H | GH98 | R K | R K | 0.755 | 0.0 | 0.03 | | | -0.0 | -0.511 | -3.307 | Conservation |
| S32.L | SL33 | | | 0.893 | 1.0 | 0.0 | | | 0.126 | -0.329 | -2.755 | Conservation |
| S88.H | SH89 | E D A | E D A | 0.745 | 0.885 | 0.02 | | | 0.402 | 0.386 | -1.724 | Conservation |
| G23.H | GH24 | A V | A V | 0.743 | 0.0 | 0.0 | | | -0.0 | 0.093 | -1.318 | Conservation |
| M1.H | MH2 | V | V | 0.888 | 0.0 | 0.0 | | | -0.0 | -1.481 | -0.929 | Conservation |
| S60.H | SH61 | A P | A P | 0.515 | 0.034 | 0.004 | | | 0.007 | -0.224 | -1.006 | Conservation |
| T103.H | TH104 | Y W H D F G | Y H W D G F | 0.308 | 0.0 | 0.003 | | | 0.0 | 0.122 | -1.399 | Conservation |
| Q66.H | QH67 | R K | R K | 0.845 | 0.913 | 0.061 | | | -0.757 | -1.895 | -0.307 | Conservation |
| N34.L | NL35 | G | G | 0.751 | 0.965 | 0.0 | | | 0.202 | 0.25 | -0.403 | Conservation |
| S31.L | SL32 | | | 0.754 | 1.0 | 0.0 | | | -0.427 | -1.362 | -0.017 | Conservation |
| G34.H | GH35 | H S | H S | 0.428 | 0.0 | 0.0 | | | -0.0 | -1.883 | -0.554 | Conservation |
| S102.H | SH103 | H Y G F | H Y G F | 0.361 | 0.109 | 0.001 | | | -0.045 | -0.301 | -0.322 | Conservation |
| S98.L | SL99 | H E R T | E H | 0.296 | 0.197 | 0.071 | | | -0.257 | -1.149 | 1.524 | Exposed Solub. |
| Q81.H | QH82 | E K H | | 0.622 | 0.188 | 0.063 | | | -0.14 | -1.192 | 3.651 | Exposed Solub. |
| Q0.H | QH1 | E | E | 0.588 | 0.617 | 0.047 | | | -0.423 | -0.891 | 3.17 | Exposed Solub. |

| | | | | | | | | | | | | |
|---|---|---|---|---|---|---|---|---|---|---|---|---|
| S82.L | SL83 | R G | R | 0.567 | 0.498 | 0.044 | | | -0.253 | -1.085 | 2.309 | Exposed Solub. |
| S27.L | SL28 | D G H | D | 0.452 | 0.407 | 0.039 | | | -0.29 | -0.998 | 1.944 | Exposed Solub. |
| Q84.L | QL85 | E | E | 0.693 | 0.41 | 0.035 | | | 0.813 | 0.976 | 3.32 | Exposed Solub. |
| Q64.H | QH65 | K | K | 0.75 | 0.566 | 0.035 | | | 0.28 | -0.643 | 1.982 | Exposed Solub. |
| A71.H | AH72 | R V K | R | 0.492 | 0.148 | 0.035 | | | 0.053 | 0.063 | 1.673 | Exposed Solub. |
| S55.H | SH56 | G D | G D | 0.543 | 0.428 | 0.028 | | | 1.184 | 1.923 | 0.74 | Exposed Solub. |
| Q47.L | QL48 | K | K | 0.595 | 0.445 | 0.027 | | | 1.106 | 1.855 | 2.813 | Exposed Solub. |
| S62.H | SH63 | K | K | 0.486 | 0.661 | 0.015 | | | 0.117 | -0.467 | 2.116 | Exposed Solub. |
| V10.H | VH11 | L | L | 0.659 | 0.172 | 0.011 | | | 0.274 | 1.128 | 2.473 | Exposed Solub. |
| R58.H | RH59 | Y H D K I | H Y D K | 0.339 | 0.272 | 0.009 | | | 0.315 | 0.833 | 0.421 | Exposed Solub. |
| P61.H | PH62 | D Q E | D | 0.451 | 0.646 | 0.009 | | | 0.408 | -0.053 | 2.539 | Exposed Solub. |
| K18.H | KH19 | R S | R | 0.586 | 0.464 | 0.001 | | | 0.281 | -0.396 | 3.42 | Exposed Solub. |
| K86.H | KH87 | R T | R T | 0.694 | 0.497 | 0.001 | | | 1.139 | 0.801 | 1.21 | Exposed Solub. |

##### Results of the single-mutation scanning at all suitable sites

| mut_id_seqIndex | mut_id_pdb | Mutation Score | Delta CamSol intrinsic score | delta_frequency | DDG (kcal/mol) | Mutation type | CamSol intrinsic score | Mutation frequency | Chain F CamSol intrinsic score | Chain H CamSol intrinsic score | [illegible] |
|---|---|---|---|---|---|---|---|---|---|---|---|
| WT | WT | 0 | 0.0 | 0.0 | | n/a | -1.891 | n/a | -0.317 | -0.612 | [illegible] |
| TH8P | TH9P | 0.447 | 0.015 | 6.839 | -0.223 | Solub. Seq. | -1.876 | 2.663 | -0.317 | -0.586 | [illegible] |
| KH18R | KH19R | 0.03 | 0.001 | 0.086 | -0.238 | Exposed Solub. | -1.89 | 3.506 | -0.317 | -0.611 | [illegible] |
| MH39P | MH40P | 0.384 | 0.026 | 1.44 | -2.715 | Solub. Seq. | -1.865 | 2.087 | -0.317 | -0.566 | [illegible] |
| SH55G | SH56G | 0.255 | 0.002 | 1.957 | -1.353 | Exposed Solub. | -1.888 | 2.696 | -0.317 | -0.608 | [illegible] |
| SH62K | SH63K | 0.077 | 0.015 | 1.004 | -0.018 | Exposed Solub. | -1.876 | 3.12 | -0.317 | -0.585 | [illegible] |
| QH64K | QH65K | 0.201 | 0.035 | 2.311 | -0.272 | Exposed Solub. | -1.856 | 4.292 | -0.317 | -0.55 | [illegible] |
| QH66R | QH67R | 0.389 | 0.061 | 4.722 | -0.443 | Conservation | -1.83 | 4.415 | -0.317 | -0.503 | [illegible] |
| QH66K | QH67K | 0.215 | 0.055 | 1.825 | -0.507 | Conservation | -1.836 | 1.519 | -0.317 | -0.514 | [illegible] |

| | | | | | | | | | | | |
|---|---|---|---|---|---|---|---|---|---|---|---|
| IH75K | IH76K | 0.282 | 0.056 | 2.625 | -0.684 | Solub. Seq. | -1.834 | 3.889 | -0.317 | -0.512 | - |
| KH86R | KH87R | 0.225 | 0.001 | 2.909 | -0.502 | Exposed Solub. | -1.89 | 4.119 | -0.317 | -0.611 | - |
| SH88E | SH89E | 0.43 | 0.02 | 6.289 | -0.333 | Conservation | -1.871 | 4.565 | -0.317 | -0.577 | - |
| SH88D | SH89D | 0.214 | 0.018 | 3.034 | -0.144 | Conservation | -1.873 | 1.311 | -0.317 | -0.58 | - |
| SH102G | SH103G | 0.115 | 0.001 | 0.634 | -0.764 | Conservation | -1.89 | 0.312 | -0.317 | -0.611 | - |
| SL27D | SL28D | 0.082 | 0.039 | 0.508 | -0.127 | Exposed Solub. | -1.852 | 2.453 | -0.317 | -0.612 | - |
| QL47K | QL48K | 0.08 | 0.027 | 0.751 | -0.075 | Exposed Solub. | -1.863 | 3.564 | -0.317 | -0.612 | - |
| TL58K | TL59K | 0.174 | 0.048 | 0.261 | -1.1 | Solub. Seq. | -1.842 | 1.303 | -0.317 | -0.612 | - |
| SL82R | SL83R | 0.089 | 0.044 | 0.256 | -0.301 | Exposed Solub. | -1.847 | 2.564 | -0.317 | -0.612 | - |
| SL98E | SL99E | 0.089 | 0.071 | 0.27 | -0.026 | Exposed Solub. | -1.82 | 1.794 | -0.317 | -0.612 | - |

This table contains information on all possible single mutations at sites identified on the basis of their contribution to the overall solubility and their solvent exposure, as well as to their conservation in the MSA (see previous table). The column 'Mutation type' describes how a site has been identified: Solub. Stru. denotes that it was a site contributing to poor local solubility in the structurally corrected profile; Solub. seq. is the same but for the sequence-based intrinsic profile; Conservation indicates that the wild-type amino acid had log-likelihood<0 in the PSSM; Exposed Solub. are additional solvent-exposed sites where the WT residue is not strongly conserved (other residues with high PSSM score exist at that position) that may be mutated to further increase solubility (albeit unlike Solub. Seq. and Solub. Stru. these are typically not close to or within candidate aggregation hotspots). The column 'mut_id_seqIndex' contains candidate mutations numbered according to the index of the mutation site along the input sequence (from index 0), while in that 'mut_id_pdb' mutations are numbered according to the residue number in the input pdb file. This table has been filtered to contain only point-mutations predicted to increase both predicted solubility and stability - and if the strict-PSSM cutoff is selected as input - also PSSM frequency (see file 7jn5single_mutation_scanning.csv for a full table with all mutations tested, including those predicted destabilising).

##### Result of single-mutation scanning at all suitable sites

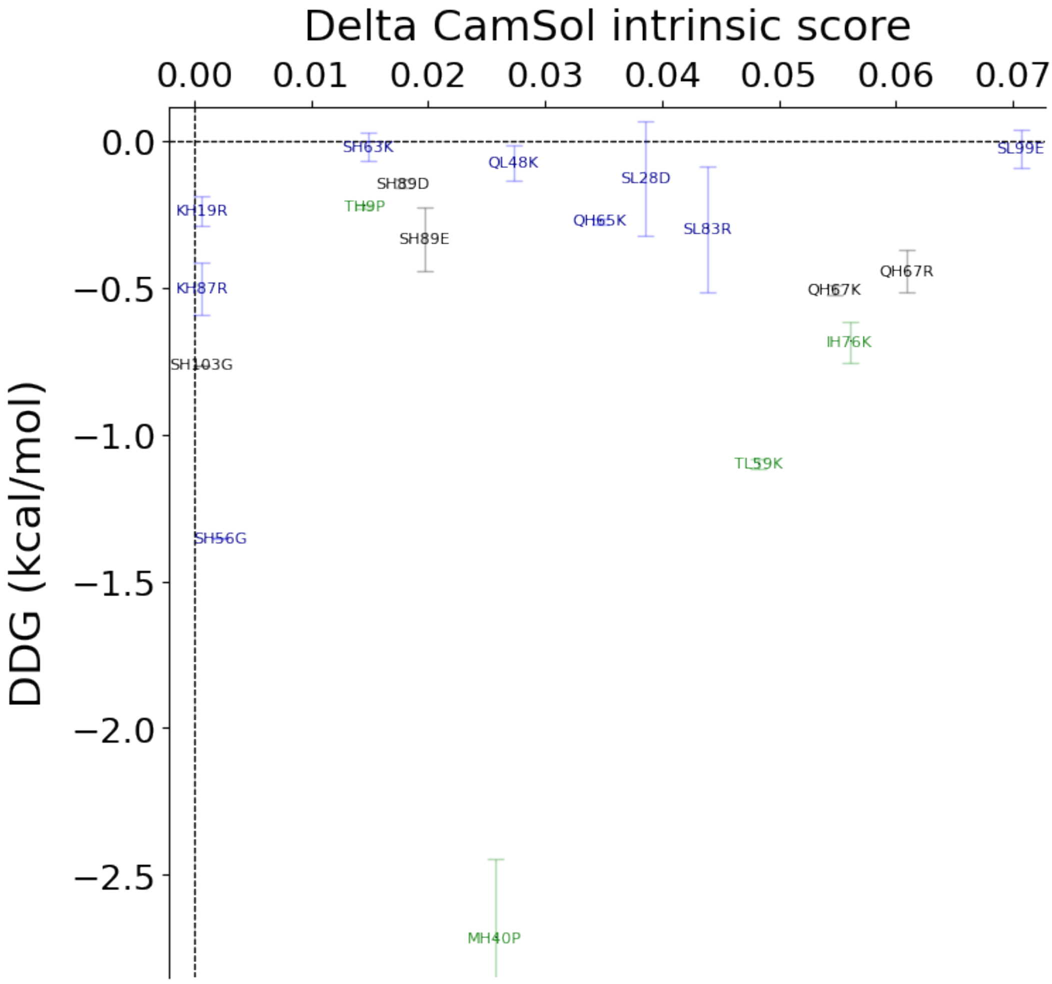

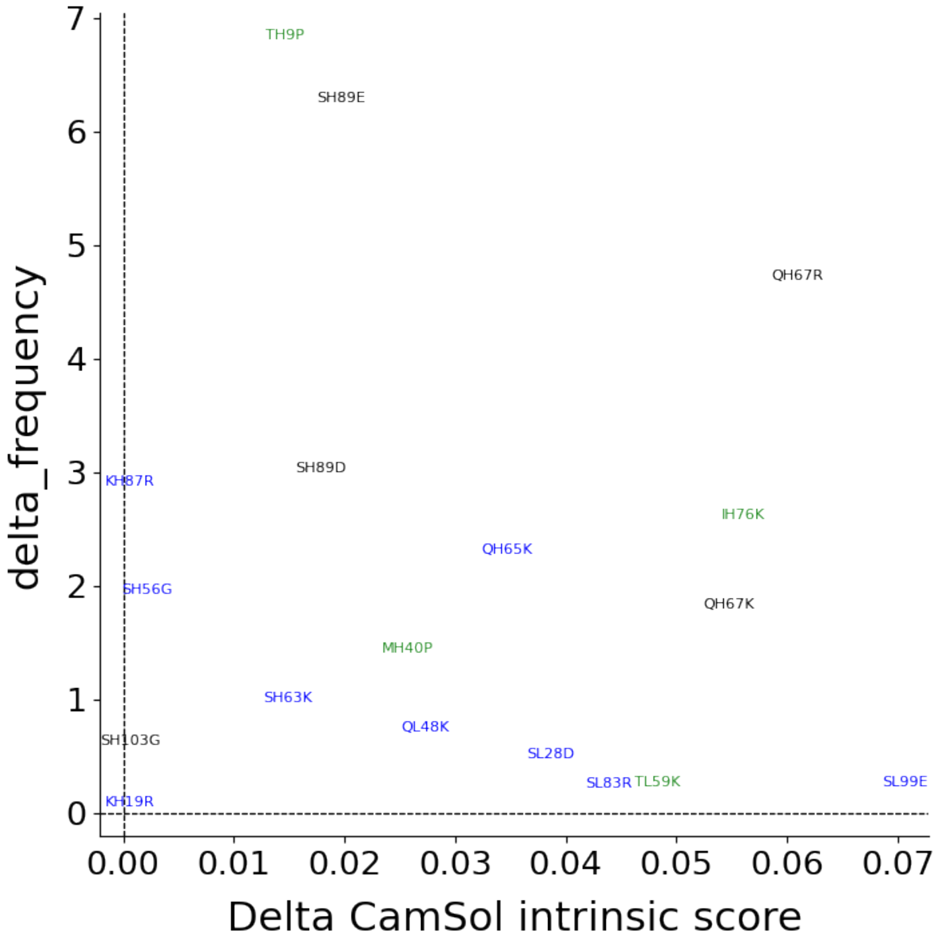


Plot of the data in the previous table, points are candidate mutations numbered according to the residue number within the input pdb file. The best mutations are those with large Delta CamSol score and large negative predicted DDG (if FoldX was run) and/or large positive Delta Frequency in MSA. Points in black correspond to mutation sites selected according to their conservation (see previous table), in green according to the sequence-based intrinsic solubility prediction, and in blue according to the structurally corrected one. Mutations at sites selected according to the solubility profiles (blue and green) are expected to have more impact on solubility than mutations at sites selected from conservation (black).

##### Mutation score of results of single-mutation scanning at all suitable sites (before normalisation)

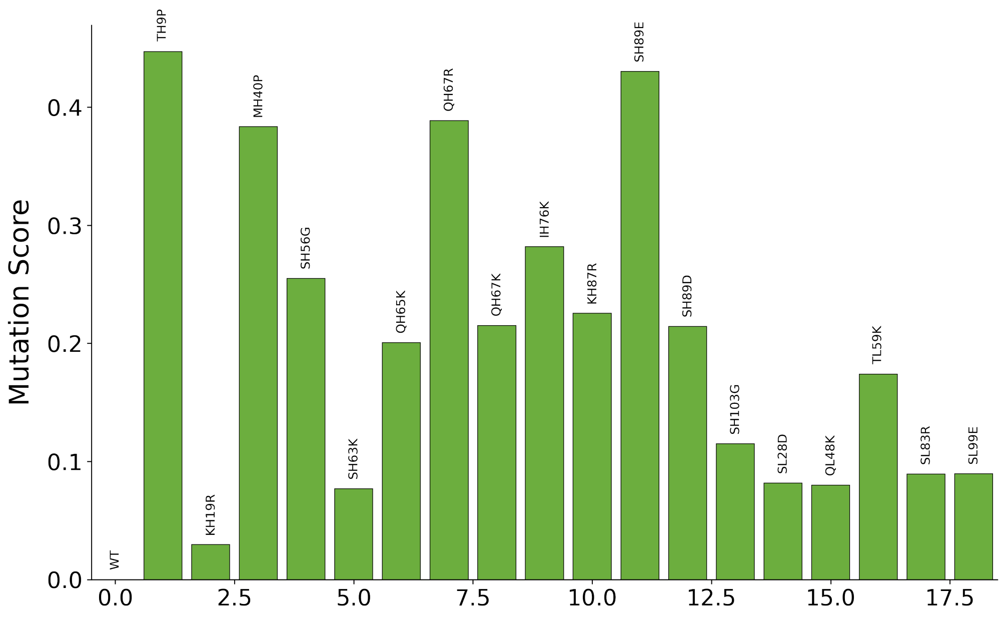


The plotted score is a rather arbitrary combination of delta solubility score, delta frequency, and predicted DDG (if FoldX was run). The highest this score the best the mutation. This score provides a nice visual ranking of mutations, but in practice one should refer to actual delta solubility score, delta frequency, and predicted DDG values to choose suitable mutations. This is for the single mutational scanning, then for combination of mutations, especially when across multiple chains, the mutation score is normalised usually resulting in a decreased contribution of the PSSM frequency (see paper).
